# One-to-one mapping between deep network units and real neurons uncovers a visual population code for social behavior

**DOI:** 10.1101/2022.07.18.500505

**Authors:** Benjamin R. Cowley, Adam J. Calhoun, Nivedita Rangarajan, Maxwell H. Turner, Jonathan W. Pillow, Mala Murthy

## Abstract

The rich variety of behaviors observed in animals arises through the complex interplay between sensory processing and motor control. To understand these sensorimotor transformations, it is useful to build models that predict not only neural responses to sensory input [1, 2, 3, 4, 5] but also how each neuron causally contributes to behavior [6, 7]. Here we demonstrate a novel modeling approach to identify a one-to-one mapping between internal units in a deep neural network and real neurons by predicting the behavioral changes arising from systematic perturbations of more than a dozen neuron types. A key ingredient we introduce is “knockout training”, which involves perturbing the network during training to match the perturbations of the real neurons during behavioral experiments. We apply this approach to model the sensorimotor transformation of *Drosophila melanogaster* males during a complex, visually-guided social behavior [8, 9, 10]. The visual projection neurons at the interface between the eye and brain form a set of discrete channels, suggesting each channel encodes a single visual feature [11, 12, 13]. Our model reaches a different conclusion: The visual projection neurons form a highly distributed population code that collectively sculpts social behavior. Overall, our framework consolidates behavioral effects elicited from various neural perturbations into a single, unified model, providing a detailed map from stimulus to neuron to behavior.

---

To understand how the brain performs a sensorimotor transformation, an emerging and popular approach is to first train a deep neural network (DNN) model on a behavioral task performed by an animal (e.g., recognize an object in an image) and then compare the animal’s neural activity to the internal activations of the DNN [1, 2, 3, 5, 14, 15, 16]. A shortcoming of this approach is that the DNN fails to predict how an individual neuron *causally* contributes to behavior, making it difficult to interpret a neuron’s role in the sensorimotor transformation. Here we overcome this drawback by perturbing the internal units of a DNN model while predicting the behavior of animals whose neurons have also been perturbed, a method we call ‘knockout training’. This approach places a strong constraint on the model: Each model unit must contribute to behavior in a way that matches the corresponding real neuron’s causal contribution to behavior. An added benefit is that the model infers neural activity from (perturbed) behavior alone. This is especially useful when studying social behaviors, as it is challenging (or impossible in some systems) to record neural activity during natural, social interactions. Here we use this approach to investigate the sensorimotor transformation of *Drosophila* males during natural courtship, including chasing and singing to a female [9].

## Training a deep neural network to model transformations from vision to behavior

The *Drosophila* visual system contains a bottleneck between the eyes and the central brain in the form of visual projection neurons with approximately 200 different cell types [17]. The primary cell types of this bottleneck (Fig. 1**a**) are the *∼*45 Lobula Columnar (LC) and Lobula Plate (LPLC) neuron types (we use ‘LC types’ to refer both to LC and LPLC neuron types) that receive input from the lobula and lobula plate in the optic lobe and send axons to a set of optic glomeruli that are read out by the central brain [11, 18]. Neurons of one and only one LC type innervate the same optic glomerulus [11, 19]. It has been suggested that each LC type acts as a “feature detector” to extract visual information and modulate sensory-driven behavior [12, 13]. Indeed, a large number of studies have each isolated a specific LC neuron type, identified its preferred visual feature, and uncovered its contribution to a specific behavior [10, 20, 21, 22, 23, 24, 25, 26, 27]. For example, LPLC2 neurons strongly respond to a looming object and innervate the giant fiber to drive an escape take off [24]. For the more complex behavior of courtship, LC10a neurons (and LC9 neurons, to a lesser extent) in males have been implicated in tracking the female’s position to influence the male’s turning behavior [10, 21, 22]. Whether other LC types, if any, contribute to courtship remains an open question.

**Figure 1:**
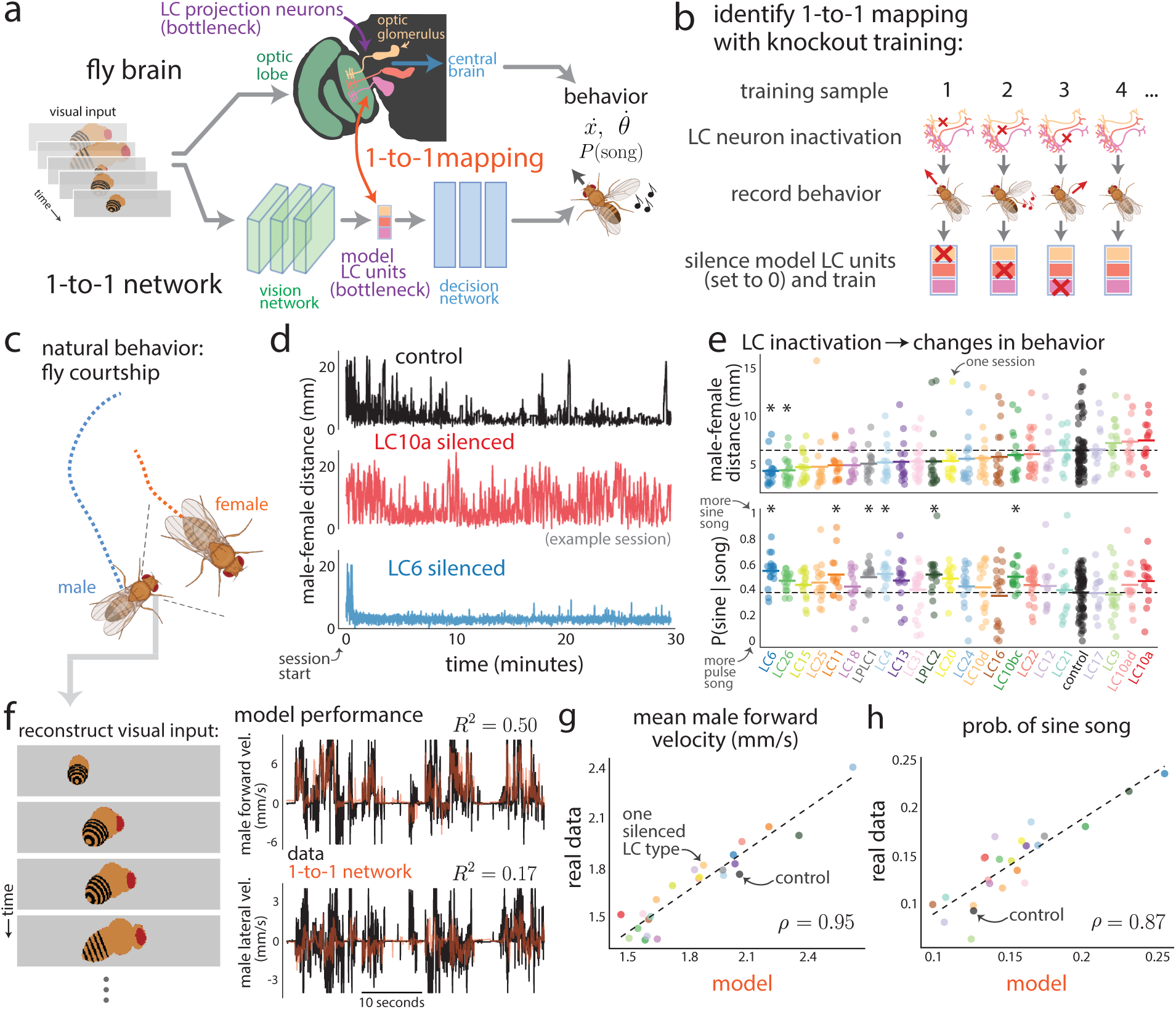
Identifying a one-to-one mapping between real neurons and internal units of a deep neural net-work with knockout training. **a**. Modeling the transformation from vision to behavior in male flies. The model (termed the 1-to-1 network) consists of a vision network with convolutional filters (green), a decision network with dense connections (blue), and a bottleneck of model LC units that matches the bottleneck of LC neuron types found in the fly’s visual system (purple). We seek a one-to-one mapping in which each model LC unit recovers the corresponding LC neuron type’s activity (i.e., the summed activity of LC neurons of the same type projecting to one optic glomerulus) as well as that LC neuron type’s causal contribution to behavior (via silencing). The model takes as input a sequence of images and outputs its prediction of the male’s movement, including forward, lateral, and angular velocity, as well the male’s song produced by wing vibration, including sine, pulse-fast (Pfast), and pulse-slow (Pslow) song. **b**. We designed a new procedure called knockout training to fit the 1-to-1 network. Data for each training sample are the visual inputs and male behaviors involving the bilateral genetic silencing of a particular LC neuron type (‘LC neuron inactivation’, red Xs). We silenced or “knocked out” the model LC unit (i.e., set its activity value to 0, red Xs) that corresponds to the silenced LC neuron type for that sample. Thus, we perturbed the model in a way similar to how the male fly was perturbed. **c**. We recorded female and male behavior during natural courtship; joint positions were extracted using pose tracking [28], and male song was extracted using audio segmentation [29]. **d**. Courtship behavior noticeably changed between control and LC-silenced male flies (using a set of sparse, specific genetic driver lines from [11]); example sessions. Control flies (top row) began each session far apart (i.e., at larger male-female distances) and eventually decreased their distance. In contrast, male flies with silenced LC10a neurons (middle row) maintained a large distance from the female, consistent with previous studies [10, 22]. Male flies with silenced LC6 neurons (bottom row) had small distances throughout the entire session, indicating these flies strongly pursued the female from the beginning of the session. **e**. Changes in the average male-to-female distance following silencing of each LC type in males (top row, average taken over entire session) and changes in the proportion of song that was sine versus pulse (bottom row, taken over entire session)—sine is typically sung at close distances to the female, whereas pulse is produced at larger distances and faster male speeds [9]. Each dot denotes one courtship session (one male-female pair). Short lines denote means across sessions for each LC type; horizontal dashed line denotes mean of control sessions. As-terisks denote signficant deviation from control; *p <* 0.05, permutation test corrected for multiple comparisons. **f**. The 1-to-1 network takes as input a sequence of 10 images (left inset, grayscale 64*×*256 pixel images—color images are shown for clarity) corresponding to the 10 most recent time frames (*∼*300 ms) of the male’s visual experience. Further increasing the number of time frames did not improve prediction. Each image is a reconstruction of what the male fly observed based on the male and female joint positions of that time frame (e.g., **c**). Based on visual inputs alone, the 1-to-1 network is able to predict the male fly’s behavior, including forward velocity (right, top row) and lateral velocity (right, bottom row). Reported *R*^2^’s are computed from held-out control behavior and not from these example traces. **g** and **h**. The 1-to-1 network also accurately predicts overall mean changes in behavior across males with different silenced LC neuron types. Correlations *ρ* were significant (*p <* 0.002, permutation test).

There is emerging evidence that these LC types may not work independently but rather as a distributed population. For example, multiple LC types have overlapping responses to similar stimuli [30, 31, 32], and optogenetically ac-tivating a single LC type can lead to different behaviors [11]. Even the striking anatomy of the LC neuron types— bundled into separate “channels” that make up the optic glomeruli—may be misleading, as olfactory glomeruli, whose anatomy closely resembles that of the optic glomeruli (e.g., each olfactory glomerulus receives input from one olfactory receptor neuron type) [33, 34], are read out in a more random way by downstream neurons in the mushroom bodies [35, 36], a hallmark of a distributed population code [37, 38]. Still, that the LC types form such a code is puzzling. For example, if LPLC2 responds to other visual features besides a looming object, what prevents LPLC2 from driving the giant fiber when responding to these other features? The extent to which these LC types form a distribution population code, especially for complex behaviors, remains unclear.

To investigate the LC code, we designed a novel DNN modeling approach for identifying the functional roles of individual LC neuron types using behavioral data from genetically altered flies (Fig. 1**a**, bottom diagram). Our approach relies on a DNN model with three components: (1) a front-end convolutional vision network that reflects processing in the optic lobe; (2) a “bottleneck” layer of LC units where each model LC unit represents the summed activity of neurons of the same LC type (i.e., the overall activity level of an optic glomerulus); and (3) a decision network with dense connections that maps LC responses to behavior, reflecting downstream processing in the central brain and ventral nerve cord. We imposed the bottleneck layer to have the same number of units as LC neuron types we manipulated, and our goal was to identify a one-to-one mapping between these model LC units and the LC neuron types. We did not incorporate strong biological realism into the vision and decision networks [39], opting instead for highly-expressive mappings to ensure high prediction; we focused on the model’s LC bottleneck. We collected training data to fit the model by genetically inactivating each of 23 different LC types in male flies [11, Nern et al., *in prep.*] and recorded the LC-silenced male’s movements and song production during courtship (see Methods). We then devised a fitting procedure called ‘knockout training’, which involves training the model using the entire dataset of perturbed and unperturbed behavior. Critically, when training the model on data from a male with a particular LC type silenced (Fig. 1**b**), we set to 0 (i.e., knocked out) the activity of the corresponding model LC unit (correspondence was arbitrarily chosen at initialization, see Methods). The resulting model captures the behavioral repertoire of each genetically altered fly when the corresponding model LC unit is silenced, thereby aligning the model LC units to the real LC neurons. In simulations (Ext. Data Fig. 2), knockout training correctly identified each silenced neuron type’s activity and contribution to behavior (i.e., a one-to-one mapping) for neuron types that, when silenced, led to changes in behavior. We refer to the resulting DNN model as the 1-to-1 network.

**Figure 2:**
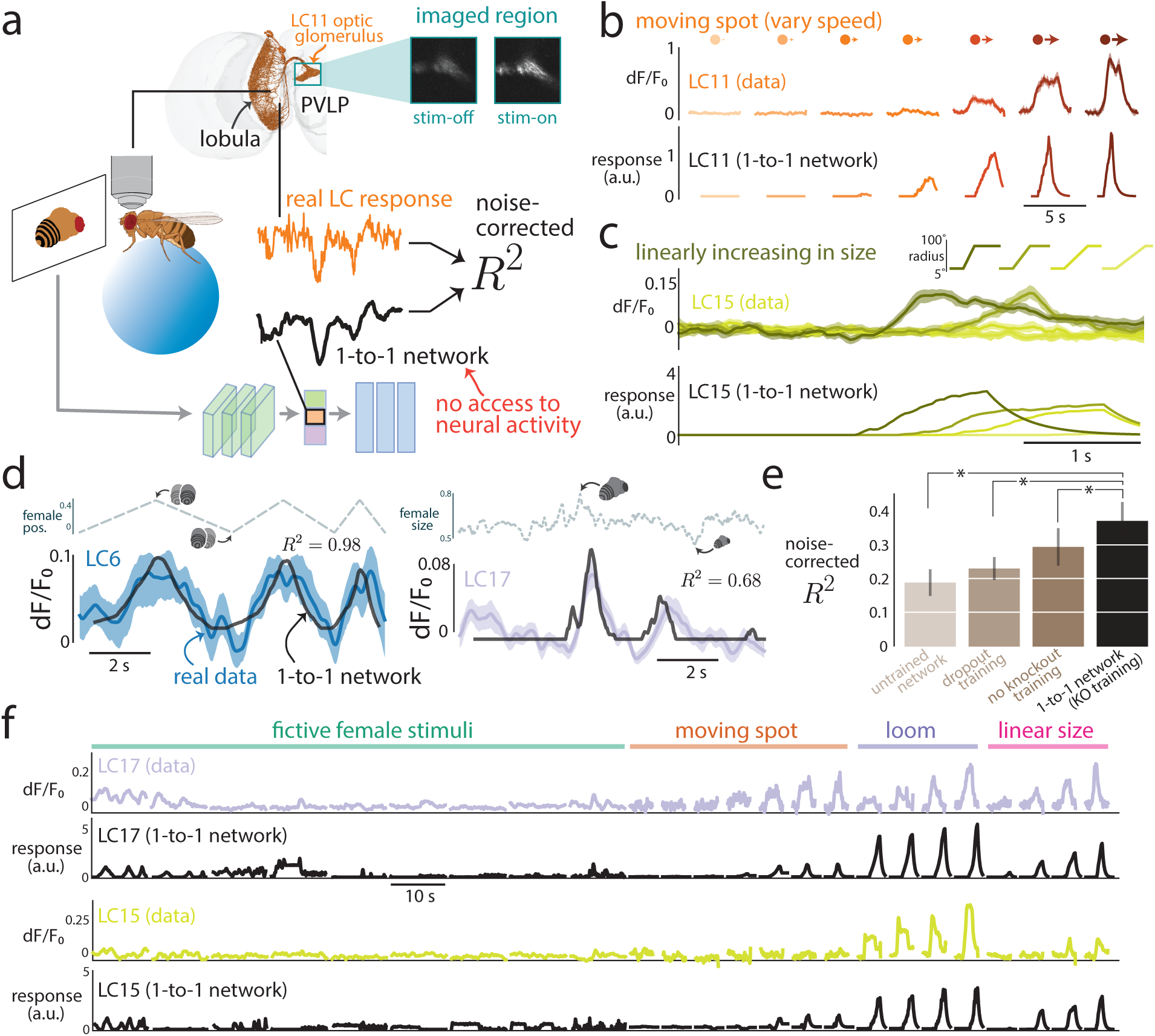
Model LC responses from the 1-to-1 network match real LC neural responses. **a**. We recorded LC responses using calcium imaging while a head-fixed male fly (walking on a moveable, air-supported ball) viewed dynamic stimulus sequences of a moving spot or a fictive female fly on a projection screen (see Methods). We fed the visual stimulus sequence as input into the 1-to-1 network and asked if the predicted responses (black trace) for a given model LC unit matched the real response of the corresponding LC neuron (green trace) by comput-ing the noise-corrected *R*^2^ between the two (normalized) traces over time (see Methods). A noise-corrected *R*^2^ of 1 indicates that a model perfectly predicts repeat-averaged responses after taking into account the amount of variability across repeats. The 1-to-1 network never had access to real LC responses during training (behavior only) and could only use one internal model LC unit for prediction. We targeted LC6, LC11, LC12, LC15, and LC17, each of which has neurons whose dendrites span the lobula and converge onto the same optic glomerulus (inset for LC11 neurons taken from FlyWire connectome [41] with the Neuroglancer software [42]). We image and take as the response the summed calcium dynamics within the region occupied by the glomerulus (insets, ‘stim-off’/‘stim-on’: activity before/during visual stimulus was presented). **b**. Real responses for LC11 to a visual stimulus in which a spot moved left to right at different speeds (top row). Model LC responses to this stimulus se-quence (bottom row) qualitatively matched the real LC responses. Because the model was only trained on images of a fictive female, for model input we replaced the spot with a fictive female facing away from the male with the same size and location of the spot for each frame (see Methods). **c**. Real responses for LC15 to a spot with linearly increasing size (top row). Model LC responses computed in the same manner as in **b** (bottom row). **d**. We also presented various stimulus sequences of a fictive female fly (top traces denote the values of a visual parameter such as female position or size) and collected both real LC responses (color traces) and model LC responses from the 1-to-1 network (black traces). Shaded regions denote 90% bootstrapped confidence intervals of the mean across repeats; noise-corrected *R*^2^s for these example stimuli and responses are indicated. **e**. Average noise-corrected *R*^2^ across all stimulus sequences and LC types for differ-ent models. The average *R*^2^ of the 1-to-1 network (*R*^2^s in **d** contribute to this average) was significantly higher than each average *R*^2^ of the other models (asterisks; *p <* 0.05, paired one-sided permutation test). The *R*^2^ for the untrained network was not 0, suggesting that the inductive bias of convolution as well as the small number of the stimuli meant a network with random weights could have some prediction (see Methods). Error bars indicate 1 s.e.m. across stimulus sequence/LC pairs. **f**. Real (color traces) and model LC responses (black traces) across all stimulus types, including naturalistic (e.g., fictive female stimuli in **d** and **e**) and artificial stimuli (moving spots, loom, and linear size; e.g., in **b** and **c**). Color traces denote mean LC response. Model LC responses were read out directly from the network and were not smoothed or scaled as was done to compute *R*^2^ (see Methods). LC17 and LC15 are shown here; LC6, LC11, and LC12 responses are shown in Extended Data Figure 10.

Before fitting the model with courtship data (Fig. 1**c**), we quantified the extent to which silencing (bilaterally) each LC neuron type changes behavior of the male fly. We wondered if only a select few LC types, once silenced, would result in large changes in behavior. Consistent with previous studies [10, 22], we confirmed that silencing LC10a neurons resulted in failures to initiate chasing, as male-to-female distances remained large over time (Fig. 1**d**, middle panel, and Fig. 1**e**, top panel). On the other hand, silencing LC6 neurons resulted in stronger and more persistent chasing as male-to-female distances remained small over time (Fig. 1**d**, bottom panel, and Fig. 1**e**, top panel). LC6 has never been implicated in courtship. We found that none of these changes matched the behavioral deficits of blind flies (Ext. Data Fig. 1), suggesting no single LC type is the sole contributor to the male’s courtship behavior. This was also true of song production, as measured by the different proportions of song type (sine or pulse song, Fig. 1**e**, bottom panel), as well as other behavioral measures (Ext. Data Fig. 1). This suggests that many of the LC types would need to be silenced together for large behavioral deficits to occur. To better understand how these LC types work together as a population, we modeled this perturbed behavioral data with the 1-to-1 network, with which we can silence any possible combination of LC types.

We performed knockout training to fit the parameters of the 1-to-1 network. The model inputs were images that reflected the visual input of the male fly (i.e., a fictive female fly changing her size, position, and rotation; see Methods) (Fig. 1**f**, left panel); the model outputs comprised the male movements (forward, lateral, and angular velocity) and song production, which included sine song and two forms of pulse song (Pfast and Pslow [29]). The 1-to-1 network reliably predicted these behavioral variables in held-out data (Fig. 1**f**, right panels, and Ext. Data Fig. 3). Importantly, the 1-to-1 network also predicted differences in behavior observed across silenced LC types, such as changes in the male’s forward velocity (Fig. 1**g**), sine song production (Fig. 1**h**), and other behavioral outputs (Ext. Data Fig. 4). We confirmed that knockout training outperformed other possible training procedures, such as dropout training [40] and training without knockout (i.e., an unconstrained network) (Ext. Data Fig. 3 and 4), and that results were consistent for different random initializations of the 1-to-1 network (Ext. Data Fig. 5). Thus, the 1-to-1 network reliably estimated the male’s behavior from visual input, even for male flies with a cell type silenced.

**Figure 3:**
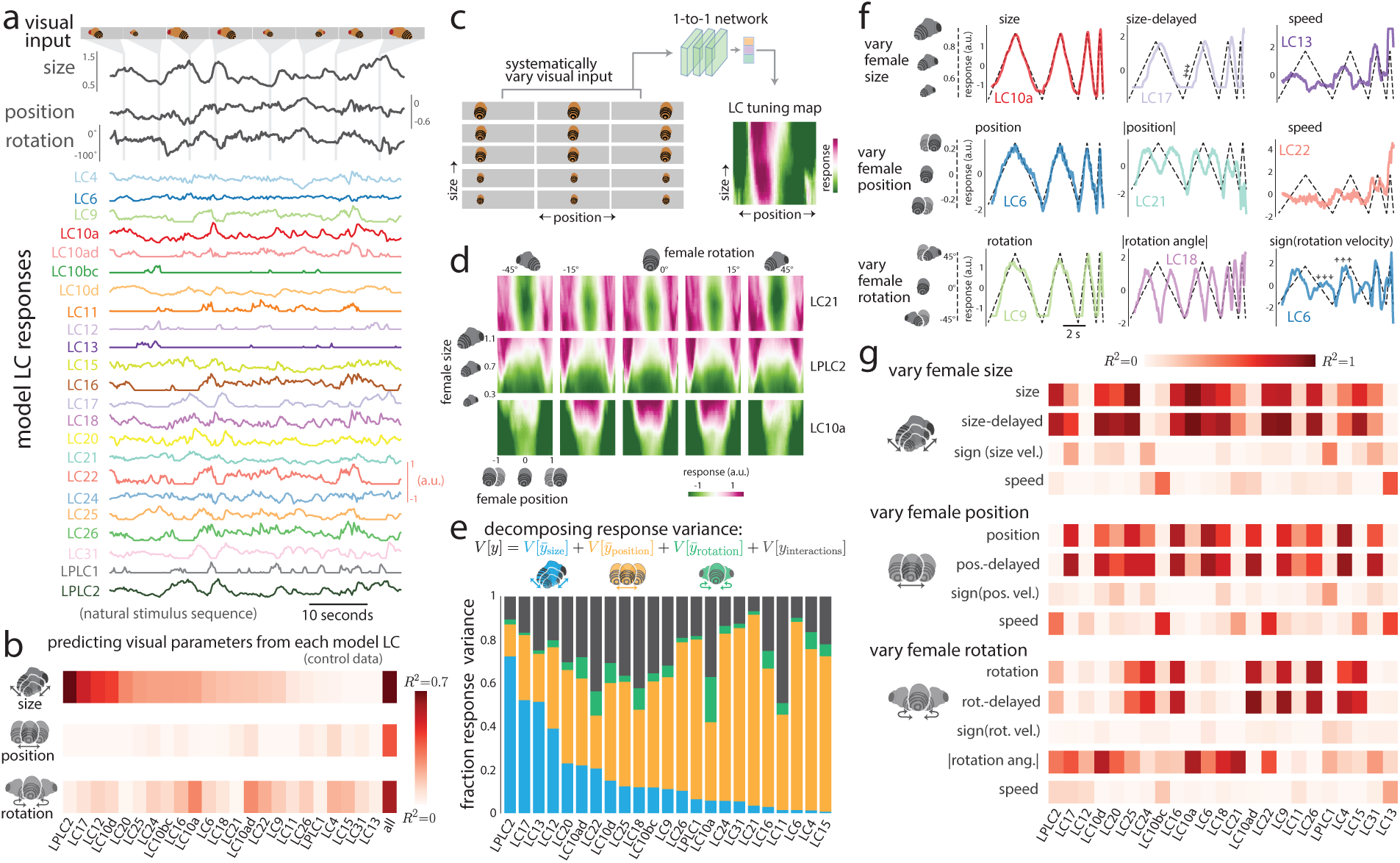
Model LC units combinatorially encode the features of female motion during courtship. **a**. Model LC responses to a natural stimulus sequence in which a fictive female changes her size, position, and rotation. **b**. Cross-validated *R*^2^ between each primary visual parameter and model LC responses for stimulus sequences ob-served during natural courtship (assessed on heldout test stimuli from control flies). Columns are sorted based on female size (first row). The end column of each row (‘all’) is the cross-validated *R*^2^ between a linear combination (identified via ridge regression) between all model LC units and a single visual parameter. Because female position and rotation are circular variables, we converted each variable *x* to a 2-d vector [cos(*x*), sin(*x*)] and took the maxi-mum *R*^2^ across both variables for each model LC unit. **c**. To characterize the stimulus preferences of each model LC unit, we systematically varied the visual parameters of female size, position, and rotation. Because the 1-to-1 network takes in as input a sequence of 10 frames, for simplicity we repeated the same image of the fictive female for all 10 frames for a given set of parameter values (i.e., the input was static). For each model LC unit, we com-puted a heatmap of the LC responses to visualize the unit’s tuning preferences. **d**. LC tuning curves as heatmaps for three example model LC units. Female size and position varies within each 2-d heatmap; female rotation varies across columns. **e**. We used variance decomposition (see Methods) to decompose the response variance of each model LC unit (i.e., the total variance of responses to the systematically-varied stimulus sequences in **c** and **d**) into a component solely due to either female size (blue), position (orange), and rotation (green) as well as interactions between these three visual parameters (black). A large fraction of response variance for a given visual parameter indicates that a model LC unit strongly changes its response to variations in this parameter relative to variations in other parameters. Because the 1-to-1 network is deterministic, all response variance can be attributed to variations of the visual parameters (i.e., no component of the variability can be attributed to ‘noise’ across repeats of the same stimulus). **f**. Example model LC responses to dynamic stimulus sequences in which the fictive female varied either her size, position, or rotation angle over time while the other two parameters remained fixed (dashed lines; dashed y-axis values correspond to plots in the first column only). Different model LC units appear either to di-rectly encode a visual parameter (e.g., LC10a encodes ‘size’) or encode features derived from the parameter, such as a delay (LC17, arrows) or speed at which female size changes (LC13). Responses for all model LC units are in Extended Data Figure 12. **g**. *R*^2^ between model responses and visual parameter features for the three different stimulus sequences in **f**. Columns have the same ordering as that of **b**.

**Figure 4:**
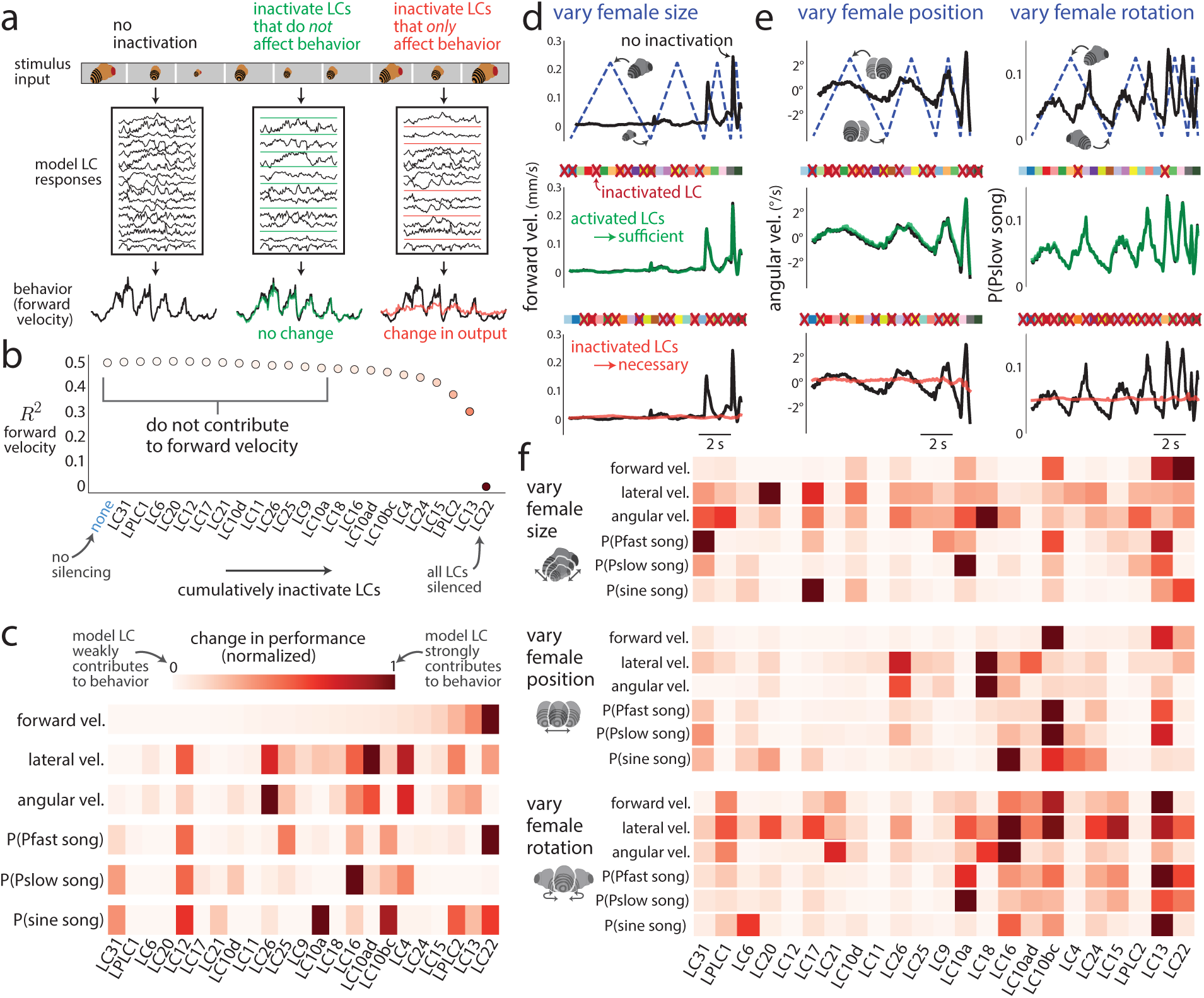
Model LC units form a combinatorial code for behavior. **a**. Illustration of our approach to investigate which model LC units contribute to which behavioral outputs for any stimulus sequence. We can assess if a group of model LC units are sufficient and necessary for behavior if we inactivate all model LC units not in that group (middle panel, ‘sufficient’) or inactivate only that group of model LC units (right panel, ‘necessary’). **b**. We iden-tify which model LC units contribute to forward velocity via cumulative inactivation for stimulus sequences in the heldout test data of control sessions. At each iteration, we inactivate the model LC unit that, once inactivated, maintains the best prediction of forward velocity (assessed by *R*^2^ between predicted and real, heldout behavior of control flies) while still inactivating all previously-inactivated model LC units (i.e., cumulative inactivation in a greedy manner). The inactivated model LC units that lead to the largest decreases in *R*^2^ (e.g., LC13 and LC22) contribute the most to forward velocity. Dot color corresponds to *R*^2^ values in **c**. **c**. Results for cumulative inactivation for all 6 behavioral outputs; forward velocity (top row) is the same as in **b**. For movement variables, normalized change in performance is the difference in *R*^2^ between no silencing (‘none’) and silencing *K* model LC units, normalized by the *R*^2^ of no silencing. For song variables, normalized change in performance is the same as for the movement variables except we use 1 *− c.e.* Columns of each row are ordered based on the ordering of for-ward velocity (top row). For exact values, along with single-unit inactivation results, see Extended Data Figure 14. **d**. We considered the model’s predicted behavior for a simple, dynamic stimulus sequence (e.g., dashed line in top panel: varying the female’s size over time while her position and rotation remained fixed). Inactivating a chosen group of model LC units (red Xs denote inactivation; model LC units are represented by squares whose colors and ordering match those in Fig. 3a) led to almost no change in the model’s output (middle panel, green trace overlays black trace). Thus, the remaining activated model LC units (squares with no red X) are sufficient to produce the model’s output. Inactivating these “sufficient” model LC units (bottom panel, red Xs are swapped from squares in middle panel) led to a large behavioral deficit (bottom panel, red trace does not match black trace), indicating that these model LC units are also necessary. Thus, this group of model LC units is both necessary and sufficient for the 1-to-1 network’s prediction of forward velocity for a stimulus sequence in which the female’s size is varied. **e**. Other example behavioral outputs and stimulus sequences to assess necessity and sufficiency. Same format is in **d**. **f**. Results of cumulative inactivation for the three dynamic stimulus sequences in **d** and **e**. Same format, color legend, and ordering of columns as in **c**.

**Figure 5:**
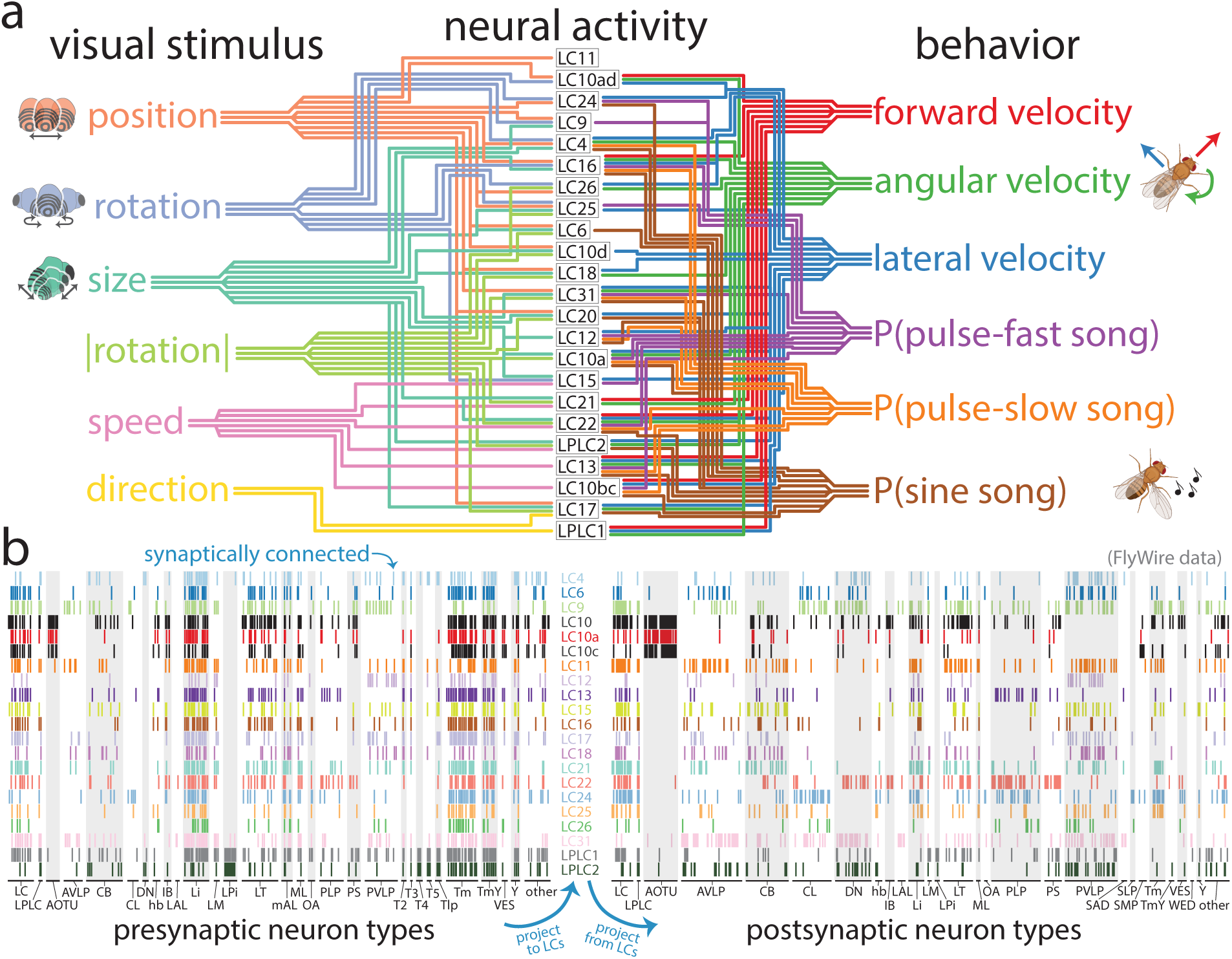
The role LC neurons play in the sensorimotor transformation of the male fly during courtship. **a**. Summary of our findings. Each line denotes a relationship between a model LC unit and either a visual feature (left, *R*^2^ *>* 0.30 in Fig. 3b or **g**) or a behavioral variable (right, a normalized change in performance greater than 30% in Fig. 4c or **f**). A lack of connection does not rule out a relationship, as relationships may exist in other con-texts or subcontexts. Even at these conservative criteria (i.e., cutoffs at 0.3), many model LC units encode more than one visual feature and contribute to more than one behavioral variable. **b**. Synaptic connectivity matrices for presynaptic neuron types projecting to LC or LPLC neurons (left) and postsynaptic neuron types receiving input from LC or LPLC neurons (right). Each row is for one LC/LPLC type. Each column is for a partner neuron type; columns are grouped into classes of neuron types/brain areas based on the naming conventions in FlyWire’s connectome dataset [45]. A tick line indicates at least 5 synaptic connections were identified between neurons of an LC/LPLC neuron type and neurons of a pre-or postsynaptic neuron type. We include synaptic connections for LC10 and LC10c, which were not one of the LC types we silenced (**a**) but were present in the FlyWire connec-tome; we leave out LC10d and LC20, which were silenced but have not yet been identified in FlyWire. Because this connectome dataset is from a female fruit fly, it may miss important sexually-dimorphic, courtship-relevant connections to downstream areas of the male fruit fly.

## Comparing real and model neural activity

Does the knockout training procedure, which leverages natural behavioral data only, enable model LC units in the 1-to-1 network to match the response properties of real LC neurons? To address this question, we recorded LC calcium dynamics in head-fixed, passively-viewing male flies expressing GCaMP6f (Fig. 2**a**, see Methods); because of the differences in behavior during courtship (on which the 1-to-1 network was trained) and behavior of the head-fixed male, we did not expect exact matches between predicted and real responses. We targeted 5 differ-ent LC neuron types (LC6, LC11, LC12, LC15, and LC17), chosen because silencing each one led to noticeable changes in courting behavior (Fig. 1**e** and Ext. Data Fig. 1) and LC10a responses to courtship-like stimuli have previously been recorded [10, 22]. We first presented an artificial spot moving laterally or enlarging (Fig. 2**b** and **c**, see Methods) commonly used to characterize LC responses in previous studies [30, 31, 32]. Despite the fact that the 1-to-1 network never had access to neural data, we found that its predicted responses largely matched their cor-responding real LC responses (Fig. 2**b** and **c**, compare top and bottom panels). We confirmed for a larger number of LC types that the model LC responses were qualitatively similar (Ext. Data Fig. 7). These matches surprised us, as these artificial stimuli were likely rare or not present in natural courtship data on which the 1-to-1 network was trained.

Given these matches for artificial stimuli, we then tested the 1-to-1 network’s predictions on more naturalistic stimulus sequences (i.e., a fictive female varying her position, size, and rotation; see Supplementary Video 1)—LC responses to such naturalistic stimuli have rarely been collected before. We found that the recorded LC neurons responded to many of these naturalistic stimulus sequences (Fig. 2**d**, color traces, and Ext. Data Fig. 8) and found good matches between real LC responses and their corresponding model LC responses (Fig. 2**d**, black traces versus color traces, and Ext. Data Fig. 8). Quantitatively, the 1-to-1 network predicted real LC responses across stimuli and LC neuron types with an average noise-corrected *R*^2^ of *∼*0.35, close to double that of an untrained network (Fig. 2**e**). This indicates that training on behavioral data was helpful in predicting real LC responses. The 1-to-1 network also better predicted real responses versus other networks with the same architecture but trained with dropout or without knockout procedures (Fig. 2**e**). That these networks outperformed an untrained network supports the distributed nature of the LC neuron types. For example, dropout training encourages a redundant code and finding a match between a randomly-chosen model LC unit and an LC neuron type becomes more likely with higher redundancy in the code.

We further tested the 1-to-1 network’s predictions in three ways. First, because our noise-corrected *R*^2^ metric normalized real and predicted responses for each stimulus, we assessed the extent to which the 1-to-1 network predicted response magnitudes across both natural and artificial stimuli. We found that the selectivity of the un-normalized 1-to-1 network responses largely matched those of the real LC neurons (Fig. 2**f** and Ext. Data Fig. 10); this was especially true for LC15, as the 1-to-1 network correctly identified LC15’s strong selectivity for looming stimuli (Fig. 2**f**, ‘LC15’). Our second test gave the 1-to-1 network access to neural data by training a linear mapping between all 23 model LC units and one real LC neuron type. We found that held-out prediction improved to a noise-corrected *R*^2^ of *∼*0.65 (Ext. Data Fig. 9). To our knowledge, the 1-to-1 network with a fitted linear mapping is the first highly-predictive, image-computable encoding model of LC population responses. Finally, our third test asked to what extent these trends held for 10 different random initializations (Ext. Data Fig. 8). We found that the 1-to-1 network was the most consistent in its neural predictions across initializations versus other training procedures (Ext. Data Fig. 6). An additional use of these 10 models is to assess the uncertainty the 1-to-1 network may have about its predictions; a large disagreement among the models (i.e., a large variability) for a particular stimulus sequence indicates that we should not expect the 1-to-1 network to accurately predict responses to this stimulus, a helpful guide for future experiments.

Overall, we conclude that the 1-to-1 network predicts changes in behavior due to silencing (Fig. 1) and neural responses to natural and artificial stimuli (Fig. 2). This motivates us to investigate the inner workings of the 1-to-1 network. For simplicity, we chose to analyze the 1-to-1 network from the 10 different initializations that led to the best predictions in behavior and neural responses (see Methods).

## Encoding of visual stimulus features by the model LC population

We next investigated how the population of 23 model LC units encodes visual stimuli. Previous studies have recorded LC responses (mostly in non-behaving, head-fixed flies) to simple visual stimuli, such as moving dots, bars, and gratings. With the 1-to-1 network, we can now probe how the LC types encode more naturalistic stimuli, such as a fictive female changing her size, position, and rotation based on actual courtship statistics. The simplest code would have three model LC units responding to and encoding the three visual parameters separately. Instead, we found that the majority of model LC units in the 1-to-1 network responded to a natural stimulus (Fig. 3**a**). Despite this, almost no model LC units linearly encoded any single visual parameter of female size, position, or rotation (Fig. 3**b**, low *R*^2^s for any one LC type). We needed a linear mapping comprising all model LC units to reliably encode each visual parameter (Fig. 3**b**, ‘all’, high *R*^2^). This suggests that the 1-to-1 network relies on multiple LC neuron types to encode a single parameter, consistent with a distributed encoding scheme. That multiple real LC neurons responded in some way to the same artificial stimulus sequences (Ext. Data Fig. 7) and naturalistic stimulus sequences (Ext. Data Fig. 8, ‘natural sequence’) is consistent with this conclusion.

We further investigated the ‘tuning’ of each model LC unit by systematically varying all three primary parameters of the female visual input and passing these input sequences into the model—we assembled the responses for each model LC unit into a 3-dimensional ‘tuning curve’ (Fig. 3**c** and Ext. Data Fig. 11). While some model LC units were largely driven by a single parameter, such as model LC21 tuned to female position (Fig. 3**d**, top row) and model LPLC2 tuned to female size (Fig. 3**d**, middle row)—consistent with previous work [24], other model LC units were tuned to interactions between two or more parameters (Fig. 3**d**, bottom row, model LC10a more strongly encodes position for females with large sizes and facing away from the male but not for small female sizes). To quantify these interactions, we decomposed the response variance [43] of each model LC unit into four components: the response variance solely due to either female size, position, or rotation and any remaining response variance (capturing interactions among two or more of the visual parameters, see Methods) (Fig. 3**e**). Most model LC units encoded changes in female position (Fig. 3**e**, orange bars), roughly half encoded female size (Fig. 3**e**, blue bars), and female rotation was weakly encoded (Fig. 3**e**, green bars are small for all model LC units). However, almost all model LC units encoded some nonlinear interaction among the three visual parameters (Fig. 3**e**, black bars, on average *∼*25% of the response variance for each model LC unit). These interactions would not have been discovered had we only varied one visual parameter at a time—motivating the use of more complex stimuli to probe LC function.

The analyses presented so far ignore changes to the visual input over time, but LC neurons do respond to dynamically-changing stimuli [30, 31, 32]. Thus, we considered dynamic stimulus sequences in which the fictive female varies in one visual parameter while the other two parameters remain fixed (Fig. 3**f**, dashed lines, see Supplementary Video 2). For example, we varied the female’s size over time with different speeds (Fig. 3**f**, top row, dashed lines). We found that some model LC units perfectly encoded female size (Fig. 3**f**, top left panel, LC10a), some model LC units encoded a time-delayed version of size (Fig. 3**f**, top row, middle panel, LC17), while other model LC units encoded the speed at which female size changed (Fig. 3**f**, top row, right plot, LC13). Similar relationships were present for other stimulus sequences and model LC units (Fig. 3**f**, bottom two rows); we note that the 1-to-1 network was predictive of real LC responses for similar types of stimulus sequences (Fig. 2**d**-**e** and Ext. Data Fig. 8). Across the three stimulus sequences, the primary visual features of female size, position, and rotation were strongly encoded (Fig. 3**g**, ‘size’, ‘position’, and ‘rotation’) with speeds and the absolute value of rotation angle also encoded. The model’s predictions were consistent with findings from previous studies, such as LC11 encoding the position of a small moving spot [26, 27] (Fig. 3**g**, LC11 has higher *R*^2^ for ‘position’ in ‘vary female position’ than in other stimulus sequences) and LPLC2 encoding loom [23] (Fig. 3**g**, LPLC2 has high *R*^2^ for ‘size’ in ‘vary female size’). We also observed that model LC10a responses do encode female position, consistent with previous findings [10, 22], but in a nonlinear way (Ext. Data Fig. 12). Finally, model units LC4, LC15, LC16, LC17, LC18, LC21, LC25, and LC26 all strongly encoded female size (Fig. 3**g**, topmost row), matching recent findings that these LC neurons respond to looming and moving objects of various sizes [30, 31, 32].

Taking all of these results together, we found that almost all of the model LC units encode some aspect of female size, position, rotation, or motion during courtship. In addition, many model LC units encoded information about the same parameter (the average signal correlation across model LC units for natural stimulus sequences was *|ρ|* = 0.29). Indeed, the response-maximizing stimulus sequence for each model LC unit strongly drove responses of other model LC units, even when optimized for these other responses to be suppressed (a “one hot activation”, Ext. Data Fig. 13). We conclude that the model LC units form a distributed, combinatorial code for visual stimuli: Each visual stimulus feature is encoded by multiple model LC units (i.e., Fig. 3**g**, rows each have multiple red squares) while each model LC unit encodes multiple visual stimulus features (i.e., Fig. 3**g**, columns each have multiple red squares). Given the distributed nature of the LC code, we wondered whether model LC units also form a population code where multiple LC types are read out to drive male courtship behavior. To test this, we next asked whether a majority of the model LC units were necessary and sufficient to perform different actions during courtship.

## Model LC units form a population code for behavior

During courtship, male flies chase, orient towards, and sing to females. Each of these behaviors could be separately governed by individual or groups of LC neuron types [12, 13]. However, our results showing that model LC units combinatorially encode the motion of the female (Fig. 3) suggest instead that a combination of LC types helps to drive male behaviors toward the female. Here, we directly test this hypothesis by systematically inactivating model LC units in all different combinations (or alone)—experiments not easily performed in a real fly—and then examined which model LC units were necessary and sufficient to guide behavior (Fig. 4**a**).

We started by asking which model LC unit, when inactivated, maintained the best performance in predicting the behavior of control flies. For predicting male forward velocity, for example, inactivating LC31 had a negligible effect on prediction performance (Fig. 4**b**, ‘LC31’ vs ‘none’), leading us to conclude that model LC31 does not contribute meaningfully to male forward velocity (although model LC31 contributes to song production, see Fig. 4**c**). Next, in a greedy and cumulative manner, we repeatedly inactivated the model LC unit that maintained the best performance while keeping all previously-chosen LCs inactivated; eventually prediction performance had to decrease (Fig. 4**b**, rightmost dots) because of the bottleneck imposed by the model LC units. The inactivated model LC units that led to the largest drops in performance were the strongest contributors to behavior (Fig. 4**b**, rightmost dots). Interestingly, separately inactivating each model LC unit resulted in little to no drop in prediction performance, including the strongest contributor model LC22 (Ext. Data Fig. 14). This suggests that only by inactivating a combination of LC types will large changes in behavior be observed, consistent with our finding that inactivating any single LC type did not lead to the large deficits in behavior observed for blind flies (Ext. Data Fig. 1).

Next, we performed this cumulative inactivation procedure for all 6 behavioral outputs (Fig. 4**c** and Ext. Data Fig. 14). We observed that different model LC units contributed to different behaviors (Fig. 4**c**, e.g., compare rows ‘forward velocity’ and ‘lateral vel’) and that most model LC units each contributed to multiple behavioral outputs (Fig. 4**c**, multiple red squares per column), consistent with previous work [11]. These results indicate that to a large degree model LC units operate as a distributed population in which almost all model LC units contribute in some way to the courtship behaviors examined here.

Because natural behavior is an aggregate over many different behavioral contexts, it may be the case that only a few LC types participate in simpler contexts. This motivated us to consider the simple, time-varying stimulus sequences for which we recorded responses of a subset of real LC neurons (Fig. 2**d**) and that we used to analyze model LC tuning properties (Fig. 3**f** and **g**). Through systematic inactivation, we again identified the model LC units that were both necessary and sufficient to produce the model’s output to these more artificial stimuli, each representing a different context. For example, we found that when we varied female size only (Fig. 4**d**, top panel, dashed line), inactivating 10 different model LC units (Fig. 4**d**, middle panel, squares with red Xs, identified via cumulative inactivation, see Methods) resulted in no change in forward velocity (Fig. 4**d**, middle panel, green trace overlays black trace, and Ext. Data Fig. 15). This suggests that the *other* activated model LC units (Fig. 4**d**, middle panel, squares without a red X) were sufficient to drive behavior. We then inactivated these “sufficient” model LC units (keeping all other model LC units activated) and found a large behavioral deficit (Fig. 4**d**, bottom panel, red trace does not overlay black trace), indicating that these inactivated model LC units were also necessary. We show examples for the same type of analysis for the male’s angular velocity in response to varying female position (Fig. 4**e**, left column) and for the production of Pslow song in response to varying female rotation (Fig. 4**e**, right column). In the latter, all but 2 model LC units (LC11 and LC25) had to be inactivated to extinguish behavior (Ext. Data Fig. 15), although not every model LC unit contributed as strongly.

Across all behavioral outputs, even for these simple stimulus sequences, we found that combinations of multiple model LC units were necessary to drive behavior. This is supported by the observations that multiple LC units contributed to the same behavior (Fig. 4**f**, multiple red squares per row) and that most model LC units each contributed to multiple behaviors (Fig. 4**f**, multiple red squares per column). Indeed, no single model LC unit was necessary and sufficient for these behaviors (Ext. Data Fig. 16), consistent with our silencing experiments (Ext. Data Fig. 1). In addition, the strengths of model LC contributions changed between natural courtship and these dynamic stimulus sequences (compare Fig. 4**c** and **f**, ‘forward velocity’; columns have same ordering). This is not unexpected: An LC neuron type may not contribute to behavior in these chosen stimulus contexts (e.g., white squares for LC12 in Fig. 4**f**) but may contribute in other contexts, leading to the LC type being a contributor in natural courtship (e.g., red squares for LC12 in Fig. 4**c**). Overall, our results support the notion that the model LC units form a distributed population code.

Previous studies have identified sexually dimorphic LC10a neurons to be involved in tracking the female during courtship chasing [10, 22], consistent with our own observation that silencing LC10a neurons reduced male velocity relative to control (Ext. Data Fig. 1). Here, we found that model LC10a does contribute to the male’s movement but also to song production (Fig. 4**c**, LC10a and sine song), a novel finding to be tested in future ex-periments. We found other matches to previous findings. For example, we also find that LC9, another LC type implicated in courtship [21], contributes to tracking of the female (Fig. 4**f**, LC9 and ‘vary female size’). LC16, LC17, and LPLC1 are strong contributors to angular velocity (Fig. 4**f**, ‘vary female size’) and, if optogenetically activated, produce turning behavior [11]. This is also consistent with LC16 contributing to backward turns [25]. LC4 and LPLC2, known to encode loom and contribute to escape responses [23, 24], both were strong contributors to forward velocity (Fig. 4**c**). One interesting prediction of the 1-to-1 network is that LC11, known to be a small object detector [26], plays a subtle role in guiding behavior, as model LC11 contributed little to behavior in these contexts (Fig. 4**c** and **f**, ‘LC11’) yet the 1-to-1 network predicted its object size selectivity (Fig. 2**b** and Ext. Data Fig. 7). The 1-to-1 network also makes new predictions, such as LC22 playing a major role in forward velocity and Pfast song production as well as LC13 encoding the speed of visual parameters (Fig. 3**g**) and strongly contributing to movement and song production. These predictions—as well as larger questions, such as how LC neurons play a role in the male’s decision to sing pulse or sine song [44]—can be tested in future experiments, guided by the 1-to-1 network.

## Distributed inputs and outputs for cell types in the connectome

We aggregated results both from how the model LC neurons encode visual input (Fig. 3) and contribute to behavior (Fig. 4) and outline these relationships (Fig. 5**a**). The picture is complicated: Many model LC units encode multiple visual features of the female (Fig. 5**a**, left connections, a “distributed" code) and contribute to multiple behavioral outputs (Fig. 5**a**, right connections, a “population” code). Given our model’s ability to predict both the changes to behavior during genetic silencing (Fig. 1**g** and **h**) and responses of real LC neurons (Fig. 2), we propose that the LC neurons in the fly visual system also form this complex, distributed population code. We note that these predictions (e.g., connecting lines in Fig. 5**a**) come from one training run of the 1-to-1 network (see Methods); we can assess the uncertainty of each connection by observing differences in predictions across different training runs (Ext. Data Figs. 5 and 6). Future experiments can test the most uncertain connections as predicted by the model.

A distributed population code would require a rich scaffolding of presynaptic connections to read from and combine a variety of visual features as well as postsynaptic connections giving downstream areas access to integrate visual information from multiple LC types. To see if this were the case for the LC neurons, we analyzed a recently-released whole-brain connectome called FlyWire [45, 46, 47] in which 35 LC and LPLC neuron types out of the *∼*45 known LC types have been identified so far. We computed the synaptic connectivity matrix for LC neuron types and their presynaptic partners (Fig. 5**b**, left) as well as their postsynaptic partners (Fig. 5**b**, right), where each entry was either 1 (denoted by a tick line) if at least 5 synapses were identified between the neurons of a given LC type and neurons of a pre-/postsynaptic type, else 0. This analysis was aided by exhaustive cell typing in the optic lobe [17]. We found evidence in support of a distributed code, as multiple upstream neurons of different types projected to neurons of the same LC type (Fig. 5**b**, left, most rows each have ticks for multiple groups, e.g., ‘LC31’) and neurons of multiple LC types read out upstream neurons of the same type (Fig. 5**b**, left, multiple ticks per column, e.g., ‘T3’)—62% of presynatpic neuron types projected to 2 or more LC types and 48% to 3 or more LC types. Likewise, we found evidence in support of a population code, as neurons from multiple LC types innervated downstream neurons of the same neuron type (Fig. 5**b**, right, each row has ticks for multiple groups, e.g., ‘LC22’), suggesting each LC type contributes to a variety of behaviors [11]. In addition, most postsynaptic partners received input from multiple LC types (Fig. 5**b**, right, multiple ticks per column, e.g., ‘PVLP’)—in fact, 54% of downstream neurons of the same type received input from 2 or more LC types and 31% from 3 or more LC types, suggesting that downstream areas integrate information from multiple LC types. An additional observation not yet reported is that many LC and LPLC types directly connect with other LC and LPLC types in the lobula and lobula plate (Fig. 5**b**, ‘LC’ and ‘LPLC’ columns in both left and right synaptic connection matrices have many ticks across rows and columns). Such recurrence muddles the idea that each LC type is an independent feature detector, although these lateral connections may implement a winner-takes-all mechanism such as divisive normalization [48, 49] or may help to sparsify the code [50]. An important caveat is that this connectome dataset is from a female fruit fly; once a male’s connectome is obtained, we can test the 1-to-1 network’s predictions about which LC types contribute to which behavioral outputs (Fig.5**a**) by assessing which downstream areas read out from which optic glomeruli (Fig. 5**b**, ‘post-synaptic neuron types’) and how those downstream areas contribute to behavior.

## Discussion

Why would the fly visual system, with its highly-organized set of optic glomeruli, use a distributed population code of LC neurons? Such a code can bring computational advantages, such as noise robustness [51, 52, 53, 54], flexibility to accommodate multiple tasks [14, 55, 56, 57, 58, 59], and coding efficiency [50, 52, 60, 61, 62]. It may be the case that in an early point in evolutionary history, the optic glomeruli were spatially arranged to assist in the quick processing needed for important reflexes, such as escaping from a looming predator [20, 23, 63]. However, as the visual and motor systems adapted to changing environmental pressures, these systems may have updated their “software” (reusing LC neuron types for different behaviors) while keeping the same “hardware” (a fixed number of glomeruli) to allow for a richer repertoire of complex behaviors (e.g., courtship, flight, landmark navigation, color discrimination, etc.) while still maintaining quick reflex circuitry. To understand how LC types are employed during the complex behavior of courtship, we developed knockout training to identify a one-to-one mapping between internal units of a DNN and real neurons. The resulting 1-to-1 network makes testable predictions, including which stimulus sequences evoke large LC responses (Ext. Data Fig. 13) and which combinations of LC types are necessary and sufficient for specific behaviors (Fig. 4). A major new finding of our work is which and to what extent LC neuron types contribute to song production, an integral part of courtship [9]. We suspect that adding more biological realism to our DNN model, such as incorporating information from the connectome of the optic lobe [16, 39] or building in recurrent connections, lateral connections between LC types (Fig. 5**b**), and delays, will lead to knockout training better identifying this one-to-one mapping with enough training data. An intriguing direction is to apply this framework to other complex visual behaviors, such as flight and navigation, and other sensory-driven behaviors, such as odor plume tracking (where each olfactory glomerulus can be genetically silenced separately). Our work shows that constraining models with causal perturbations of neurons during complex behavior is an important ingredient in revealing the relationships between stimulus, neuron, and behavior.

## Methods

### Flies

For all experiments, we used 4-7 day old virgin flies harvested from density-controlled bottles seeded with 8 males and 8 females. Fly bottles were kept at 25°C and 60% relative humidity. Virgined flies were then housed individually across all experiments. Female virgined flies were individually housed and kept in behavioral incubators under a 12-12 hr light-dark cycling; individual males were paired with a PIBL (pheromone insensitive and blind) female—see Supplemental Table 1 for more info on genotype. UAS-TNT-C was obtained from the Bloomington stock center. All LC lines and the spGAL4 control line [64] were generously provided by M. Reiser, A. Nern, and G. Rubin—see Supplemental Table 1 for more information.

### Courtship experiments

Behavioral chambers were constructed as previously described [9, 65]. Each recording chamber had a floor lined with white plastic mesh and equipped with 16 microphones (Ext. Data Fig. 1). Video was recorded from above the chamber at a 60 Hz frame rate; features for behavioral tracking were extracted from the video and downsampled to 30 Hz for later analysis. Audio was recorded at 10KHz. Flies were introduced gently into the chamber using an aspirator. Recordings were timed to be within 150 minutes of the behavioral incubator lights switching on to catch the morning activity peak. Recordings were stopped either after 30 minutes or after copulation, whichever came sooner. All flies were used; we did not use any criteria (e.g., if males sang during the first five minutes of the experiment or not) to drop fly sessions from analyses. In total, behavior was recorded and analyzed from 459 male flies; the number of flies per condition were as follows:

| LC type | LC4 | LC6 | LC9 | LC10a | LC10ad | LC10bc | LC10d | LC11 | LC12 | LC13 | LC15 | LC16 | LC17 |
| --- | --- | --- | --- | --- | --- | --- | --- | --- | --- | --- | --- | --- | --- |
| number of flies | 17 | 19 | 18 | 13 | 15 | 16 | 16 | 14 | 14 | 17 | 16 | 19 | 14 |

| LC type | LC18 | LC20 | LC21 | LC22 | LC24 | LC25 | LC26 | LC31 | LPLC1 | LPLC2 | control | total |
| --- | --- | --- | --- | --- | --- | --- | --- | --- | --- | --- | --- | --- |
| number of flies | 14 | 16 | 15 | 22 | 18 | 16 | 18 | 24 | 16 | 17 | 75 | 459 |

Joint positions for the male and female for every frame were tracked with a deep neural network trained for multi-animal pose tracking called SLEAP [28]. We used the default values for the parameters and proofread the resulting tracks to correct for errors. We estimated the presence of sine, Pfast, and Pslow song for every frame using a song segmenter on the audio signals recorded from the chamber’s microphones according to a previous study [66, 29].

From the tracked joint positions and song recordings, we extracted the following 6 behavioral variables of the male fly that represented his moment-to-moment behavior. 1) *Forward velocity* was the difference between the male’s current position minus his position one frame in his past; this difference in position was projected onto his heading direction (i.e., the vector from the male’s body position to his head position). 2) *Lateral velocity* was the same difference in position as computed for forward velocity except this difference was projected onto the direction orthogonal to his heading direction; rightward movements were taken as positive. 3) *Angular velocity* was the angle between the male’s current heading direction and the male’s heading direction one frame in the past; rightward turns were taken as positive, and angles were reported in visual degrees. 4) *Probability of sine song* was computed as a binary variable for each frame, where a value of 1 was reported if sine song was present during that frame, else 0 was reported. 5) *Probability of pulse-fast (Pfast) song* and 6) *probability of pulse-slow (Pslow) song* were computed in the same manner as that for the probability of sine song. These six behavioral output variables described the male’s movement (forward, lateral, and angular velocity) as well as his song production (probability of sine, Pfast, and Pslow song).

Often a male fly spends large periods of time without noticeable courtship of the female (e.g., the ‘whatever’ state as labeled in [67]). Predicting the male fly’s behavior from his visual input within these periods is difficult for any model. In addition, these time periods can make up a large enough fraction of the training data to bias models to output “do nothing” due to the imbalanced training data. To mitigate these effects, we devised a set of loose criteria to identify “courtship frames” in which the male is likely in the courtship state (e.g., chasing or singing to the female); we then only train and test on these courtship frames.

We devised the following four criteria to determine if a frame is a courtship frame:

1. The male and female distance (taken between the joint positions of their thoraxes) averaged over the time window is less than 5 mm.
2. The proportion of frames in which the male produced song (Pfast, Pslow, or sine) during the time window is greater than 0.1.
3. The angle of the female’s location from the male’s heading direction (with respect to the male’s head), averaged over the time window, is no more than 45 visual degrees.
4. The male is traveling at least 4.5 mm/s towards the female, averaged over the time window.

The time window was 20 s long, centered on the candidate frame. Only one criterion needed to be met to classify a frame as a courtship frame. Given these criteria, roughly 70% of all frames in control sessions were considered as courtship frames. Although silencing an LC type likely alters the amount of courtship during a session, we ensured that enough courtship frames were present for training the model. LC9-silenced males had the lowest percentage of courtship frames over the entire session at 42% (consistent with its high male-to-female distance, Fig. 1**e**, top panel); the average across LC types was roughly 70% and similar to that of the control sessions.

### Visual input reconstruction

To best mimic how a male fly transforms his retina’s visual input into behavior, we desired an image-computable model (i.e., one that takes as input an image rather than abstract variables determined by the experimenter, such as female size or male-to-female distance). We approximately reconstructed the male’s visual input based on the joint positions of both the male and female fly during courtship, as described in the following process. For each frame, we created a 64-pixel *×* 256-pixel grayscale image with a white background. Given the female rotation, size, and location (see below), we placed an image patch of a grayscale fictive female (composed of ellipses that represented the head, eyes, thorax, and tail of the female; no wings were included) occluding the white background. Because male flies perceive roughly 160 visual degrees on either side [68], we removed from the image the 40 visual degrees directly behind the male, leading to images with 64 *×* 228 pixels. Example input images are shown in Figure 1**f**, where the reconstructed female flies were colored and on gray background for illustrative purposes. Example videos of input image sequences are present in Supplementary Video 1 and Supplementary Video 2.

We computed the female’s rotation, size, and location in the following way. For *female rotation*, we computed the angle between the direction of the male head to female body and the direction of the female’s heading. A rotation angle of 0° indicates the female is facing away from the male, *±*180° indicates the female is facing towards the male, and -90°/+90° indicates the female is facing to the left/right of the male. We pre-processed a set of 360 image patches (25 *×* 25 pixels) that depicted a rotated female for each of 360 visual degrees. Given the computed rotation angle, we accessed the image patch corresponding to that rotation angle. For *female size*, we treated the female fly as a sphere (whose diameter matched the average size of a female fly, *∼*4 mm) and computed as size the visual angle between the two vectors of the male’s head position to the two outermost points on the sphere that maximize the visual angle (i.e., the two furthest points along the horizontal center line); this angle was normalized so that a size of 1 corresponded to 180 visual degrees. This size determined the width (and height, equal to the width) of the selected image patch to be placed into the 64 *×* 228-pixel image. Here, size indicates the size of the image patch, not the actual size of the fictive female (which may vary because a female facing away is smaller than a female facing to the left or right). We measured the approximate visual degrees of a fictive female’s body with a size of 1.0 facing away from the male in the center as 65 visual degrees. For *female position*, we computed the visual angle between the male’s heading direction and the direction between the male’s head and the female’s body position. We normalized this angle such that a position of 0 is directly in front of the male, a position of either -1 or 1 is directly behind the male fly, and a position of -0.5/+0.5 is 90 visual degrees to the left/right. We then used this position to place the image patch (with its chosen rotation and size) at a certain pixel location along the horizontal centerline of the image. Because the male and female flies did not have room to fly in the experimental chamber, we assumed that only the female’s lateral position (and not vertical position) could change.

### Description of 1-to-1 network

We designed our 1-to-1 network to predict the male fly’s behavior (i.e., movement and song production) only from his visual input. Although the male can use other sensory modalities such as olfaction or proprioception to detect the female, we chose to focus solely on visual inputs because 1) the male relies primarily on his visual feedback for courtship [9, 69], 2) we wanted the model to have a representation solely based on vision to match the representations of visual LC neurons, and 3) it is unclear to what extent the fly can access its past movement history in the form of proprioception. Interestingly, past movement information is predictive of forward and lateral velocity but not angular velocity (Ext. Data Fig. 3), suggesting turning is better predicted by visual input than proprioception.

Our choice to use a feedforward DNN to model real neurons is motivated by past success in using DNNs to link object recognition and the primate visual cortex [70], navigation and grid cells [71], and reaching and motor cortex [2]. Recent work has begun to use DNNs to model fruit fly vision [16, 39]. However, using a DNN comes with the caveat that we have made simplifying assumptions about the complexity of the true computations in the brain. Our 1-to-1 network has no recurrent connections despite recurrence found in the optic lobe and central brain. We suspect that adding more biological constraints, such as recurrence, delays, excitation/inhibition, and connectome information [18, 45], will allow knockout training to better identify our model’s one-to-one mapping given enough training data. Still, as we more strongly constrain the model, we risk losing the expressivity needed for accurate prediction. A feedforward DNN unconstrained by biological realism has key advantages: It is easily trained with backpropagation; it is highly-expressive and can fit complex mappings; and software packages like Keras [72] are readily available. The only part of the 1-to-1 network that we analyzed was the LC bottleneck layer—we did not analyze other subnetworks (i.e., the vision and decision networks, see below) and treated them as “black box” mappings.

The 1-to-1 network comprised three parts: a vision network, an LC bottleneck, and a decision network (Fig. 1**a**). Hyperparameters, such as the number of filters in each layer, the number of layers, and the types of layers were chosen based on prediction performance assessed on a validation set of the control sessions separate from the test set. Unless specified, each convolutional or dense layer was followed by a batchnorm operation [73] and a relu activation function. The 1-to-1 network took as input the images of the 10 most recent time frames (corresponding to *∼*300 ms)—longer input sequences did not lead to an improvement in predicting behavior. Each grayscale image was 64 *×* 228 pixels (with values between 0 and 255) depicting a fictive female fly on a white background (see “Visual input reconstruction” above). Before being fed into the network, the input was first re-centered by subtracting 255 from each pixel intensity to ensure the background pixels had values of 0. The model’s output was 6 behavioral variables of the male fly: forward velocity, lateral velocity, angular velocity, probability of sine song, probability of Pfast song, and probability of Pslow song (see “Courtship experiments” above).

#### Vision network

The first layer of the vision network was spatial convolutions with 32 filters (kernel size 3 *×* 3) and a downsampling stride of 2). The second and third layers were identical to the first except with separable 2-d convolutions [74]. The final layer was a two-stage linear mapping [75] which first spatially pools its input of activity maps and then linearly combines the pooled outputs across channels into 16 embedding variables; pooling the spatial inputs in this manner greatly reduced the number of parameters for this layer. Batchnorm and relus did not follow this two-stage layer. The vision network processed each of the 10 input images separately; in other words, the vision network’s weights were shared across time frames (i.e., a 1-d convolution in time). Allowing for 3-d convolutions of the visual inputs (i.e., 3-d kernels for the two spatial dimensions and the third time dimension) did not improve prediction performance (Ext. Data Fig. 3), likely because of the increase in the number of parameters. For simplicity, the vision network’s input was the entire image (i.e., the entire visual field); we did not include two retinae. We found that incorporating two retinae into the model, while more biologically-plausible, made it more difficult to interpret the tuning of each LC neuron type. For example, for a two-retinae model, it is difficult to determine if differences in tuning for two model units of the same LC type but in different retinae are true differences in real LC types or instead differences due to overfitting between the two retinal vision networks. The 1-to-1 network avoids this discrepancy through the simplifying assumption that each LC type has a similar tuning across both retinae.

#### LC bottleneck

The next component of the DNN model was the LC bottleneck, which received 10 16-dimensional embedding vectors corresponding to the past 10 time frames. These embedding vectors were passed through a dense layer with 64 filters followed by another dense layer with number of filters equal to the number of silenced LC types (23 in total). We call the 23-d output of this layer as the “LC bottleneck”. Each model LC unit represents the summed activity of all neurons of the same LC type (i.e., projecting to the same glomerulus), which makes it easy to compare to calcium imaging recordings of LC neurons which track the overall activity level of a single optic glomerulus. Another possible choice was to treat the responses of neurons for the same LC type as the outputs of a convolutional layer (where the neurons’ receptive fields tile the visual field) and then sum this output as the predicted overall activity of the optic glomerulus. This is essentially the same as our choice for the LC bottleneck layer, which integrates over the spatial information of the vision network (itself made up of convolutional layers); the key difference between the two is that here we do not assume that the overall activity of the optic glomerulus results from a simple sum over neurons but instead from a possibly nonlinear readout that is relevant to behavior. We found that adding additional unperturbed “slack” model LC units to match the total number of LC types (e.g., 45 model LC units instead of 23 units) did not improve prediction performance; in the extreme case, adding a large number slack variables encourages the network to ignore the “unreliable” knocked-out units in favor of predicting shared behavior across silenced and control sessions (i.e., similar to training without knockout). For two perturbations (LC10ad and LC10bc), we only had genetic lines to silence two LC neuron types together. For simplicity, we corresponded each of these with its own model LC unit, which represented the summed activity of all neurons from both types (e.g., LC10a and LC10d for LC10ad). Because the LC bottleneck reads from all 10 past time frames, each model LC unit integrates information over time (e.g., for motion detection). Additionally, the model LC responses are guaranteed to be nonnegative because of the relu activation functions.

#### Decision network

The decision network took as input the activations of the 23 LC bottleneck units and comprised 3 dense layers, where each layer had 128 filters. The decision network predicted the movement output variables (forward velocity, lateral velocity, and angular velocity) each with a linear mapping and the song production variables (probability of sine, Pfast, and Pslow song) each with a linear mapping followed by a sigmoid activation function.

### Knockout training

We sought a one-to-one mapping between the model’s 23 LC units in its bottleneck and the 23 LC neuron types in our silencing experiments (Fig. 1**a**). To identify this mapping, we devised knockout training. We first describe the high-level training procedure and then give details about the optimization. For a randomly initialized 1-to-1 network, we arbitrarily assigned model LC units to real LC types (i.e., in alphabetical order of the LC names). For each training sample, we “knocked out” (i.e., set to 0 via a mask) the model LC unit that corresponded to the silenced LC type; no model units were silenced for control data (Fig. 1**b**). This is similar to dropout training [40] except that hidden units were purposefully—not randomly—chosen. The intuition behind knockout training is that the remaining unperturbed model LC units must encode enough information or “pick up the slack” to predict the silenced behavior; any extra information will not be encoded in the unperturbed units (as the back-propagated error would not contain this information). For example, let us assume that LPLC1 solely encodes female size and contributes strongly to forward velocity. To predict LPLC1-silenced behavior (which would not rely on female size), the other model LC units would need only to encode other features of the fictive female (e.g., her position or rotation). In fact, any other model LC unit encoding female size would hurt prediction because LPLC1-silenced behavior does not depend on it. Another view of knockout training is that we constrain the model not only to predict behavior but also to predict behavior with certain constraints on what internal representations the model may use. These constraints are set by the perturbations (e.g., genetic silencing) we use in our experiments.

The optimization details are as follows. The model was trained end-to-end using stochastic gradient descent with learning rate 1e-3 and momentum 0.7. Each training batch had 288 samples, where each sample was a sequence of 10 images and 6 output values. Each batch was balanced across LC types (24 in total including control), where each LC type had 12 samples. The batch was also balanced for types of song (sine song, pulse song, or no song), as different flies sang different amounts of song. The model treated different flies for the same silenced LC type as the same to capture overall trends of an “average” fly. We z-scored the movement behavioral variables (forward, lateral, and angular velocity) based on the mean and standard deviation of the control data in order to have similarly-sized gradients from each output variable. The loss functions were mean squared error for forward, lat-eral, and angular velocity and binary cross-entropy for the probabilities of sine, Pfast, and Pslow song. The model instantiation and optimization was coded in Keras [72] on top of Tensorflow [76]; we used the default random initialization parameters to initialize weights. We stopped training when prediction performance for forward velocity (evaluated on a validation set, see below) began to decrease (i.e., early stopping).

#### Training and test data

After identifying courtship frames (see ‘Courtship experiments’ above), we split these frames into train, validation, and test sets. To form a test set for a given LC type (or control sessions), we randomly selected 3 second windows across all flies until we had 15 minutes of data (27,000 frames). Selecting contiguous windows ensured that no frame in the visual input of the test data overlapped with any training frames. For control sessions, after selecting the test set, we also randomly sampled from the remaining frames to form a validation set (27,000 frames) in the same way as we did for the test set; the validation set was used for hyperparameter choices and early stopping. All remaining frames were used for training. Because the number of control sessions greatly outnumbered LC-silenced sessions, we randomly sampled a subset of frames over all sessions for a given LC type and stopped when we reached 600,000 frames (*∼*5.5 hours). In total, our training set had *∼*11.6 million training samples. To account for the observation that flies tend to prefer to walk along the edge of the chamber in either a clockwise or counter-clockwise manner—biasing lateral and angular velocities to one direction—we augmented the training set by flipping the visual input from left to right and correspondingly changing the sign of the lateral and angular velocities; each training sample had a random 50% chance of being flipped.

#### Dropout and no knockout training

For comparison to knockout training, we considered three networks with the same architecture as the 1-to-1 network but with other training procedures (Ext. Data Fig. 3**b**). First is the un-trained network for which no training is performed (i.e., all parameters remain at their randomized initial values). Second, we performed a version of dropout training [40] by setting to 0 a randomly-chosen model LC unit for each training frame independent of the frame’s silenced LC type; no model LC unit’s values are set to 0 for frames from control sessions. This training procedure “knocks out” the exact same number of units as that of knockout training. Third, we consider training a network *without* knocking out any model LC units. A key difference between these three networks and the 1-to-1 network is that none of these networks were given any information about which LC type was silenced for a training sample; they can only identify computations shared among sessions with differently silenced LC types as well as control sessions. This helped us to ground the prediction performance of knockout training when predicting moment-to-moment behavior (Ext. Data Fig. 3 and Ext. Data Fig. 4) and real LC responses (Fig. 2**e**) as well as consistency in training (see below).

#### Consistency across different training runs

Because DNNs are optimized via stochastic gradient descent, the training procedure of a DNN is not deterministic; different random intializations and different orderings of the training data may lead to DNNs with different prediction performances. To assess whether the 1-to-1 network is consistent across training runs, we trained 10 runs of the 1-to-1 network with different random initializations and different random orderings of training samples. We chose 10 runs as a compromise between having enough runs to estimate consistency while remaining computationally feasible (each 1-to-1 network took 1 week to train). For comparison, we also trained 10 networks either with dropout training or without knockout training (see section above) as well as 10 untrained networks. For a fair comparison across training procedures (knockout, dropout, without knockout, and untrained), each run had the same parameter intialization and ordering of training samples. We compared the 1-to-1 network to these three networks by assessing prediction performance of moment-to-moment behavior (Ext. Data Fig. 3**d**), overall mean changes to behavior across silenced LC types (Ext. Data Fig. 4), consistency both in behavioral predictions (Ext. Data Fig. 5) and neural predictions (Ext. Data Fig. 6), prediction performance of real LC responses for a one-to-one mapping (Fig. 2**e** and Ext. Data Fig. 8**b**) and prediction performance of real LC responses for a fitted linear mapping (Ext. Data Fig. 9). We opted to investigate the inner workings of a single 1-to-1 network in Figures 3 and 4 both for simplicity and because some analyses can only be performed on a single network (e.g., the cumulative ablation experiments in Fig. 4). Different runs of the 1-to-1 networks had some differences in their predictions (Ext. Data Fig. 5 and Ext. Data Fig. 6), but the overall conclusion that the LC bottleneck in the 1-to-1 network formed a distributed population code remained true over all runs. For our analyses in Figures 3ãnd 4, we chose the 1-to-1 network that had the best prediction for both behavior and neural responses (model 1 in Ext. Data Fig. 3**d** and in Ext. Data Fig. 8**b**).

### Two-photon calcium imaging

We recorded LC responses of a head-fixed male fly using a custom-built two-photon microscope with a 40x objective and a two-photon laser (Coherent) tuned to 920 nm for imaging of GCaMP6f. A 562 nm dichroic split the emission light into red and green channels, which were then passed through a red 545-604 nm and green 485- 555 nm bandpass filter respectively. We recorded the imaging data from the green channel with a single plane at 50 Hz. Before head-fixation, the male’s cuticle above the brain was surgically removed, and the brain was perfused with an extracellular saline composition. The male’s temperature was controlled at 30°C by flowing saline through a Peltier device and measured via a water bath with a thermistor (BioscienceTools TC2-80-150). We targeted LC neuron types LC6, LC11, LC12, LC15, and LC17 (Fig. 2**a**) for their proximity to the surface (and thus better imaging signal), prior knowledge about their responses from previous studies [30, 31, 32], changes to behavior when silenced (Fig. 1**e** and Ext. Data Fig. 1), and their corresponding model LC responses to stimuli (Fig. 3**b** and **g**). Genotypes for the different LC lines are in Supplemental Table 2.

Each head-fixed male fly walked on a freely-moveable, air-supported ball and viewed a translucent projection screen placed in the right visual hemifield (matching our recording location in the right hemisphere). The flat screen was slanted 40 visual degrees from the heading direction of the fly and perpendicular to the axis along the direction between the fly’s head and the center of the screen (with a distance of 9 cm between the two). An LED-projector (DLP Lightcrafter LC3000-G2-PRO) with a Semrock FF01-468/SP-25-STR filter projected stimulus sequences onto the back of the screen at a frame rate of 180 fps. A neutral density filter of optical density 1.3 was added to the output of the projector to reduce light intensity. The stimulus sequences (described below) comprised a moving spot and a fictive female that varied her size, position, and rotation.

We recorded a number of sessions for each targeted LC: LC6 (5 flies), LC11 (5 flies), LC12 (6 flies), LC15 (4 flies), and LC17 (5 flies). We imaged each glomerulus at the broadest cross-section, typically at the mid-point, given that we positioned the head of the fly to be flat (tilted down 90°, with the eyes pointing down). We hand-selected ROIs that encompassed the shape of the glomerulus within the 2-d cross-section. We computed dF/F_0_ for these targeted ROIs using a baseline ROI for F_0_ that had no discernible response and was far from targeted ROIs. For each LC and stimulus sequence, we concatenated repeats across flies. To remove effects due to adaptation across repeats and differences among flies, we de-trended responses by taking the z-score across time for each repeat; we then scaled and re-centered each repeat’s z-scored trace by the standard deviation and mean across all the original repeats (i.e., the original and de-noised repeats had the same overall mean and standard deviation). To test whether an LC was responsive to a stimulus sequence or not, we computed a metric akin to a signal-to-noise ratio for each combination of LC type and stimulus sequence in the following way. For a single run, we split the repeats into two separate groups (same number of repeats per group) and computed the repeat-averaged response for each group. We then computed the *R*^2^ between the two repeat-averaged responses by computing the Pearson correlation over time and squaring it. We performed 50 runs with random split groups of repeats to establish a distribution of *R*^2^’s. We compared this distribution to a null distribution of *R*^2^’s that retained the timecourses of the responses but none of the time-varying relationships among repeats. To compute this null distribution, we sampled 50 runs of split groups (same number of repeats as the actual split groups) from the set of repeats for all stimulus sequences; in addition, the responses for each repeat were randomly reversed in time or flipped in sign, breaking any possible co-variation across time among repeats. For each combination of LC type and stimulus, we computed the sensi-tivity *d*′ [77] between the actual *R*^2^ distribution and the null *R*^2^ distribution. We designated a threshold *d*′ *>* 1 to indicate that an LC was responsive for a given stimulus sequence (i.e., we had a reliable estimate of the repeat-averaged response). After this procedure, a total of 27 combinations of stimulus sequence and LC type out of a possible 45 combinations remained (Ext. Data Fig. 8).

We considered two types of stimulus sequences: a moving spot and a moving fictive female. The moving spot (black on isoluminant gray background) had three different stimulus sequences (Fig. 2**b** and **c**). The first stimulus sequence was a black spot with fixed diameter of 20° that moved from the left to right with a velocity chosen from candidate velocities {1, 2, 5, 10, 20, 40, 80} °/s; each sequence lasted 2 seconds. The second stimulus sequence was a spot that loomed from a starting diameter of 80° to a final diameter of 180° according to the formula *θ*(*t*) = *−*2 tan*^−^*^1^(*−r/v ·* 1*/t*), where *r/v* is the radius-to-speed ratio with units in ms and *t* is the time (in ms) until the object reaches its maximum diameter (i.e., *t* = *t*_final_ - *t*_current_) [20]. A larger *r/v* establishes a slower object loom. We presented different loom speed ratios chosen from candidate *r/v ∈* {10, 20, 40, 80} ms. Once a diameter of 180° was reached, the diameter remained constant. The third stimulus sequence was a spot that linearly increased its size from a starting diameter of 10° according to the formula *θ* = 10 + *v · t*, where *v* is the angular velocity (in °/s) and *t* is the time from stimulus onset (in seconds). The final diameter of the enlarging spot for each velocity (30°, 50°, 90°, or 90°, respectively) was determined based on the chosen angular velocity *v ∈ {*10, 20, 40, 80*}* °/s. Once a diameter of 90° was reached, the diameter remained constant.

The second type of stimulus sequence was a fictive female varying her size, position, and rotation. The fictive female was generated in the same manner as that for the input of the 1-to-1 network (see Methods section “Visual input reconstruction”). We took the angular size of the fictive female (65 visual degrees for a size of 1.0, where the female faces away from the male at the center of the image) and used it to set the angular size of the fictive female on the projection screen. We considered three kinds of fictive female stimulus sequences with 9 different sequences in total (Supplementary Video 1 and Ext. Data Fig. 8); we first describe them at a high level and then separately in more detail. The first kind consisted of sequences in which the female varied only one visual parameter (e.g., size) while the other two parameters remained fixed (e.g., position and rotation); we varied this parameter with three different speeds. Second, we generated sequences that optimized a model output variable (e.g., maximizing or minimizing forward velocity). Third, we used a natural image sequence taken from a courtship session. Each stimulus sequence lasted for 10 seconds (300 frames).

Details of the fictive female sequences are as follows:

- *vary female position*: The female varied only her lateral position (with a fixed size of 0.8 and a rotation angle of 0° facing away from the male) from left-to-right (75 frames) then right-to-left (75 frames). Positions were linearly sampled in equal intervals between the range of -0.1 and 0.5. This range of positions was biased to the rightside of the visual field to account for the fact that the projection screen was oriented in the male’s right visual hemifield. After the intial pass of left-to-right and right-to-left (150 frames total), we repeated this same pass two more times with shorter periods (100 frames and 50 frames in total, respectively), interpolating positions in the same manner as the initial pass.
- *vary female size*: The same generation procedure as for ‘vary female position’ except that instead of position, we varied female size from 0.4 to 0.9 (sampled in equal intervals) with a fixed position of 0.25 and a rotation angle of 0° facing away from the male.
- *vary female rotation*: The same generation procedure as for ‘vary female position’ except that instead of position, we varied the female rotation angle from -180° to 180° (sampled in equal intervals) with a fixed position of 0.25 and a fixed size of 0.8.
- *optimize for forward velocity*: We optimized a 10-second stimulus sequence in which female size, position, and rotation were chosen to maximize the 1-to-1 network’s output of forward velocity for 5 seconds and then minimize forward velocity for 5 seconds. In a greedy manner, the next image in the sequence was chosen from candidate images to maximize the objective. We confirmed that this approach did yield large variations in the model’s output. To ensure smooth transitions, the candidate images were images “nearby” in parameter space (i.e., if the current size was 0.8, we would only consider candidate images with sizes in the range of 0.75 to 0.85). Images were not allowed to be the same in consecutive frames and had to have a female size greater than 0.3 and a female position between -0.1 and 0.5.
- *optimize for lateral velocity*: The same generation procedure as for ‘optimize for forward velocity’ except that we optimized for the model output of lateral velocity. In this case, maximizing/minimizing lateral velocity is akin to asking the model to output the action of moving to the right/left.
- *optimize for angular velocity*: The same generation procedure as for ‘optimize for forward velocity’ except that we optimized for the model output of angular velocity. In this case, maximizing/minimizing angular velocity is akin to asking the model to output the action of turning to the right/left.
- *optimize for forward velocity with fixed position*: The same generation procedure as for ‘optimize for forward velocity’ except that we limited female position *p* to be within the tight range of 0.225 *< p <* 0.275. This ensured that most changes of the female stemmed from changes in either female size or rotation, not position.
- *optimize for lateral velocity with multiple transitions*: The same generation procedure as for ‘optimize for lateral velocity’ except that we had four optimization periods: maximize for 2.5 seconds, minimize for 2.5 seconds, maximize for 2.5 seconds, and minimize for 2.5 seconds.
- *natural stimulus sequence*: A 10-second stimulus sequence taken from a real courtship session. This sequence was chosen to ensure large variation in the visual parameters and that the female fly was mostly in the right visual field between positions -0.1 and 0.5 (as stimuli were presented to the right visual hemifield of the male fly).

For each recording session, we presented the stimuli in the following way. For the moving spot stimuli, each stimulus sequence was preceded by 400 ms of a blank, isoluminant gray screen. For the fictive female stimuli, a stimulus sequence of the same kind (e.g., ‘vary female size’) was presented in three consecutive repeats for a total of 30 seconds; this stimulus block was preceded by 400 ms of a blank, isoluminant gray screen. All stimulus sequences (both moving spot and the fictive female) were presented one time each in a random ordering. Another round (with the same ordering) was presented if time allowed; usually, we presented 3 to 4 stimulus rounds before an experiment concluded. This typically provided 9 or more repeats per stimulus sequence per fly.

### Predicting real neural responses

To obtain the model predictions for the artificial moving spot stimuli (Fig. 2**b** and **c**, bottom rows), we generated a fictive female facing away from the male whose size and position matched that of the moving spot. This was done to prevent any artifacts from presenting a stimulus (e.g., a high-contrast moving spot) on which the model had not been trained, as the model only observed a fictive female. We matched the angular size of the fictive female to that of the presented stimulus by using the measured conversion factor of 65 visual degrees for a fictive female size of 1.0. For the stimulus of the moving spot with varying speed (Fig. 2**b**), the fictive female translated from left to right (i.e., same as the stimuli presented to the male fly). Because the 1-to-1 network’s responses could remain constant and not return to 0 for different static stimuli (i.e., no adaptation mechanism), we added a simple adaptation mechanism to the model’s responses such that if responses were the same for consecutive frames, the second frame’s response would return to its initial baseline response with a decay rate of 0.1. To obtain model predictions for the fictive female stimuli (Fig. 2**d**-**e**), we input the same stimululus sequences presented to the fly except that we changed the grayscale background to white (to match the training images).

To evaluate the extent to which the 1-to-1 network predicted the repeat-averaged LC responses for each stimulus sequence of the moving fictive female, we sought a prediction performance metric (e.g., *R*^2^) that accounted for the fact that our estimates of the repeat-averaged responses were noisy. Any metric not accounting for this noise would undervalue the true prediction performance (i.e., the prediction performance between a model and a repeat-averaged response with an infinite number of repeats). To measure prediction performance, we chose a noise-corrected *R*^2^ metric recently proposed [78] that precisely accounts for noise across repeats and is unbiased at estimating the ground-truth normalized *R*^2^. A noise-corrected *R*^2^ = 1 indicates that our model perfectly predicts the ground-truth repeat-averaged responses up to the amount of noise across repeats. We note that our noise-corrected *R*^2^ metric accounts for differences in mean, standard deviation, and sign between model and real responses, as these differences do not represent the information content of the responses.

We computed this noise-corrected *R*^2^ between the 1-to-1 network and real responses for each LC type and stimulus sequence (Fig. 2**e**) for which the LC was responsive (i.e., *d^i^ >* 1). Importantly, the 1-to-1 network never had access to any neural data in its training; instead, for a given LC type, we directly took the response of the corresponding model LC unit as the 1-to-1 network’s predicted response. This is a stronger criterion than typical evaluations of DNN models and neural activity, where a linear mapping from DNN features (*∼*10,000 feature variables) to neural responses is fit [1]. To account for the smoothness of real responses due to the imaging of calcium dynamics, we causally smoothed the predicted responses with a linear filter. We fit the weights of the linear filter (filtering the 10 past frames) along with the relu’s offset parameter (accounting for trivial mismatches due to differences in thresholding) to the real responses. This fitting only used responses of one model LC unit, keeping in place the one-to-one mapping; we also fit a linear mapping using all model LC units (Ext. Data Fig. 9). We performed the same smoothing procedure not only for the 1-to-1 network but also for an untrained network, a network trained with dropout training, and a network trained without knockout (see “Knockout training” above).

This procedure was only performed for predicted responses in Figure 2**d** and **e** and Extended Data Figure 8. For analyzing response magnitudes (Fig. 2**f** and Ext. Data Fig. 10), the responses came directly from model LC units (i.e., no smoothing or fitting of the relu’s offset was performed).

### Analyzing model LC responses to visual input

To better understand how each model LC unit responds to the visual input, we systematically varied female size, position, and rotation to generate a large bank of stimulus sequences. We input these stimulus sequences into the 1-to-1 network and formed heatmaps out of the model LC responses (Fig. 3**c**ãnd **d**). For each input stimulus sequence, each of its 10 images was a repeat of the same image of a fictive female with a given size, lateral position, and rotation angle (i.e., the fictive female remained frozen over time for each 10-frame input sequence). Across stimulus sequences, we varied female size (50 values linearly interpolated between 0.3 to 1.1), lateral position (50 values linearly interpolated between -1 to 1), and rotation angle (50 values linearly interpolated between -180 and 180 visual degrees), resulting in 50 *×* 50 *×* 50 = 125, 000 different stimulus sequences that enumerated all possible combinations. To understand the extent to which each visual parameter contributed to a model LC unit’s response, we decomposed the total response variance into different components (Fig. 3**d**) [43]. The first three components represent the variance of the marginal response to each of the 3 visual parameters (which we had independently varied). We computed these marginalized variances by 1) taking the mean response for each value of a given visual parameter by averaging the other two parameters over all stimulus sequences and 2) taking the variance of this mean response over values of the marginalized parameter (50 values in total). Any remaining variance (subtracting the three marginalized variances from the total response variance) represents response variance arising from interactions among the three visual parameters (e.g., the model LC response depends on female size but only if the female is in the center and faces away from the male, see Fig. 4**d**, ‘LC10a’). Because the 1-to-1 network was deterministic, no response variance was attributed to noise across repeats (unlike trial-to-trial variability observed in the responses of real neurons).

While analyzing the model LC responses to a large bank of static stimuli is helpful to understand LC tuning (Fig. 3**c**-**e**), we may miss important relationships between the features of the visual input and model LC responses without considering dynamics (e.g., the speed at which female size changes). To account for these other temporal features, we devised three dynamic stimulus sequences that varied in time for roughly 10 seconds each (Fig. 3**f** and Supplementary Video 2); these stimuli were similar to a subset of stimuli we presented to real male flies (see Methods section “Two-photon functional imaging”). For each stimulus sequence, we varied one visual parameter while the other two remained fixed at nominal values chosen based on natural sequence statistics.

The first 2.5 seconds of each stimulus were the following:

1. *vary female size*: linearly increase from 0.5 to 0.9 with fixed position = 0 and rotation = 0°
2. *vary female position*: linearly increase from -0.25 to 0.25 with fixed size = 0.8 and rotation = 0°
3. 3. *vary female rotation*: linearly increase from -45° to 45° with fixed size = 0.8 and position = 0

The next 2.5 seconds were the same as the first 2.5 seconds except reversed in time (e.g., if the female increased in size the first 2.5 seconds, then the female decreased in size at the same speed for the next 2.5 seconds). Thus, the first 5 seconds was one period in which the female increased and decreased one parameter. The stimulus sequence contained 4 repeats of this period with different lengths (i.e., different speeds): 5, 3.33, 1.66, and 0.66 seconds (corresponding to 150, 100, 50, and 10 time frames, respectively). We passed these stimulus sequences as input into the 1-to-1 network (i.e., for each time frame, the 10 most recent images were passed into the model) and collected the model LC responses over time. We directly computed the squared correlation *R*^2^ between each model LC unit’s responses and the visual parameters (and features derived from the visual parameters, such as speed) for all three stimulus sequences (Fig. 3**g**). Velocity and speed were computed by taking the difference of the visual parameter between two consecutive time frames.

### Analyzing how model LCs contribute to behavior

Because the 1-to-1 network identifies a one-to-one mapping, the model predicts not only the response of an LC neuron but also how that LC neuron causally relates to behavior. We wondered to what extent each model LC unit causally contributed to each behavioral output variable. We designed an ablation approach (termed the CumuLa-tive Inactivation Procedure or CLIP) to identify which model LCs contributed the most to each behavioral output. The first step in CLIP is to inactivate each model LC unit individually by setting a model LC’s activity value for all time frames to a constant value (chosen to be the mean activity value across all frames). We then test to what extent the 1-to-1 network with the inactivated model LC unit predicts the behavioral output of heldout test data from control flies (from the test set). We choose the model LC unit that, once inactivated, leads to the *least* drop in prediction performance (i.e., the model LC unit that contributes the least to the behavioral output). We then iteratively repeat this step, keeping all previously-inactivated model LC units still inactivated. In this way, we greedily ablate model LC units until only one model LC unit remains. After performing CLIP, we obtain an ordering of model LC units from weakest to strongest contributor of a particular behavioral output (Fig. 4**b** and **c**). We then use this ordering (and prediction performance) to infer which model LC units contribute to which behavioral outputs. We performed CLIP to predict heldout behavior from control flies (Fig. 4**c**) as well as the model output to simple, dynamic stimulus sequences (Fig. 4**d**-**f**). For the dynamic stimulus sequences (where we did not have real behavioral data), we used the model output when no silencing occurred as ground truth behavior. Because different behavioral outputs had different prediction performances (Ext. Data Fig. 3), we normalized each model LC unit’s change in performance by the maximum change in performance (i.e., prediction performance for no inactivation minus that of inactivating all model LC units); for model LC units for which inactivation led to an increase in performance due to overfitting (Ext. Data Fig. 14), we clipped their change in performance to be 1.

### Connectome analysis

To obtain the pre-and postsynaptic partners of LC and LPLC neuron types, we leveraged the recently-released FlyWire connectome of an adult female *Drosophila* [45, 46], for which optic lobe intrinsic neurons were recently typed [17]. We downloaded the synaptic connection matrix at https://flywire.ai/ of the public release version 630. We isolated the following 35 LC and LPLC types: LC4, LC6, LC9, LC10a, LC10, LC10c, LC11, LC12, LC13, LC15, LC16, LC17, LC18, LC21, LC22, LC24, LC25, LC26, LC27, LC29, LC31, LC33a, LC34, LC35, LC36, LC37a, LC40a, LC41, LC43, LC45a, LC46a, LC46b, LPLC1, LPLC2, and LPLC4. We did not consider LC10d and LC20, which were silenced but not yet identified in the FlyWire connectome, and we do not consider the combined lines LC10ad and LC10bc. We summed the number of synaptic connections across all neurons of the same type that were either inputs or outputs of one of the LC and LPLC neuron types. We denoted a connection (Fig. 5**b**, tick lines) if at least 5 synaptic connections existed between an LC/LPLC neuron type and another neuron type; we chose 5 synapses to account for the fact that some neuron types had 1-2 neurons in total. Previous studies typically set a threshold of 50 synapses for a group of neuron types (e.g., a connection exists if 50 synapses are present between all LC4 neurons and all DN neurons, regardless of the specific DN cell type) [10, 41, 79]; instead, we do not group neuron types in order to have finer granularity. We identified 417 presynaptic partners and 660 postsynaptic partners (each partner being one neuron type).

### Statistical analysis

Unless otherwise stated, all statistical hypothesis testing was conducted with permutation tests, which do not assume any parametric form of the underlying probability distributions of the sample. All tests were two-sided and non-paired, unless otherwise noted. Each test was performed with 1,000 runs, where *p <* 0.001 indicates the highest signficance achievable given the number of runs performed. When comparing changes in behavior due to genetic silencing versus control flies (Fig. 1**e**), we accounted for multiple hypothesis testing by correcting the false discovery rate (FDR) with the Benjamini-Hochberg procedure with *α*=0.05. Paired permutation tests were performed when comparing prediction performance between models (Fig. 2**e**) for which paired samples were randomly permuted with one another. Error bars of the response traces in Figure 2**b**-**d** were 90% bootstrapped confidence intervals of the means, computed by randomly sampling repeats with replacement. No statistical methods were used to predetermine sample sizes, but our sample sizes are similar to those of previous studies [e.g., 10, 11, 30, 31]. Experimenters were not blinded to the conditions of the experiments during data collection and analysis.

## Acknowledgments.

We thank R. Pang and M. Aragon for comments on the manuscript and E. Ireland, K. Thieringer, and Y. Gao for assistance with data collection and processing. We thank A. Nern and G. Rubin for sharing the LC31 split-GAL4 lines ahead of publication. We thank A. Matsliah, S.-C. Yu, S. Seung, and the FlyWire team for sharing information on optic lobe cell types ahead of publication. We thank Tom Clandinin for sharing LC response data. This work was supported by a C.V. Starr Fellowship to B.R.C., a Simons Collaboration on the Global Brain Postdoctoral Fellowship to A.J.C., NIH grant K99-EY032549 to M.H.T., the Simons Collaboration on the Global Brain Investogator Awards and NIH BRAIN Initiative Award (R01 NS104899) to M.M. and J.W.P., and an HHMI Faculty Scholar Award and NINDS R35 Research Program Award to M.M.

## Data availability

Data are available at https://doi.org/10.34770/rmry-cs38.

## Code availability

The code for extracting fly body positions (SLEAP) is available at https://sleap.ai/. Song segmentation was performed with code found at https://github.com/murthylab/MurthyLab_FlySongSegmenter. Model weights, example stimuli, and code are available at https://github.com/murthylab/one2one-mapping. The FlyWire connectome is available at https://flywire.ai/.

## Author contributions

B.R.C., A.J.C., J.W.P., and M.M. conceived of and designed the study. A.J.C. designed and performed the silencing experiments. N.R. and M.H.T. designed and performed the imaging experiments. B.R.C. analyzed the imaging data. B.R.C. and A.J.C. designed the model; B.R.C. trained and analyzed the model. B.R.C. analyzed the connectome data. B.R.C. and M.M. wrote the manuscript with input from J.W.P., A.J.C., N.R., and M.H.T.

## Extended Data Figures

**Extended Data Figure 1:**
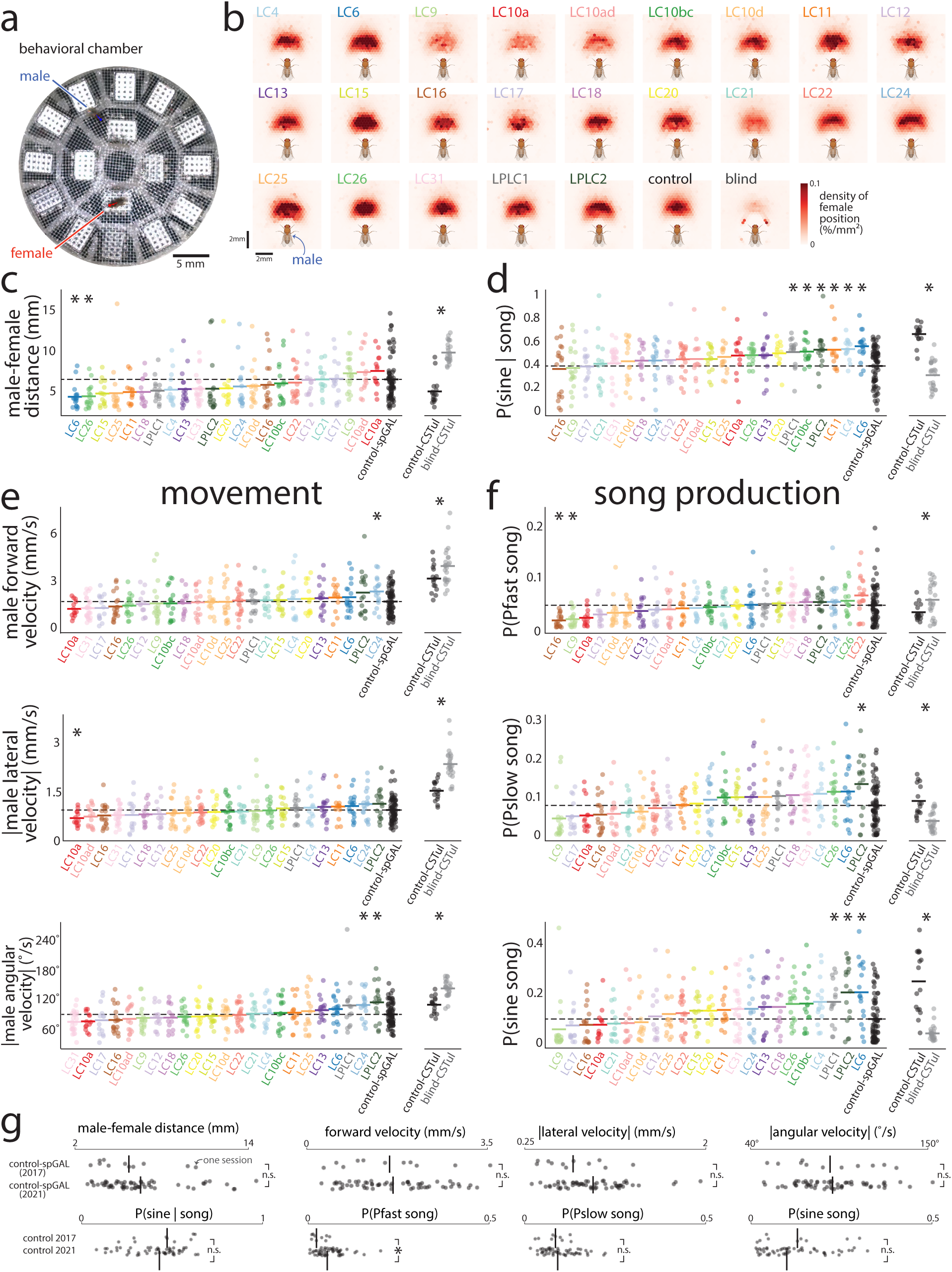
Different changes in behavior when silencing different LC neuron types of the male’s visual system. The main finding is that no single LC type showed a substantial change relative to control compared to the change observed between blind and control flies— suggesting no single LC type is the sole contributor to courtship behaviors. **a.** Image of circular behavioral chamber used to the positions of a male (blue) and female (red) fruit flies during courtship. Joint positions for each frame were identified with the behavioral tracking software SLEAP [28]. Audio waveforms of song were detected with 16 microphones tiling the chamber (white boxes). **b.** Density of female position relative to the male’s egocentric view, conditioned on which LC type was silenced in the male as well as control-spGAL (‘control’) and blind-CSTul (‘blind’) males (multiple sessions per heatmap). Silencing any single LC neuron type did not extinguish courtship chasing (compare LC-silenced heatmaps to that of blind males); however, silencing some LC types did lead to a noticeable decrease in the amount of time females were positioned in front of the male versus control sessions (e.g., compare LC9, LC10a, LC10ad, and LC21 to control). **c.** Male-female distance averaged across the entire session for each silenced LC type (reproduced from Fig. 1**e**, top panel). Each dot is for one session; lines denote means and dashed line denotes the mean for control sessions. Statistically significant changes from control flies are indicated by an asterisk (*p <* 0.05, permutation test, corrected for the false discovery rate of multiple hypothesis testing by the Benjamini-Hochberg procedure). We also considered changes to behavior between control and blind male flies in CSTul flies (right, data from [9, 67] recorded in an 8-microphone arena, asterisks denote *p <* 0.05, permutation test); the change in male-female distance between control and blind flies (an average of +4.80 mm) was substantially larger than the largest change among LC types (for LC10a, an average of +1.03 mm; for LC6, -2.15 mm). Differences between our control-spGAL flies and control-CSTul flies are most likely due to the criteria for keeping a session (CSTul sessions were stopped and discarded if the male failed to begin courtship in the first 5 minutes; we did not have such restrictions for our control or LC-silenced sessions). Thus, only the relative changes between control-spGAL and LC-silenced sessions and the relative changes between control-CSTul and blind-CSTul should be compared. **d.** Proportion of sine song given song production. Same data as in Figure 1e (bottom panel) except the LC types are ordered based on increasing proportions. Same format as in **a**. **e.** Mean changes in movement, including forward velocity (top panel), lateral velocity (middle panel), and angular velocity (bottom panel), averaged over the entire session. The absolute value was taken for lateral and angular velocity (i.e., speed), as we were interested in changes away from the male’s heading direction (e.g., a large turn to the right or left both indicated a large deviance). Same format as in **a**. **f.** Changes in the male’s song production, including the probability of sine, Pfast, and Pslow song. Same format as in **a**. Although we observed some significant changes in behavior (asterisks), overall we did not observe any LC types that, after silencing, resulted in changes to behavior on par with the changes observed between control and blind flies—opposite of what we were expecting if only one or two LC types were the dominant contributors to courtship. This suggests that multiple LC types need to be silenced together to obtain large deficits in behavior, consistent with our modeling results (Fig. 4). Previous studies have identified LC types LC10a and LC9 as contributing to courtship [10, 21, 22], and our results are consistent: LC10a and LC9 show an increase in male-to-female distance, as previously reported (**a**, ‘LC10a’ and ‘LC9’). A new implication for LC10a and LC9 is for song production: Both LC types tend towards a reduction in song production for all three song types (**d**, ‘LC10a’ and ‘LC9’). The metrics we use here (e.g., taking the mean forward velocity across an entire session) are coarse summary statistics and do not represent all possible ways in which behavior may change due to silencing. In addition, variability across sessions per LC type was large, making it difficult to identify true changes. This motivated us to use the 1-to-1 network to model the silenced and control behavior, as the 1-to-1 network can be used to directly identify the largest changes to the sensorimotor transformation due to LC silencing. In particular, we can use a metric—the coefficient of determination *R*^2^—that considers more possible changes than simply a change in mean offset. We use *R*^2^ when comparing changes to behavior for the 1-to-1 network (Figure 4), but we cannot use *R*^2^ for the data here, as the visual inputs were not the same across silenced behavioral datasets. **g.** Our behavioral experiments comprised two sets of data collection that were 4 years apart, and we wondered if large deviations occurred for control-spGAL sessions between the two sets (both sets had the same genetic lines). We separated the control sessions into two groups (‘control 2017’ and ‘control 2021’, named for the year of collection) and found no significant difference between them across the movement and song statistics (n.s. denotes *p >* 0.05, permutation test) except for Pfast song (asterisk, *p <* 0.05, permutation test). Thus, we felt confident in merging the two sets of data collection for further analysis.

**Extended Data Figure 2:**
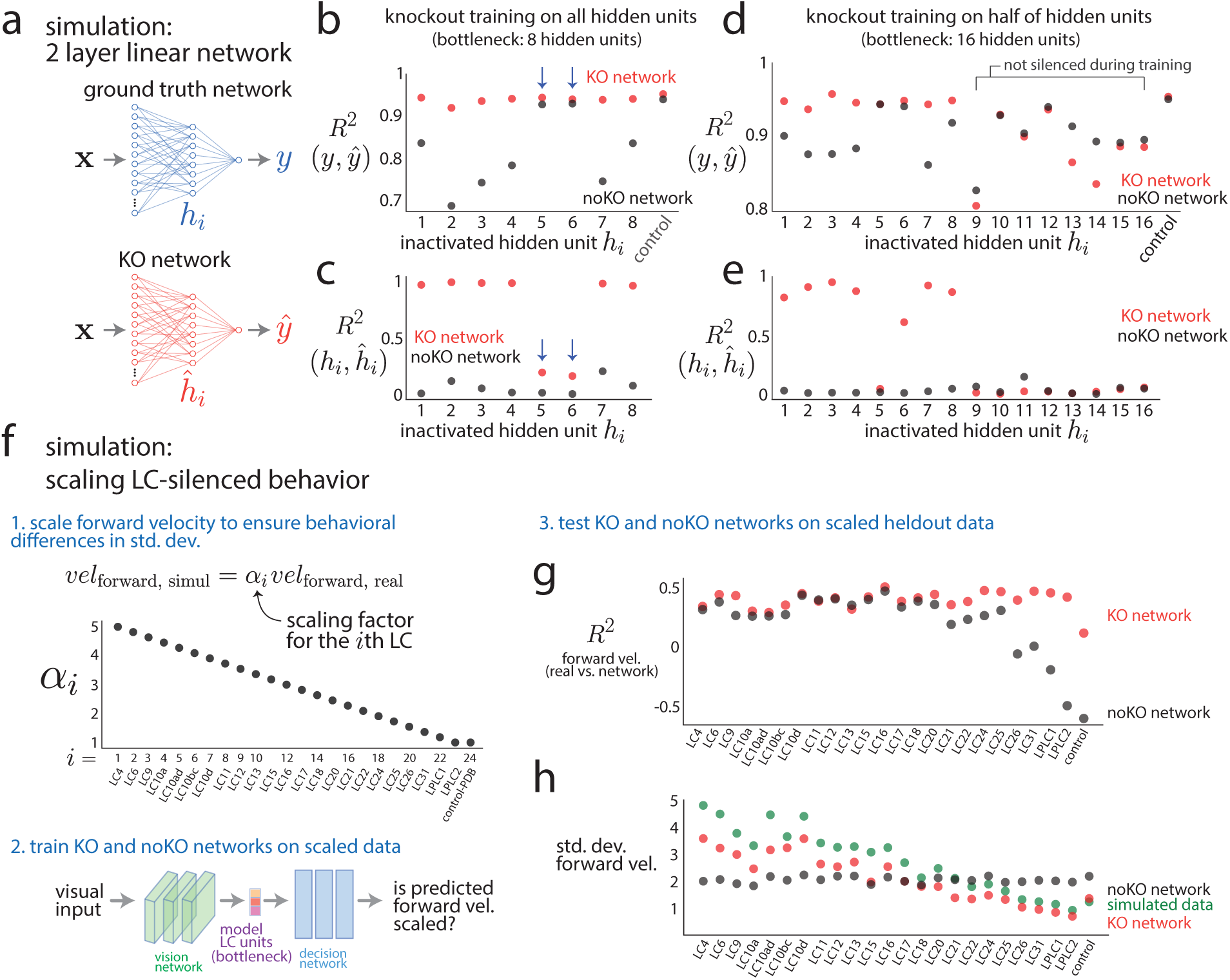
Testing the efficacy of knockout training with simulations. We tested the ability of knockout training to correctly identify the one-to-one mapping of silenced neuron types with two simulations. We compare a network trained with knockout (‘KO network’) to a network trained *without* knockout (‘noKO network’) for which no model units are inactivated (i.e., the noKO network has no knowledge that any silencing occurred). **a.** A simple simulation with 2 layer linear networks. The ground truth network (top) is a randomly-initialized, untrained 2 layer linear network with 48 input variables (**x** *∈ R*^48^ where *x_j_ ∼ N* (0, 1)), 8 hidden units (*h_i_* for *i* = 1*, . . .,* 8), and 1 output unit (*y*). We use the same network architecture for the KO network (bottom). We seek a one-to-one mapping between the ground truth hidden units *h_i_* and the model’s estimated hidden units *h*^^^*_i_*. We generated training data by silencing each hidden unit of the ground truth network (i.e., setting *h_i_* = 0) and recording the resulting silenced output *y* as well as observing control data (for which no silencing occurred). For each case, we drew 1,000 input samples, which yielded 9, 000 training samples in total. We then trained the model either using knockout training (‘KO network’) or without it (‘noKO network’). We generated a test set in the same way as the training set but independent of the training set; the test set also had 9, 000 test samples in total. **b.** We tested the KO network’s ability to correctly predict the silenced output *y* of the ground truth network. We collected the KO network’s predicted output *y*^ to 1, 000 test samples for each silenced hidden unit of the ground truth network by knocking out the corresponding hidden unit in the KO network. We then computed the *R*^2^ (coeffi-cient of determination) between *y* and *y*^ for each silenced unit as well as control (red dots). We evaluated the noKO network with the same test set but did not knockout any hidden units during training or evaluation (black dots). We found that the KO network better predicted silenced output than that of the noKO network for most of the hidden units (red dots above black dots) but performance was roughly equal for control data (‘control’ red and black dots overlap). The KO and noKO networks had similar prediction performance for some of the silenced hidden units (*i* = 5 and 6, arrows); these units contributed little to the output of the ground truth network and, when silenced, led to outputs similar to those observed during control sessions. **c.** We then tested the KO network’s ability to correctly predict the hidden unit activity *h_i_*for the *i*th hidden unit of the ground truth network (i.e., its “neural” responses). For the same test set as in **b**, we collected the KO network’s responses of its hidden units *h*^^^*_i_* and computed the *R*^2^ (Pearson’s correlation squared) between *h_i_* and *h*^^^*_i_* (red dots). We performed the same evaluation for the noKO network (black dots) and found that the KO network substantially better predicted the activity of the ground truth’s hidden units versus the noKO network (red dots above black dots). We observed some hidden units with low prediction performance both for the KO and noKO networks (*i* = 5 and 6, arrows). These hidden units matched those with high prediction performance of model output in **b**; as expected, knockout training cannot identify mappings for hidden units that contribute little to the ground truth network’s output. Taking **b** and **c** together, we conclude that knockout training successfully identified the one-to-one mapping. **d.** We wondered to what extent does knockout training recover the one-to-one mapping when not all ground truth hidden units are silenced. This setting is more similar to our modeling of the fruit fly visual system, where we cannot silence all possible LC types. To test this, we gave the ground truth network and the model network each 16 hidden units (instead of 8 units) but only silenced the first 8 hidden units of the ground truth network (*i* = 1, 2*, . . .,* 8). We generated the training and test sets in the same manner as in **a**, ignoring the extra last 8 hidden units (*i* = 9, 10*, . . .,* 16), and trained the network with knockout training. The KO network correctly predicted output *y* for the first 8 silenced units (*h*_1_*, h*_2_*, . . ., h*_8_) but not for output resulting from silencing the last 8 units on which the KO network was not trained (*h*_9_*, h*_10_*, . . ., h*_16_, red and black dots overlap). The noKO network had worse prediction than that of the KO network for hidden units that contributed to the output (*h*_1_*, h*_2_*, . . ., h*_8_, black dots below red dots for most hidden units); inactivated hidden units with similar performance between KO and noKO networks (red dots and black dots overlap, *i* = 5 and 6) are due to the same reasons as that in **b** (arrows). **e.** Same as **c** except for 16 hidden units. As expected, the KO network recovers the activity for most of the first 8 hidden units (*h*_1_*, h*_2_*, . . ., h*_8_, red dots above black dots) but fails to recover the activity of the last 8 hidden units (*h*_9_*, h*_10_*, . . ., h*_16_, red and black dots close to *R*^2^ = 0). We note that the KO and noKO networks have similar poor performance *R*^2^(*h_i_, h*^^^*_i_*) for hidden units 5 and 6 for the same reasons as the hidden units 5 and 6 in **b**. Taking **d** and **e** together, we conclude that knockout training still works to identify a one-to-one mapping (predicting both output/behavior and response) for hidden units that have been silenced—even if the remaining units in the bottleneck are never silenced. This motivates us to train the 1-to-1 network on behavioral data from silencing 23 LC types individually even though we do not have access to behavioral data from silencing the other remaining LC types in the bottleneck (*∼*45 LC types total in the bottleneck). **f.** Given that knockout training works in a simple simulation setting (**a**-**e**), we moved to testing knockout training for the 1-to-1 network used to model the fruit fly visual system (Fig. 1**a**). Although we could simulate data coming from a trained 1-to-1 network as ground truth, we were more interested in the case where there was a mismatch between the model and the real system—almost certainly the case using the 1-to-1 network to predict LC neuron types. Still, we sought some way to assess a ground truth change in behavior for the LC-silenced data and devised the following approach. For the *i*th LC type, we scale its forward velocity by *α_i_*, where *α_i_* decreases from 5 to 1 as index *i* increases incrementally. We then train a KO network on this scaled data in the same manner as that of the 1-to-1 network; we also train a noKO network. We only train the networks to predict forward velocity (no other behavioral outputs). **g.** We computed prediction performance *R*^2^ (coefficient of determination) between predicted and actual forward velocity on heldout frames for each LC-silenced behavior. The KO network had better prediction than that of the noKO network both for the most scaled and least scaled LC types (red dots above black dots for leftmost and rightmost dots). **h.** This change in performance between KO and noKO networks for the most and least scaled LC types in **g** can be explained by how the KO and noKO networks each predicts the standard deviation of forward velocity. As expected from our scaling of the real data (**f**), the standard deviation of the simulated data linearly falls as we consider the later LC types (green dots, compare with black dots in **f**). We find that the standard deviations predicted by the KO network also linearly decrease (red dots) while those predicted by the noKO network remain relatively flat (black dots). Because the noKO network has no information about which LC type was silenced, the noKO network must predict roughly the same standard deviation for all LC types, choosing an intermediate standard deviation (around 2 s.d.). This also helps to explain why the KO and noKO networks differed in prediction more for the rightmost LC types (**g**, ‘LC26’ to ‘control’) because the noKO network overestimated the standard deviation for these LC types (black dots above green dots for ‘LC26’ to ‘control’) leading to larger errors (and a negative coefficient of determination) versus underestimating the standard deviation which does not lead to as large of drops in *R*^2^. These simulations show that knockout training can reliably identify one-to-one mappings between model units and internal units given behavior resulting from silencing those internal units.

**Extended Data Figure 3:**
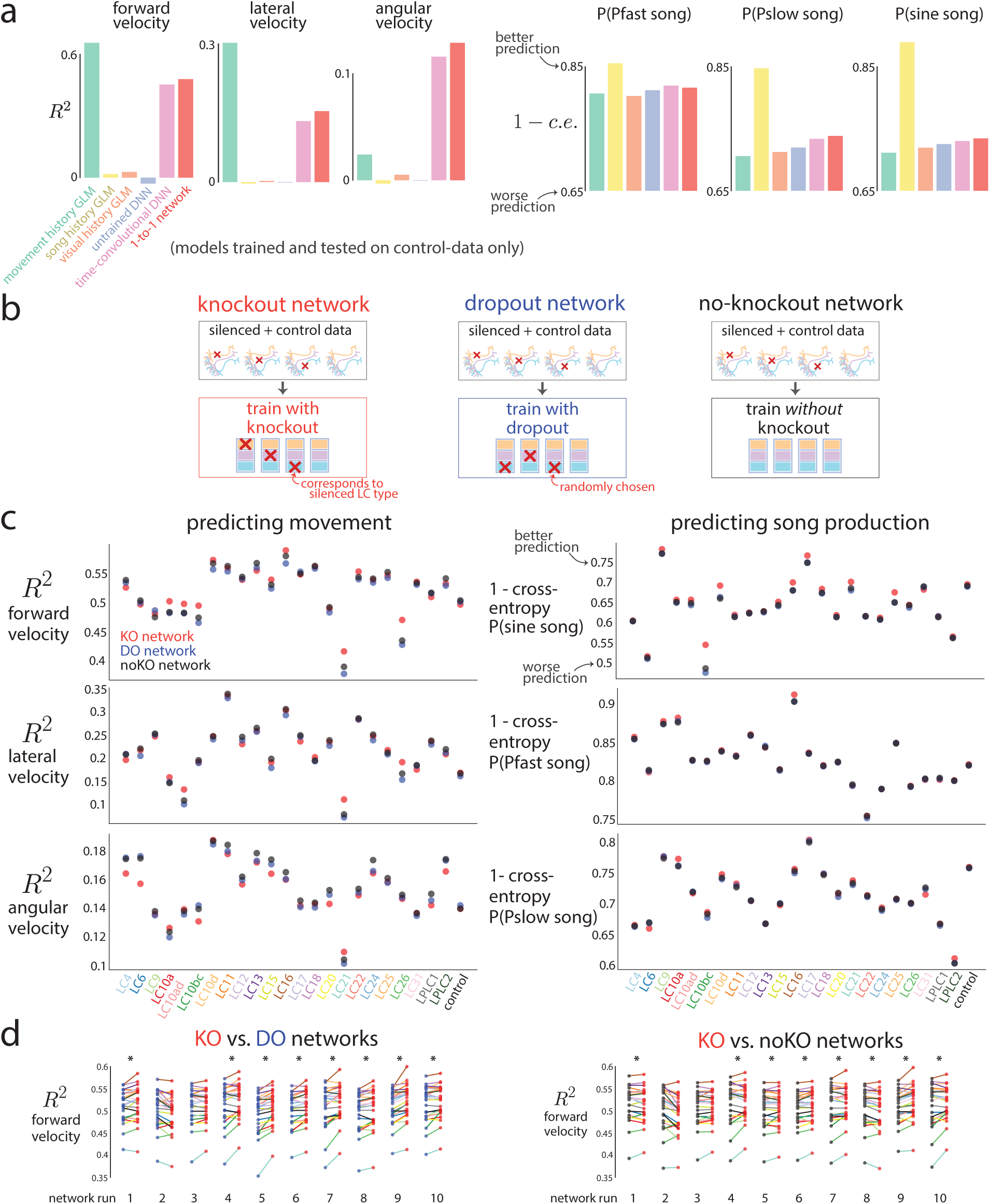
Predicting behavior frame-by-frame. To determine the architecture of the 1-to-1 network, as well as to compare deep neural network models versus baseline models, we trained and tested different models on behavior from control sessions only. Then, as a test of the 1-to-1 network’s ability to uncover a one-to-one mapping between model LC units and real LC neurons, we tested the extent to which the 1-to-1 network accurately predicts behavior on heldout courtship frames for each LC type. An important comparison is to measure the 1-to-1 network’s prediction performance relative to networks with the same architecture and training data but with different training procedures. **a.** We considered different network architectures for the 1-to-1 network and compared their prediction performance to baseline models. We trained each model on control sessions only and tested on heldout test frames of control sessions. For baseline models, we considered a generalized linear model (GLM) that took as input the last 300 ms of movement history, including forward, lateral, and angular velocity (‘movement history GLM‘); past song history, including Pfast, Pslow, and sine song (‘song history GLM’); as well as past visual history, including female size, position, and rotation (‘visual history GLM’). The movement-history-GLM had good prediction of forward and lateral velocity (two leftmost plots), as expected, but failed at predicting angular velocity and song production. Its good prediction stems from the fact that an animal’s forward velocity at time step *t* is likely similar to its forward velocity at time step *t −* 1 based on the physics of the movements. Likewise, the song history GLM best predicted song production (three rightmost plots), as songs often occur in bouts, but failed at predicting moment-to-moment movement (three leftmost plots). Also expected was the poor prediction performance of the visual-history-GLM, whose inputs of the fictive female’s parameters likely must pass through a strong nonlinear transformation to accurately recover behavior (all orange bars are low). Next, we considered the DNN architecture of the 1-to-1 network (Fig. 1**a**). We trained the 1-to-1 network on control data only (i.e., no knockout training was performed) for this analysis. The 1-to-1 network’s prediction performance was better than any GLM model for angular velocity and showed good performance for song production (red bars), confirming that a nonlinear transformation was needed to map visual input to behavioral output. The 1-to-1 network did not outperform the movement-history-GLM on forward and lateral velocity; providing past movement history to the 1-to-1 network is an intriguing direction not investigated in this work. We confirmed that an untrained network with the same architecture as the 1-to-1 network (‘untrained DNN’, only last readout layer was trained) had little prediction ability. Finally, we trained a more complicated version of the 1-to-1 network which had 3-d convolutions in both space (2-d) and time (1-d) in the vision network (‘time-convolutional DNN’ with 3 *×* 3 *×* 3 convolutional kernels). This greatly increased the number of parameters but ultimately did not improve prediction performance versus the 1-to-1 network (pink versus red bars). We suspect that with more data, the time-convolutional DNN may outperform the current architecture of the 1-to-1 network, as motion processing occurs before the LC bottleneck [80]. **b.** Three different training procedures. Knockout training (left, red) sets to 0 the model LC unit that corresponds to the LC neuron type silenced for that training frame (no model LC units’ values are set to 0 for frames from control sessions). We refer to the resulting trained network as the knockout (KO) network or, interchangeably, as the 1- to-1 network. Dropout training [40] (middle, blue) sets to 0 a randomly-chosen model LC unit for each training frame, independent of the frame’s silenced LC type (no model LC units’ values are set to 0 for frames from control sessions). In this case, the number of ‘dropped out’ units equals that of the ‘knocked out’ units. We refer to the resulting trained network as the dropout (DO) network. Finally, we train a network *without* knocking out any of the model LC units and refer to it as the noKO network (right, black). An important difference between the KO network and the DO and noKO networks is that the latter have no knowledge that any LC silencing has occurred; in other words, the DO and noKO networks assume all male flies, regardless of an LC being silenced or not, have the same computations. Thus, the DO and noKO networks cannot reliable detect changes in behavior for different silenced LC types unless the statistics of the visual input itself differs across silenced LC types (e.g., if silenced flies do not chase the female, the female will be visually smaller for most frames, leading DO and noKO networks to correctly predict a decrease in song production, as song is produced in close proximity to the female). **c.** We tested the KO, DO, and noKO network’s performance of predicting the male fly’s movement (left) and song production (right) for the next frame given the 10 past frames of visual input across many LC-silenced and control flies. All test frames were held out from any training or validation sets and sampled randomly across sessions (27,000 test frames per each LC type and control, see Methods). We computed the coefficient of determination *R*^2^ for behavioral outputs of movement (forward, lateral, and angular velocity) and 1 - binary cross-entropy (where a value close to 1 indicates good prediction) for behavioral outputs of song production (probabilities of Pfast, Pslow, and sine song). We found that overall, the KO network better predicts forward velocity than the DO and noKO networks (top left, red dots above black and blue dots) as well as probability of sine song (top right). Changes in prediction performance between KO and DO/noKO networks across LC types were relatively small, suggesting changes in behavior were subtle, consistent with overall mean changes in behavior (Ext. Data Fig. 1). In addition, *R*^2^ may change little for large second-order changes in behavior, such as variance (Ext. Data Fig. 2**g**, leftmost dots). We confirmed in Figure 1g-h and Extended Data Figure 4 that the KO network accurately predicted mean changes in behavior better than DO and noKO networks. We note that *R*^2^ values for movement (left column, *R*^2^ *≈* 0.5 for forward velocity, *R*^2^ *≈* 0.15 for lateral and angular velocity) were not close to 1 because we predict rapid changes to movement variables frame-to-frame (with a frame rate of 30 Hz). Because the 1-to-1 network is deterministic (i.e., returning the same output for the same visual input), it fails to account for the fact that a male fly’s moment-to-moment decision is stochastic—in other words, the male responds differently to repeats of the same stimulus sequence. To take this stochasticity into account, one would need to present identical repeats of the same visual stimulus sequence and record the resulting behavior. This is not possible for our natural courtship experiments, where a male fly’s visual experience is determined by his behavior. However, this may be possible in experiments using virtual reality, where the experimenter has greater control over a male fly’s visual input. **d.** Results in **c** were for a KO network with one random initialization. To see if this effect holds for different initializations, we trained 10 runs of the KO network, each with a different random initialization and random ordering of training samples. We found that for 8 of the 10 runs, KO networks outperformed DO networks (left) and noKO networks (right) in predicting forward velocity. For each run (i.e., ‘network run 1’), the same randomly initialized network and randomized order of training samples was used as a starting point for the KO, DO, and noKO network. Each connected pair of dots denotes one LC type with the color of the line connecting two dots denoting the LC type identity (same colors as in **c**). An asterisk denotes a significant difference in means (*p <* 0.05, paired, one-sided permutation test). Network run 1 was chosen as the 1-to-1 network in **c** as well as Figures 1-4 due to its high prediction for both behavior and neural responses.

**Extended Data Figure 4:**
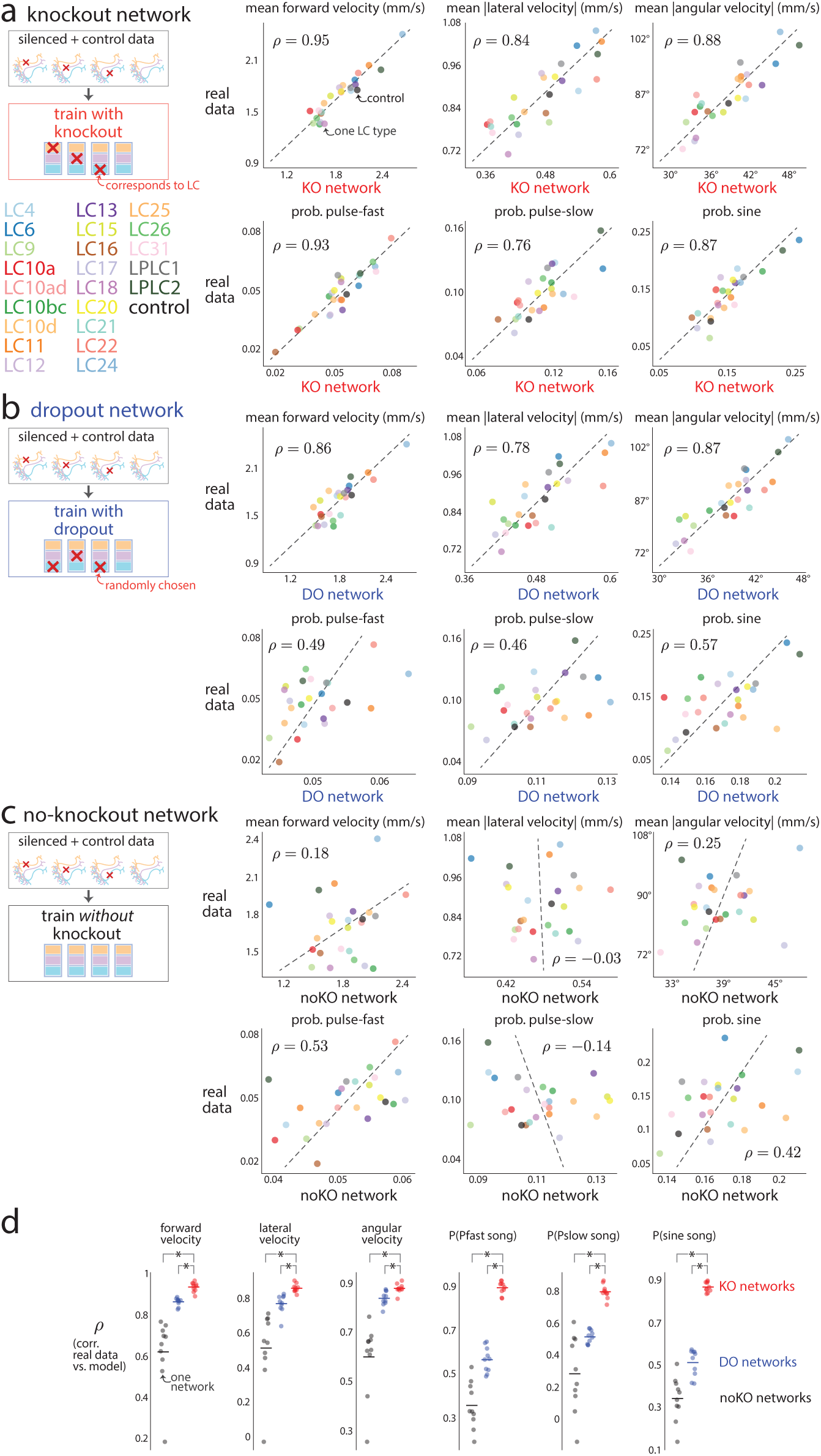
Assessing model predictions of mean changes in behavior. Given the mean changes in behavior due to silencing (where the mean is taken over the entire session, Ext. Data Fig. 1), we wondered to what extent the knockout (KO) network predicted these overall changes versus training a dropout (DO) network and a noKO network for which no inactivation of any model LC unit was performed during training. **a.** For each LC neuron type, we computed the average behavior across all heldout courtship frames in the test set (‘real data’). We then computed the mean behavior as predicted by the KO network across the same frames (‘KO network’). Each dot denotes one LC type (or control session); colors are indicated at left. Dashed lines are the best linear fit between the two variables; the correlation *ρ* is taken across all LC types excluding the control sessions (black dot). The KO network has large *ρ*’s across behavior outputs, indicating good prediction of overall changes. **b.** Same as in **a** except for the DO network (where we do not inactivate any model LC units during evaluation). Correlations were smaller for the DO network than for the KO network (compare *ρ*’s between **a** and **b**). However, for the movement variables, correlations for the DO network were only slightly smaller than those of the KO network. Because the DO network had no access to which LC type was silenced, this suggests that the distribution of visual inputs differed across LC types. For example, imagine if the DO network accurately predicted the behavior of control male flies, including that the male does not sing when the female is far away. Then, if silencing LC10a resulted in the male not being interested in courting the female, the female would be far away in most frames, and the DO network would predict a decrease in song production, which we observe (Ext. Data Fig. 1). Thus, DO training is a good control to ask whether the sensorimotor transformation has changed or if the male has altered his desire to pursue courtship. **c.** Same as in **a** and **b** except for the noKO network. Correlations were substantially smaller than for the KO and DO networks (compare *ρ*’s between **a** and **c**), indicating that the noKO network could not recover behavior from LC-silenced flies. We trained 10 networks each for KO, DO, and noKO training. Each of the 10 networks had different random initializations and different random orderings of training samples. For a fair comparison, the same initialized network and ordering was shared across KO, DO, and noKO training for each of the 10 runs. We then computed the *ρ*’s of overall mean behaviors for each network and real data. For each of the six behavioral outputs, we found that the KO network better predicted the changes in behavior across LC types better than the predicted changes of the DO and noKO networks (red dots above blue and black dots). Each dot denotes one network, and each asterisk denotes that the mean of the KO network is significantly greater than the mean of either the DO or noKO network (*p <* 0.05, paired, one-tailed permutation test). Network run 1 was chosen as the exemplar network in **a**-**c** (as well as in Figs. 1-4).

**Extended Data Figure 5:**
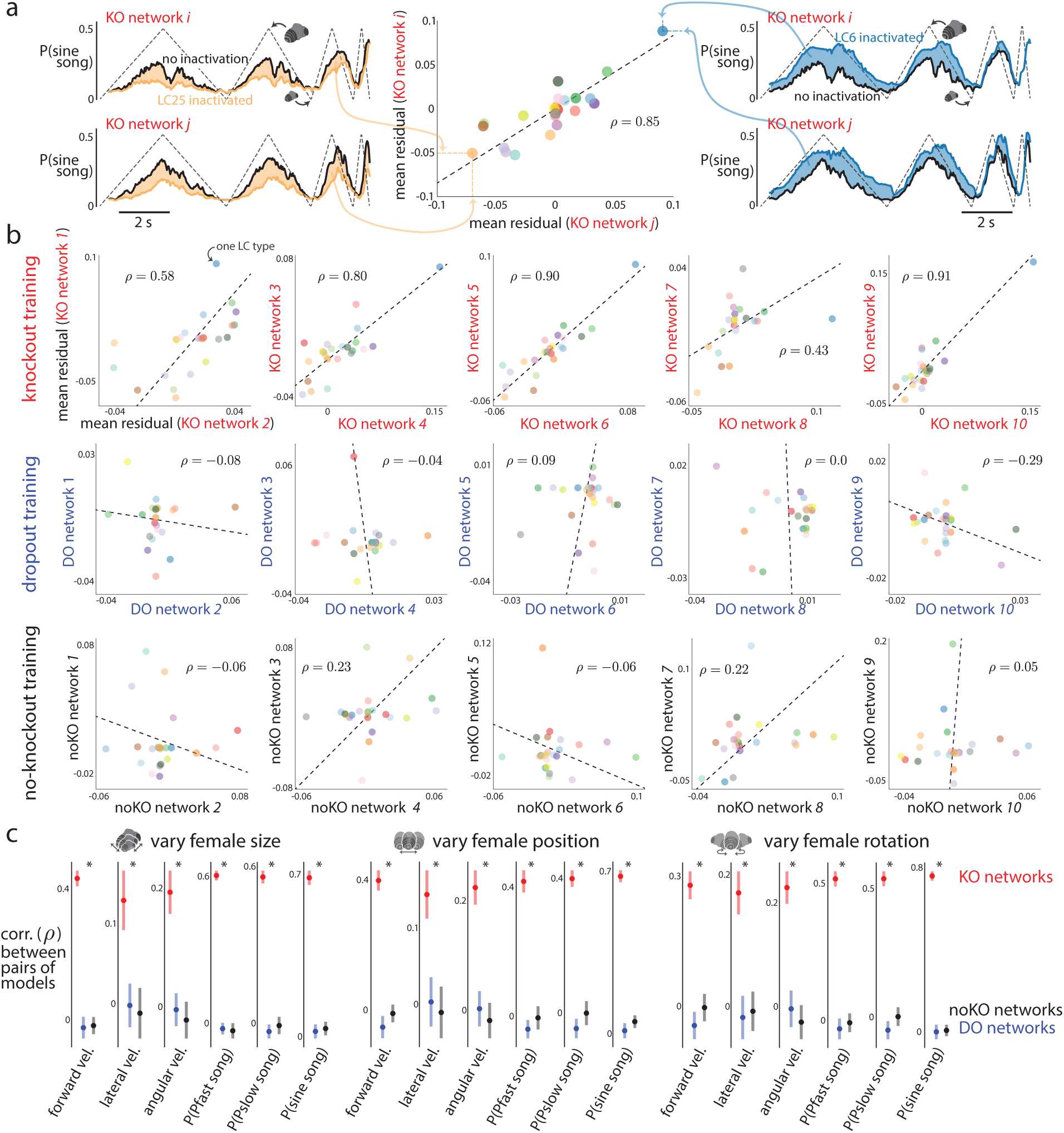
Consistency in behavioral predictions across networks with different random initializations. Because deep neural networks are trained with stochastic gradient descent, the final, optimized network is dependent both on its initialization and the ordering of training samples. We wondered to what extent the solution identified by knockout (KO) training changes for different random initializations and different orderings of training data. If KO training is consistent for the 1-to-1 network’s architecture, then we would expect to see that different training runs of a KO network should converge to similar predictions in behavior. See Extended Data Figure 6 for the 1-to-1 network’s consistency in neural predictions. **a.** To test this, we trained 10 KO networks, each with a different random initialization and different ordering of training samples. We then passed as input a dynamic stimulus sequence in which the fictive female varied her size over time (dashed trace in top left plot; female position and rotation remained fixed). Inactivating LC25 (orange line, top left plot) resulted in an overall decrease in the probability of song relative to that of no inactivation (black line, top left plot); we can compute the overall change in behavior by taking the mean residual between the two (orange shade, top left plot). If two KO networks were consistent, we would expect that this KO network *i* should match its mean residual if we were to perform the same procedure for another KO network *j*; indeed, this is what we saw (compare top and bottom panels on the left). Inactivating LC6 resulted in an increase in probability of song for both KO networks (rightmost plots). We quantified the consistency between the two KO networks by computing the correlation across LC types of the mean residuals (middle scatter plot); a correlation *ρ* close to 1 indicates that both KO networks consistently have the same predictions of behavior for different silenced LC types. **b.** Scatter plots for 5 pairings of the 10 KO networks (top row) of the mean residuals of probability of sine song for the stimulus in which the fictive female varies in size (same as in **a**). Each dot denotes one LC type, and colors correspond to LC names in Extended Data Figure 4. Dashed lines denote best linear fit. We also assessed the consistency of DO networks (middle row) and noKO networks (bottom row), which tended to have lower *ρ*’s. **c.** Correlations of mean residuals for the three dynamic stimulus sequences and all six behavioral outputs. Each dot is the mean across all 45 pairs of networks; error bars denote 1 s.e.m. The KO networks had significantly larger mean correlations (asterisk denotes *p <* 0.05, paired permutation test) than those for the DO and noKO networks (red dots above blue and black dots) for all stimulus sequences and behavioral outputs. We conclude that the KO networks are consistent in behavioral predictions. An important use of the ensemble of 10 KO networks is for estimating model uncertainty for a particular stimulus sequence. A single KO network can only give one prediction for a stimulus sequence (Fig. 4); one may erroneously conclude that the model is equally certain about all stimulus sequences. Instead, the 1-to-1 network may be more uncertain for different stimulus sequences, especially those that are rarely observed during natural courtship. Thus, before experimentally testing the 1-to-1 network’s predictions, one may first check if the 1-to-1 network is confident in its prediction by assessing the extent to which different network runs agree on the same prediction. If there is large agreement (as seen here), the 1-to-1 network is confident in its predictions. On the other hand, a mismatch in its predictions and experimental data is more interesting than for a stimulus sequence for which the 1-to-1 network is uncertain (and thus likely will not agree with experimental data).

**Extended Data Figure 6:**
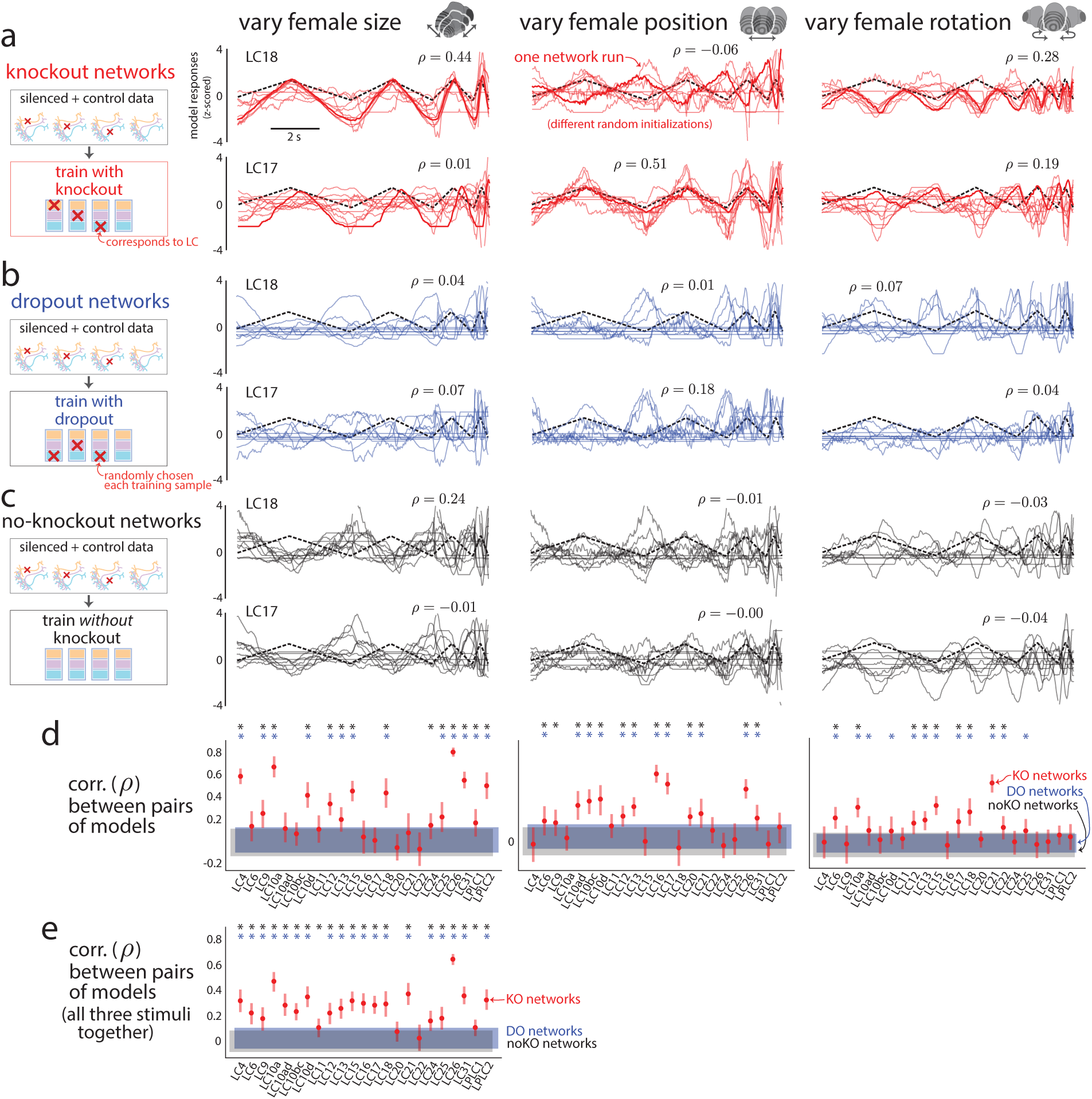
Consistency in LC response predictions across networks with different random initializations. We wondered to what extent knockout training converged to different solutions in predicting LC responses given different random initializations and different orderings of training data. See Ext. Data Fig. 5 for consistency in behavioral predictions. **a**. We performed knockout training on 10 different runs—each run had a different random initialization and different random ordering of training data. We then passed into the KO networks as input three dynamic stimulus sequences in which the fictive female varied her size (left column), position (middle column), and rotation (right column) (same sequences as in Fig. 3**f** and Fig. 4**d** and Fig**e**). For LC18 (top row), model responses were consistent for female size and rotation but not position. Each trace is from one KO network run; the bold trace is for network run 1 (chosen as the 1-to-1 network in Figs. 1-4). Traces across all three stimulus sequences were z-scored and then flipped in sign to ensure the largest possible mean correlation *ρ* over time. For LC17 (bottom row), model responses were consistent for female position but not size or rotation. This indicates that the KO networks were consistent for different stimulus sequences depending on the LC type. This is in line with the idea that knockout training can only identify a one-to-one mapping for stimulus sequences for which silencing an LC type leads to a change in behavior (Ext. Data Fig. 2); KO networks disagree on stimulus sequences for which little to no change in behavior occurs, as some change is needed in order to identify an LC type’s role in driving behavior. **b**-**c**. We assessed the consistency of the dropout (DO) networks (**b**) and noKO networks for which no inactivation was performed (**c**). Both DO and noKO networks had poor consistency for LC18 and LC17 across all stimulus sequences (the largest *ρ* = 0.24). **d.** We computed the mean correlation (dots) across all 45 pairs of networks and found that the KO networks had significantly larger mean correlations than DO networks (blue asterisks, *p <* 0.05, paired, one-sided permutation test) and noKO networks (black asterisks, *p <* 0.05, paired, one-sided permutation test) for the three different stimulus sequences. **e.** We concatenated the responses for each network across all three stimulus sequences and re-computed the mean correlation (dots). Almost all of the LC types show a significant increase in mean correlation. Error bars in **d** and **e** denote 1 s.e.m. Taken together, these results indicate that knockout training identified consistent KO networks that reliably predict neural responses. That KO networks were more consistent than DO and noKO networks suggests that knockout training captured meaningful changes in behavior. Because KO networks may disagree more for different stimulus sequences (a notion of uncertainty), future experiments should take this uncertainty into account when testing the 1-to-1 network’s predictions. In fact, presenting stimulus sequences for which the KO networks disagree the most may be the most informative, as we can use the responses to these sequences to rule out some of the KO networks.

**Extended Data Figure 7:**
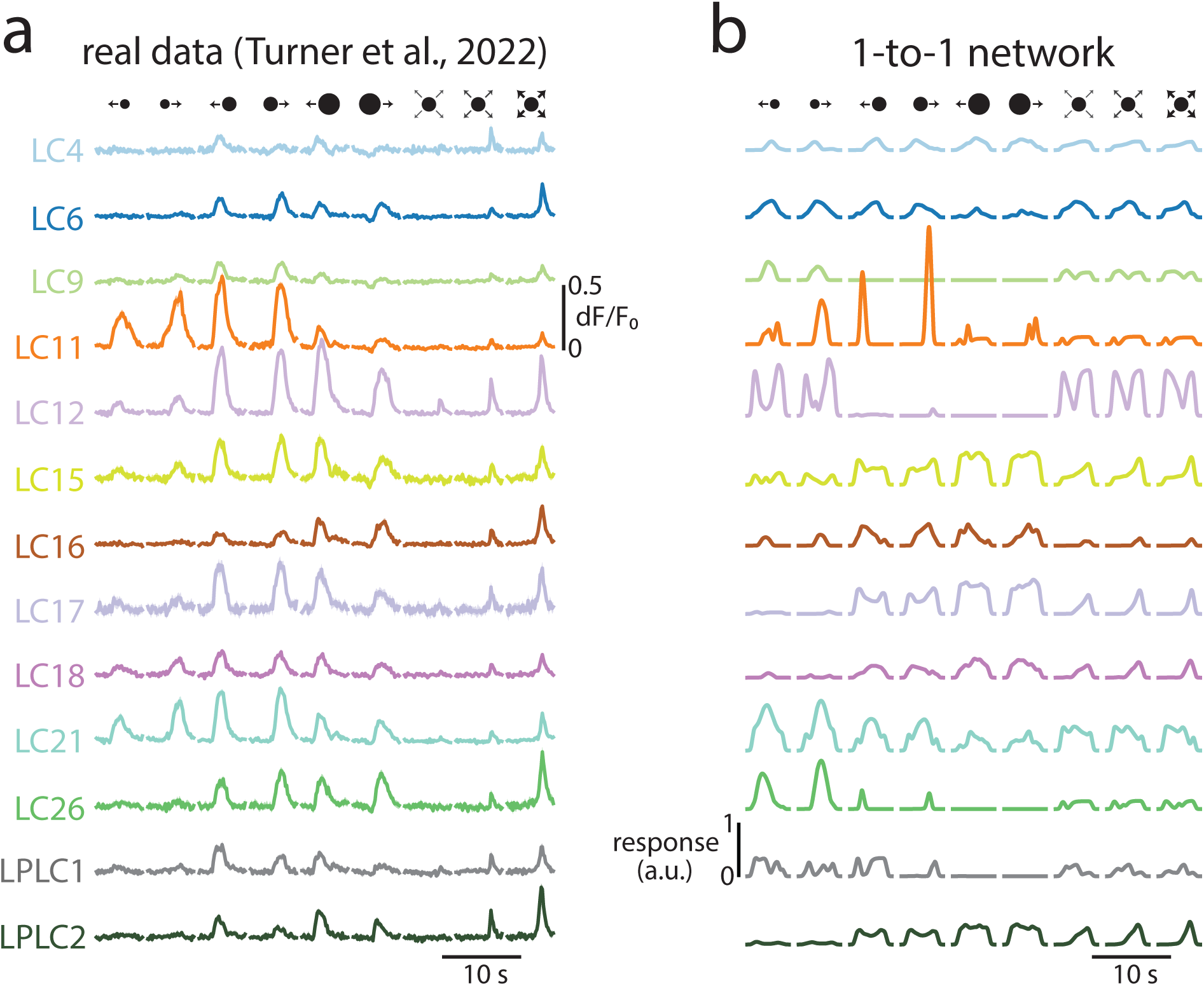
Predicting real LC responses to artificial stimuli. We further tested the 1-to-1 network’s neural predictions on a large number of LC neuron types whose responses were recorded in another study [32]. **a.** Real LC responses from [32]. We considered responses to artificial stimuli of laterally moving spots with different diameters and different movement directions as well as looming spots with different loom accelerations (top row). Traces denote responses averaged over repeats and flies, shaded regions denote 1 s.e.m. (some regions are small enough to be hidden by the mean traces). Data same as in Fig. 3**a** of [32]. **b.** Model LC responses from the 1-to-1 network. We passed through the same stimulus dynamics but changed the spot to a fictive female facing forward (to better match these artificial stimuli to the fictive female stimuli on which the 1-to-1 network was trained). For visual comparison, we matched the mean and standard deviation (taken across all stimuli) of each LC type’s model responses to those of the real LC responses; we also flipped the sign of a model LC unit’s responses to ensure a positive correlation with the real LC type (flipping was only performed for LC6 and LC21). To account for adaptation effects, model LC unit’s responses decayed to their initial baseline after no change in the original responses occurred (i.e., for static input; see Methods). Overall, it appeared that almost all the real LC neurons and model LC units respond to these artificial stimuli, providing more support that the LC types form a distributed code. Some of the best qualitative matches were LC11— where the 1-to-1 network correctly identified the object size selectivity of LC11 neurons [26]—LC15, LC17, and LC21. A failure of the model was predicting LC12 responses; this was true of our LC recordings as well (Ext. Data Fig. 8 and Ext. Data Fig. 10). This failure may be due to an unlucky random initialization, although networks trained with knockout over 10 training runs better agreed on LC12’s responses than networks trained with dropout or without knockout (Ext. Data Fig. 6). Another explanation is that LC12 only weakly contributes to behavior for these simplified stimuli. If this were the case, then KO training would not be able to identify LC12’s contributions to behavior nor its neural activity. One piece of evidence that this might be the explanation is that solely inactivating LC12 for simple, dynamic stimulus sequences did not lead to any change in the model’s behavioral output (Fig. 4**f** and Data Fig. 16). For natural stimulus sequences, LC12 does appear to play a role (Fig. 4**c**), motivating the use of more naturalistic stimuli when recording from LC types (Fig. 2).

**Extended Data Figure 8:**
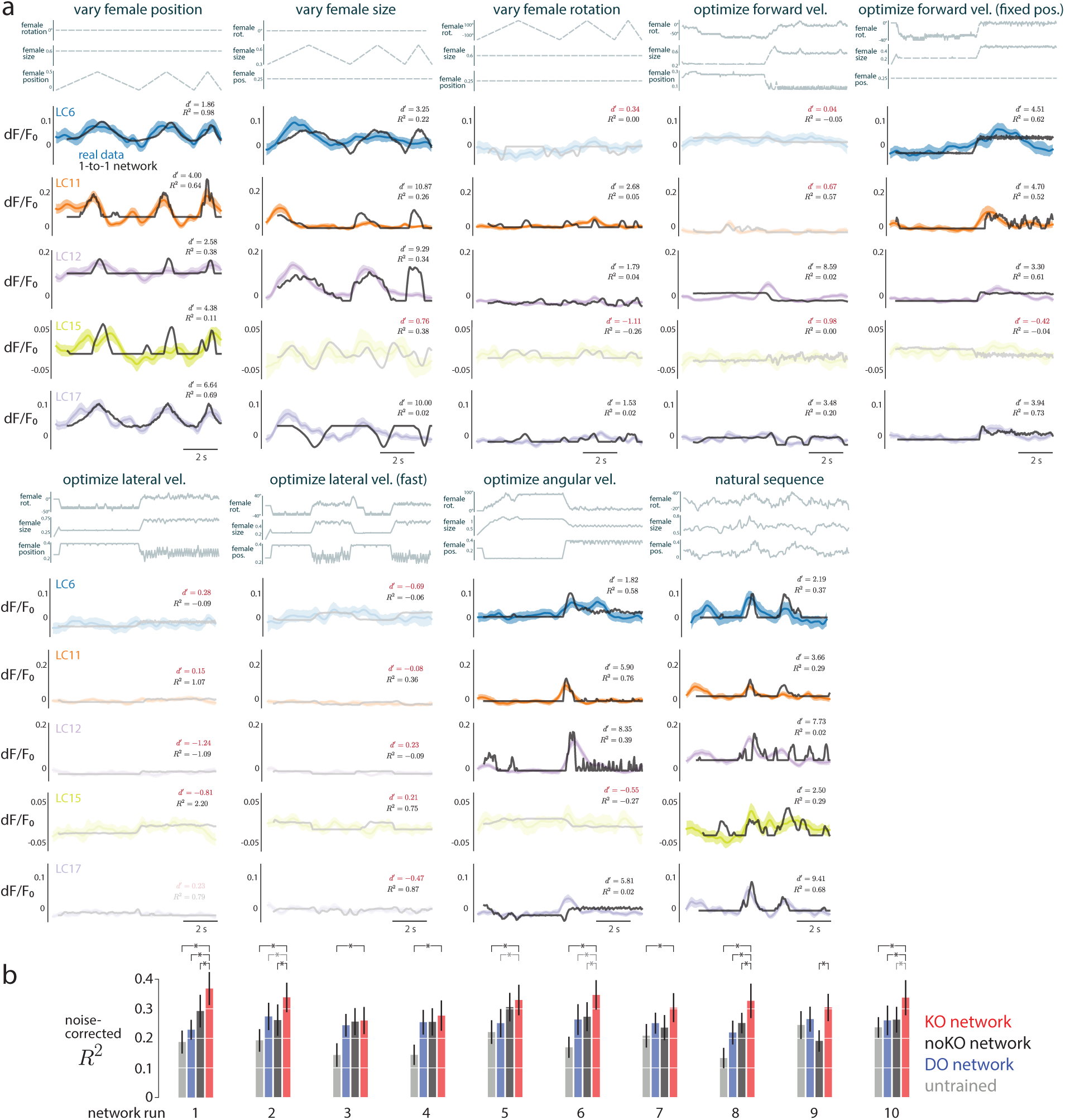
Real LC responses and predicted responses to stimulus sequences of a moving fictive female. **a.** We considered 9 different stimulus sequences in which a female varied her rotation, size, and position (three top traces for each stimulus sequence, see Methods for stimulus descriptions). We found that the 1-to-1 network’s predictions (black traces) largely predicted the responses of the real LC neurons (color traces), despite the facts that the 1-to-1 network was never given access to neural data and that we directly read out from a single model LC unit. The average of all reported noise-corrected *R*^2^s here is the same as that reported in Figure 2e. We only considered stimulus sequences for which the real LC responses reliably varied across time for the stimulus sequence. To measure this, we computed the *d^i^*between splits of repeats (i.e., a signal-to-noise ratio across repeats) and considered any stimulus sequence with a *d^i^<* 1 as unreliable, removing it from our analyses (translucent traces; see Methods). For some LC types, we detected a reliable response to only one or a few stimuli (e.g., LC15 only responded to ‘vary female position’ and ‘natural sequence’). We noticed that none of the LC neurons responded to stimulus sequences for which the fictive female’s parameters were chosen to optimize the 1-to-1 network’s output of lateral velocity (‘optimize lateral vel.’ and ‘optmize lateral vel. (fast)’, see Methods). This may be due to the fast changes in female position which were not present in other stimulus sequences. For each stimulus and LC type, we computed a noise-corrected *R*^2^ between the real and model predicted responses. This noise-corrected *R*^2^ overlooks any differences in mean, standard deviation, and sign of the response, which are unidentifiable by the KO network. For visual clarity, we centered, scaled, and flipped the sign of the 1-to-1 network predictions (black traces) to match the mean, standard deviation, and sign of the LC responses (color traces) for each stimulus. We accounted for the smoothness of calcium traces by applying a causal smoothing filter to the model LC responses as well as fitting the mean offset of the relu thresholding (see Methods). Interestingly, all LC types responded reliably to the varying of female position (‘vary female position’, color traces) despite the facts that the optic glomeruli have weak retinotopy [11, 41] and that the calcium trace is a sum of the activity of almost all neurons for the same LC type (presumably averaging away any spatial information). This suggests that either our targeted region for calcium imaging (Fig. 2**a**) was biased to read out from a subset of LC neurons with nearby receptive fields or that these LC neurons have some selectivity in female position (perhaps as direction selectivity). The latter may be more likely, as the male needs to better estimate female position than can be done simply by comparing coarse differences between the two optic lobes. Consistent with our findings, a previous study has identified another LC type—LC10a—to respond to an object’s position [10]. That our 1-to-1 network also predicted positional selectivity in the LC types (black traces) supports the notion that some optic glomeruli may track female position despite weak retinotopy. More work is needed to understand how object position is encoded within a single optic glomerulus and how that information is read out [41]. **b.** Results in **a** were for a KO network with one random initialization. To see if this effect holds for different initializations, we trained 10 runs of the KO network, each with a different random initialization and random ordering of training samples. We compared the KO network to dropout (DO) networks as well as noKO networks for which no knockout occurred during training. These are the same networks used to predict moment-to-moment behavior (Ext. Data Fig. 3**d**). Error bars denote 1 s.e.m., and black asterisks denote a significant increase in mean *R*^2^ (*p <* 0.05, paired, one-sided permutation test), gray asterisks denote a trend (*p <* 0.1). We observed that random initialization played more of a role for neural prediction than for behavioral prediction (Ext. Data Fig. 3**d**). This is not so surprising, as the networks were never trained to predict neural responses. Still, the KO networks tended to outperform the other types of networks (red bars larger than other bars). In addition, the untrained networks performed poorly (gray bars), indicating that behavioral training did improve neural predictions.

**Extended Data Figure 9:**
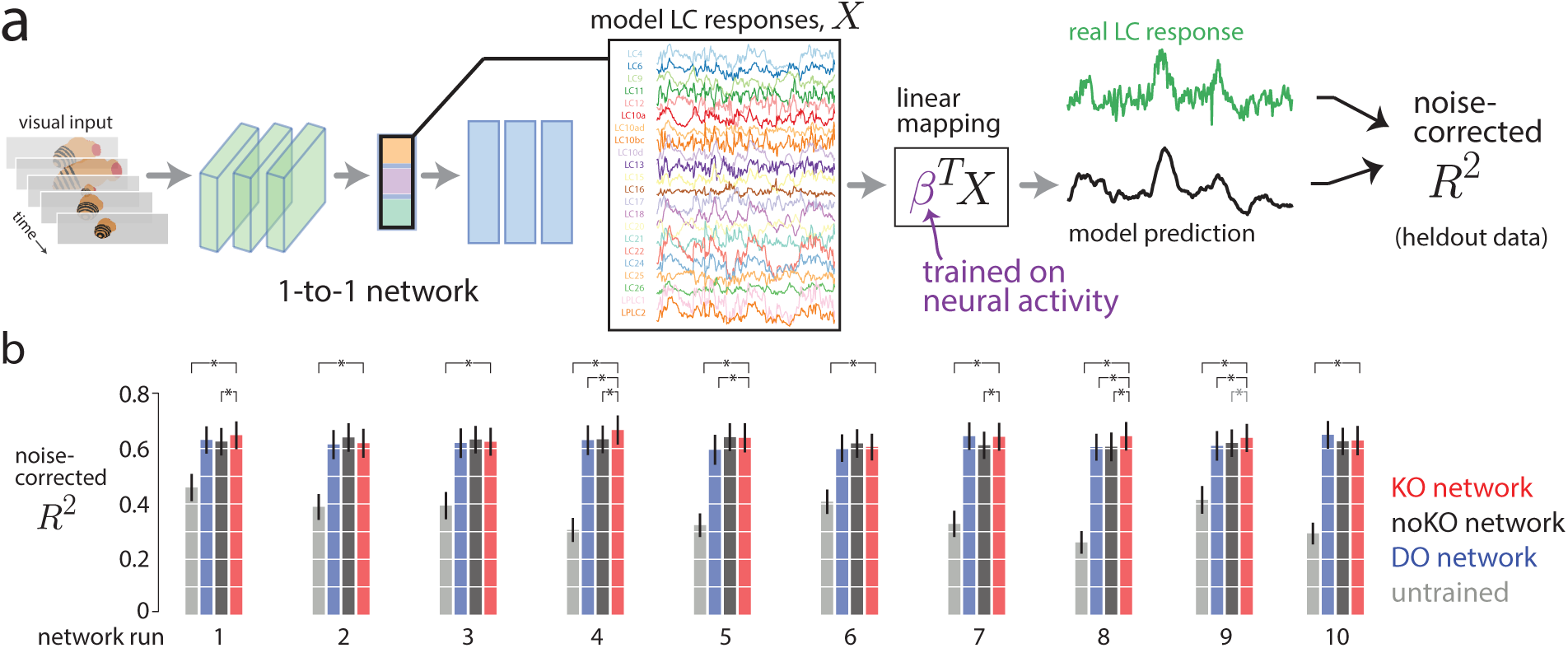
A linear mapping from all model LC units to real LC neurons yields high prediction performance. The 1-to-1 network, with no access to neural data during training, was predictive of real LC responses to stimulus sequences in which a fictive female varied her size, position, and rotation (Fig. 2). Here, we wondered if we were to give our 1-to-1 network access to neural data for training (i.e., train a linear mapping from all model LC units to real LC responses), to what extent would the model’s prediction of real LC responses improve. **a.** Basic setup. We input a stimulus sequence into the 1-to-1 network (fully trained with knockout training) and collect responses from all model LC units, denoted as *X ∈ R^K×T^* for *K* model LC units (here, *K* = 23) and the *T* timepoints of the stimulus sequence. We then define a linear mapping *β ∈ R^K^* to map the *K* model LC responses to the real LC response. We use real LC responses to train *β*. Specifically, for each of the 4 cross-validation folds, we train *β* on 75% of the real LC responses (randomly selected) using ridge regression. We then predict the responses for the remaining heldout timepoints. We concatenate the predictions across the 4 folds and then compute the noise-corrected *R*^2^ in the same way as in Figure 2f-h. Thus, the reported cross-validated noise-corrected *R*^2^’s indicate to what extent the 1-to-1 network, given neural data on which to train, can predict heldout real LC responses. Another view is that in this setting, the 1-to-1 network is a task-driven model trained on behavioral data with an internal representation (the model LC bottleneck) that reflects the activity of real LC neurons up to a linear transformation [1]. **b.** Prediction performance using the linear mapping for different networks and network runs (see Methods). For each network, we trained a new linear mapping between the model LC responses and the real LC responses. Overall, prediction performance greatly increased: The 1-to-1 network (or KO network) with the linear mapping had a noise-corrected *R*^2^ at *∼*65% (network run 1, averaged over all recorded LCs and fictive female stimulus sequences), an additive increase of *∼*30% over that of the 1-to-1 network with the one-to-one mapping comparison (*∼*35%, Fig. 2**e**). We also found that, for the linear mapping, the performance of the 1-to-1 network was similar to those of the other networks trained with dropout (DO) or no knockout (noKO) (leftmost plot, red bar close to black and blue bars). This similarity in performance was not unexpected and indicates that all 3 networks (KO, DO, and noKO) have similar internal representations (up to a linear transformation) at the layer of their LC bottlenecks. However, the 1-to-1 network’s representation is better aligned along its coordinate axes (i.e., where each model LC unit corresponds to one axis) than those of the other networks (Fig. 2**e** and Ext. Data Fig. 8**b**) when comparing those axes to the LC neurons. The untrained network was predictive of LC responses (bar for ‘untrained’ above 0), indicating that this network’s convolutional filters, even with randomized weights, could detect large changes of the visual stimulus (e.g., a fictive female moving back and forth). That a linear combination of random features is often predictive in a regression setting is a well-studied phenomenon in machine learning [81] and has been observed in predicting visual cortical responses [82]. This trend in similarity of performance held across all 10 network runs for the different training procedures: The KO network consistently better predicted real LC responses than the untrained network but less so when compared to the DO and noKO networks (red bars at similar heights to black and blue bars across network runs); for a number of runs, the KO network did perform the best (runs 4, 8, and 9). A black asterisk indicates a KO network with a mean prediction performance significantly above that of another network (*p <* 0.05, paired, one-sided permutation test); a gray asterisk indicates a trend (0.05 *< p <* 0.1). Network run 1 was the chosen 1-to-1 network for Figures 1-4. These results indicate that by simply training a network on courtship behavioral data (i.e., a task-driven approach), we have identified a highly-predictive image-computable model of LC neurons. To our knowledge, ours is the first image-computable model of the LC population proposed. An important point is that this encoding model (using a linear mapping) does not identify a one-to-one mapping between model LC units and LC types, as the model is unable to relate the encoded LC neurons to behavior—this is precisely the reason we built the 1-to-1 network. Training the 1-to-1 network both on behavior and neural responses is a worthwhile goal, but care is needed to ensure the neural responses are recorded during natural behavior to achieve as best a match as possible.

**Extended Data Figure 10:**
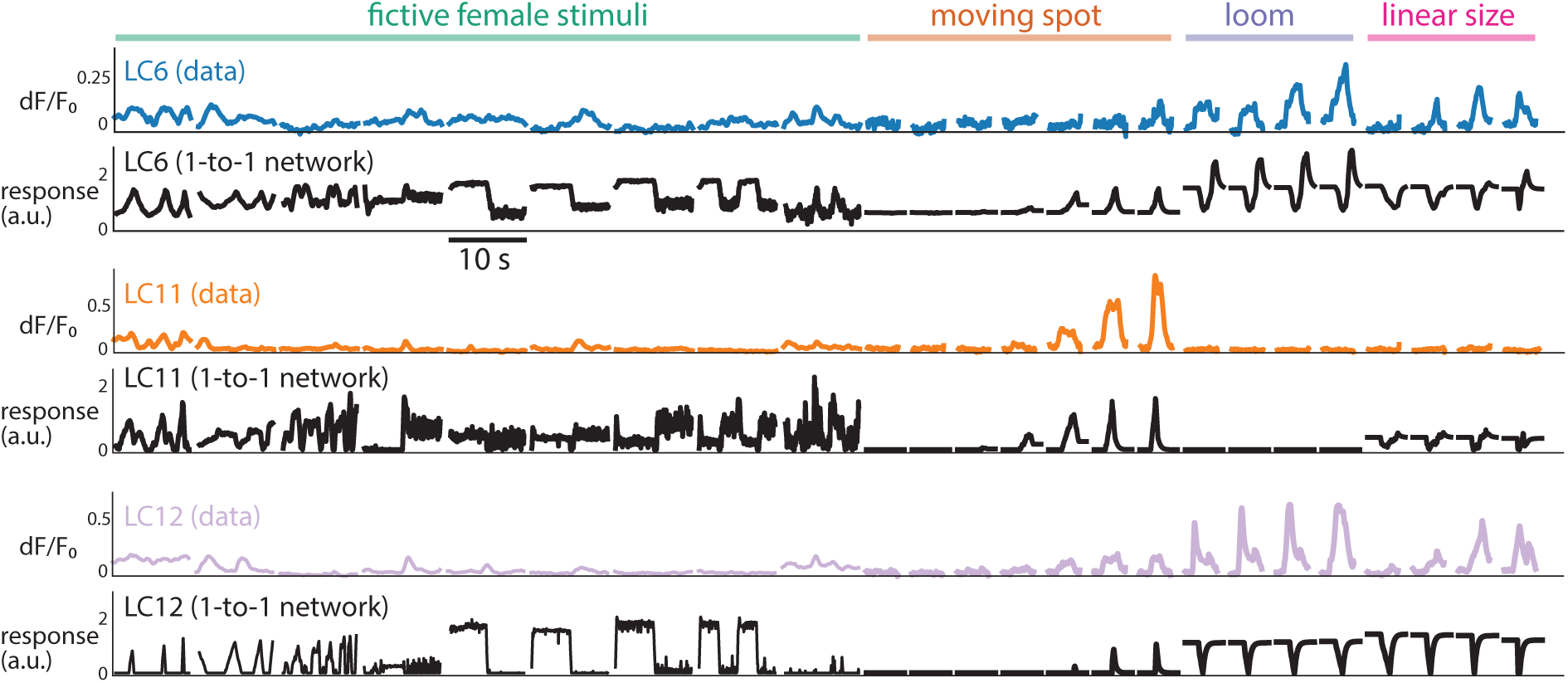
Predicting magnitude of LC responses across both naturalistic and artificial stimuli. One way to assess a viusal neuron’s selectivity is to compare its response magnitudes for different types of stimuli. We wondered whether the relative magnitudes of model LC responses across all stimulus sequences qualitatively matched that of real LC responses. If so, it indicates that the model’s selectivity for certain stimuli matches real LC selectivity. This is different from our quantitive comparisons that normalized model LC responses for each stimulus separately (Fig. 2**d-e** and Ext. Data Fig. 8). We note that *a priori*, we would not expect the 1-to-1 network to predict response magnitude, as downstream weights could re-scale any activity of the model LC units. However, as found when comparing the internal representations of deep neural networks to one another [83], the relative magnitudes of internal units may be an important part of encoding informative representations. Same format as in Fig. 2**f** for the three remaining recorded LC types (LC6, LC11, and LC12). For LC6, the 1-to-1 network correctly predicts a larger response to loom than responses to a moving spot and a spot varying its size linearly (‘linear size’); however, it overestimates the responses to fictive female stimuli. For LC11, the model accurately identifies LC11’s object selectivity (‘moving spot’) and suppression to loom and linear size. Similar to LC6, the 1-to-1 network overestimates LC11’s response magnitudes to the fictive female stimuli. For LC12, the 1-to-1 network has overly large responses to the fictive female stimuli but does predict magnitudes for moving spot, loom, and linear size. The model LC12 responses to loom and linear size appear to be inverted (i.e., flipping model LC12 responses to loom and linear size would better match the real LC12 responses)—this is likely a consequence of the fact that the sign of an LC’s response is unidentifiable for the 1-to-1 network, as one could simply flip the sign of the model LC unit’s response and the readout weights of downstream units. The poor prediction of real LC12 responses is not completely unexpected, as the 1-to-1 network does not perform better at predicting LC12- silenced behavior than DO and noKO networks (Ext. Data Fig. 3); in addition, the KO networks, over 10 runs, are not in strong agreement of LC12’s responses (Ext. Data Fig. 6). Taken together, this suggests that silencing LC12 leads to little changes in behavior compared to control flies, making it difficult to identify LC responses.

**Extended Data Figure 11:**
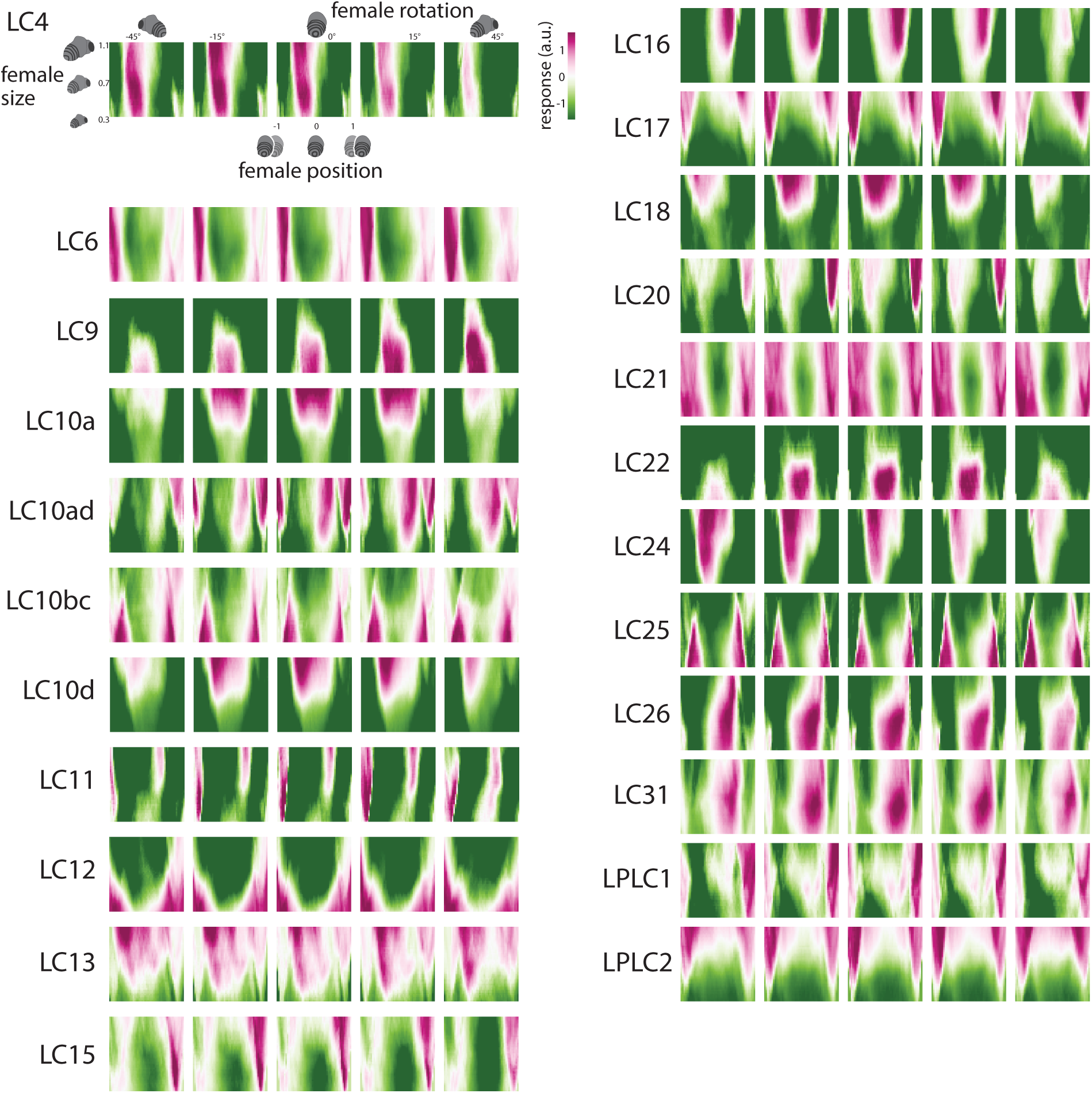
Model LC tuning heat maps. Each “pixel” in the heatmap corresponds to the response of the model LC unit to one input stimulus sequence in which a static fictive female fly has a given size, position, and rotation (i.e., all 10 images of the input sequence were the same, see Methods). We then systematically varied female size, position, and rotation across stimulus sequences (125,000 sequences in total). Same format as in Figure 3c and d but for all model LC units.

**Extended Data Figure 12:**
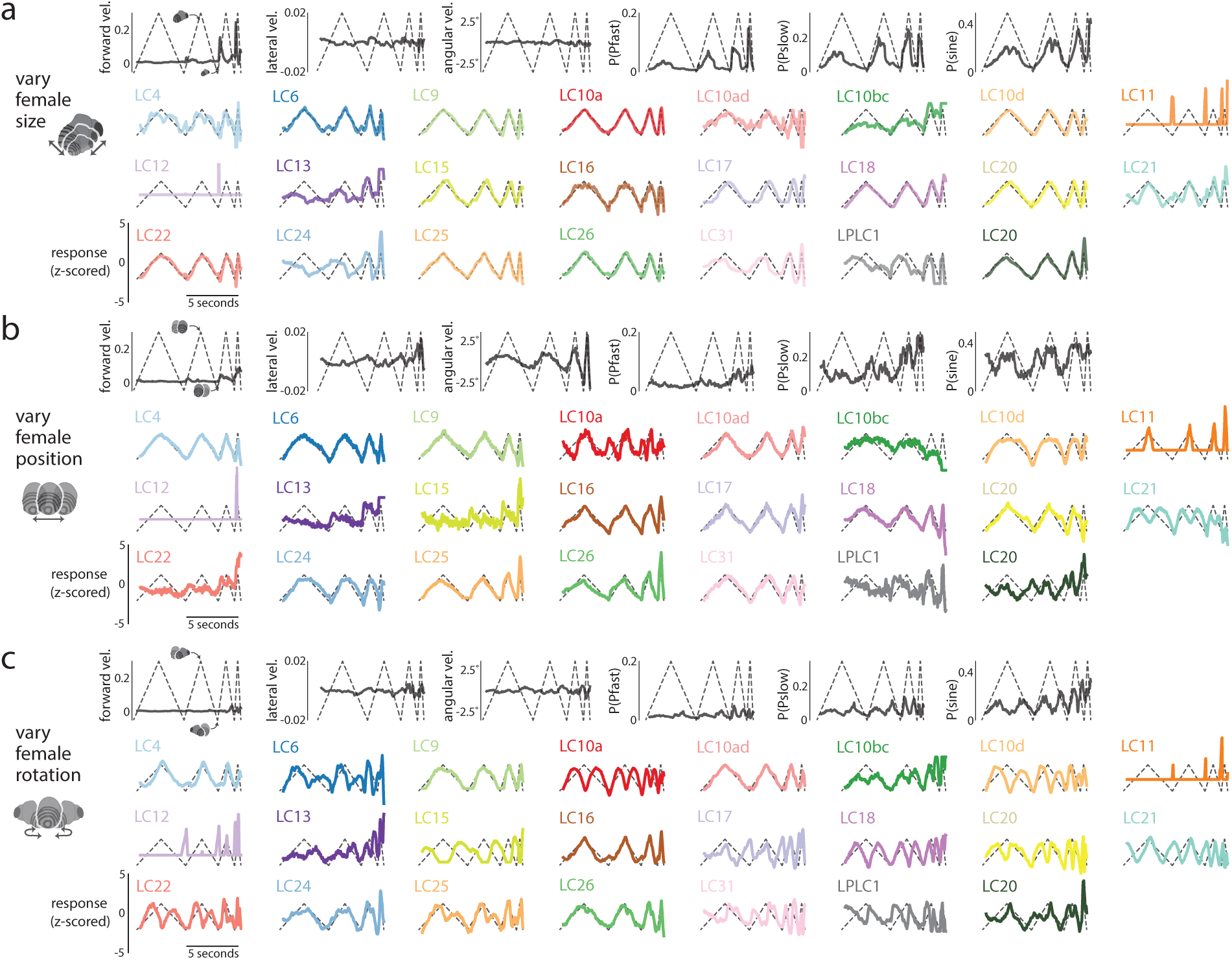
All model LC responses to simple, dynamic stimulus sequences in which only one visual parameter of the fictive female varied. Same dynamic stimulus sequences and format as in Figure 3f; these responses were used to compute the *R*^2^’s in Fig. 3**g**). We also show the 1-to-1 network’s behavioral output for each dynamic stimulus (top rows, black traces). Stimulus sequences include the following (see Methods for exact parameter values): **a.** Varying female size while the female stays in the middle facing away from the male. **b.** Varying female position while the female has a fixed, large size and faces away from the male. **c.** Varying female rotation while the female has a fixed, large size and stays in the middle. Each trace’s sign was flipped to have a positive correlation with the varying visual parameter of the corresponding stimulus sequence.

**Extended Data Figure 13:**
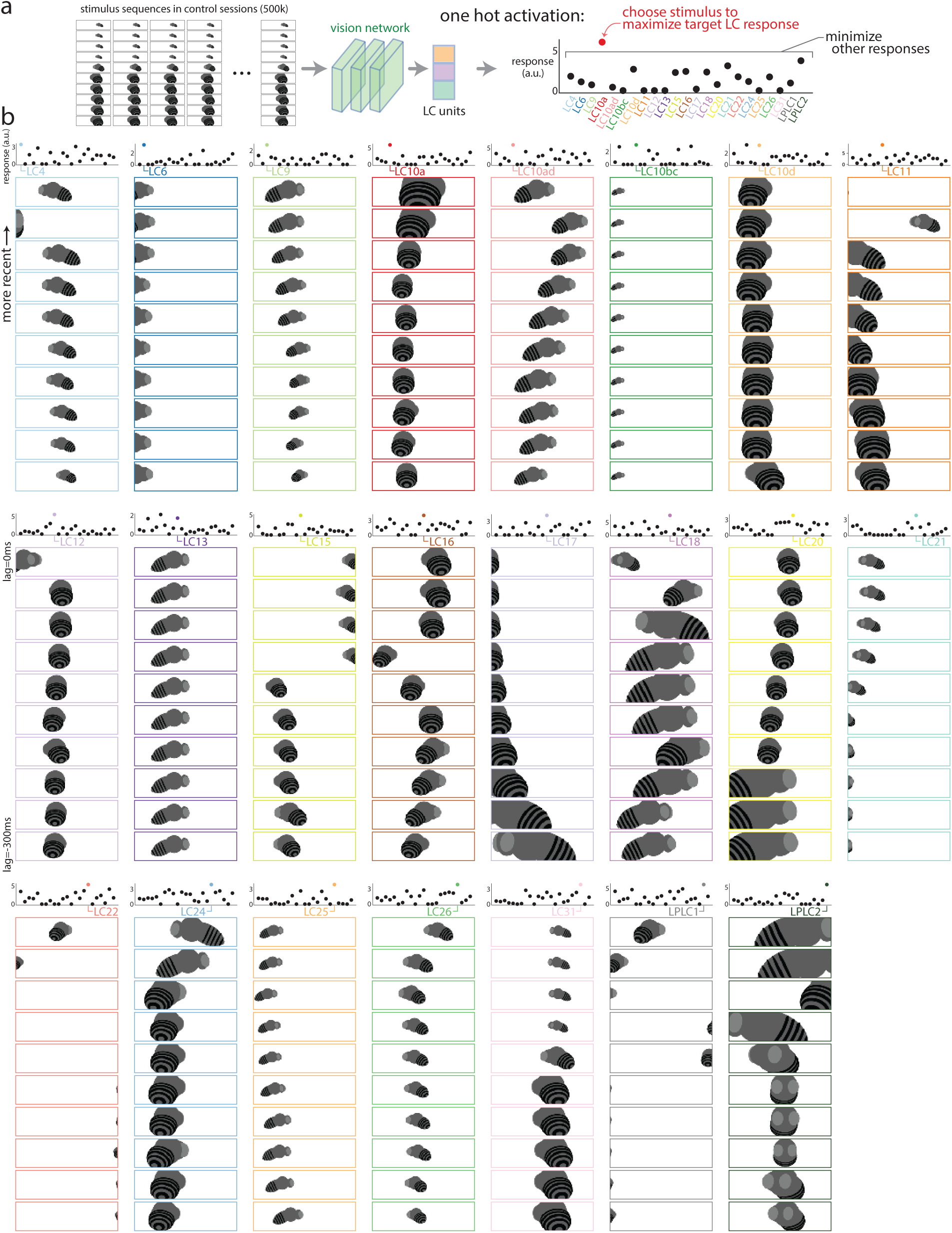
Maximizing visual inputs for each model LC unit. To better understand the differences in stimulus preference across the model LC units, we optimized the visual input history that maximized each model LC unit’s response while minimizing responses of all other model LC units (i.e., a ‘one-hot’ maximizing stimulus). **a.** We considered a large number of candidate stimulus sequences taken from the training data set of control sessions (500,000 stimulus sequences in total). We passed each stimulus sequence as input into the 1-to-1 network, extracting the responses of the model LC units. We chose the stimulus sequence that maximized a chosen model LC unit’s response while minimizing the responses for all other model LC units. We used the following objective function *f_i_*(**x**) for the *i*th chosen model LC unit, adopted from [84]:

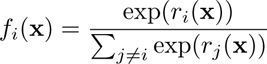 where **x** is the visual input sequence of 10 frames and *r_i_*is the response of the *i*th model LC unit. The objective function *f* (**x**) is maximized for large responses of the *i*th model LC unit and responses as small as possible for all other units. Thus, we optimize stimulus sequences as “one-hot maximizations”. **b.** Maximizing stimulus sequences for each model LC unit with the most recent frame as the top image. One-hot maximization worked for most model LC units (top panel shows responses of all model LC units to that stimulus sequence); a failure (e.g., LC4, top left panel, multiple model LC units have large responses) indicates that this model LC unit shares stimulus preferences with other model LC units. Some stimulus sequences have smooth changes to the fictive female’s parameters, such as LC10a and the increase in female size. However, other maximizing stimulus sequences show large jumps of the fictive female (e.g., LC4, LC12, LC22, etc.); even though these stimulus sequences were chosen from natural courtship, they likely represent outliers that strongly drive responses. These maximizing stimulus sequences represent predictions of the 1-to-1 network that can be tested in future experiments to see if they truly elicit large responses from LC neurons, much like recent work has identified images to drive visual cortical neurons of macaque monkey [84, 85, 86, 87]. Other objective functions, such as maximizing the response variation across time with a longer stimulus sequence, and other constraints, such as restricting how much a fictive female may change between consecutive frames or requiring the fictive female to not remain static, are easily possible with the 1-to-1 network. Our main finding here is that many of the one-hot maximizing stimuli failed to only activate the targeted LC type; this is further evidence that the LC population forms a distributed code.

**Extended Data Figure 14:**
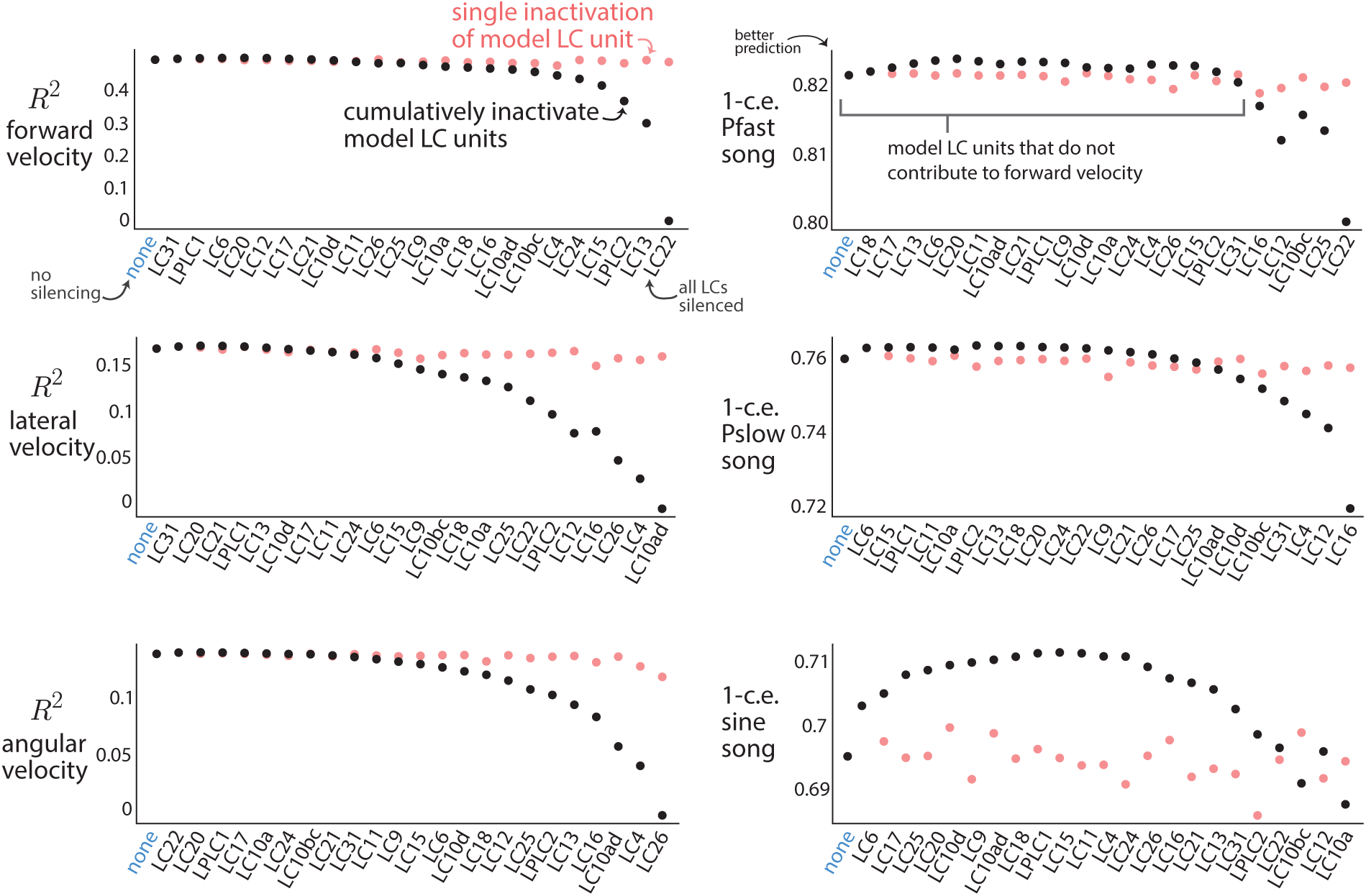
Inactivating model LC units for each of the 6 behavioral output variables during natural courtship. We inactivated each model LC unit separately and re-computed the predicted performance *R*^2^ or 1-*c.e.* on heldout behavioral data from control flies (red dots). Inactivating any single model LC unit did not lead to a large drop in performance, consistent with our experimental findings (Ext. Data. Fig. 1). This indicates that only by inactivating multiple model LC units at the same time will we see a deficit in prediction; in other words, the model LC units form a population code, as multiple model LC units contribute to the same behavioral output. To identify these combinations, we inactivated model LC units in a cumulative, greedy manner (black dots) and observed to what extent the responses for the remaining model LC units predict heldout behavioral data from control flies. Same format as in Figure 4b. For each plot, model LC units on the left contribute the least to the given behavioral output; model LC units on the right contribute the most. We found that when inactivating some model LC units, performance actually *increased* (e.g., LC12 for sine song, bottom right). This is because LC10bc and LC12 used excitation and inhibition to cancel out some of each other’s responses—ablating one decreases performance while ablating both increases performance as both excitatory and inhibitory effects are removed via ablation. The LC neurons themselves need not be either excitatory or inhibitory; readouts by downstream neurons may rely on positively or negatively weighting the LC responses. Performance also increased by removing a number of model LC units for sine song (bottom right, LC6); this is possibly due to overfitting by the 1-to-1 network. By removing “noisy” model LC units that are overfit to the training data, the rest of the model LC units better generalize. Interestingly, the strongest contributor of the model LC units, if inactivated alone, did not lead to a large decrease in performance. For example, LC22 was the strongest contributor for forward velocity but, when inactivated alone, resulted in little decrease to *R*^2^ (red dot above black dot). This is consistent with our finding that silencing LC22 led to little change in mean forward velocity (Ext. Data Fig. 1). This suggests that the model LC units work together as a population code to sculpt behavior: There is no sole contributor to any particular behavior. The red squares of the heatmaps in Figure 4c (which condense the information plotted here) correspond to the differences between the performance value (*R*^2^ or 1-*c.e.*) for each model LC unit and no inactivation (‘none’), divided by the maximum difference (e.g., the difference between the value for the rightmost model LC and the value for ‘none’). To avoid the effects of overfitting, any positive differences (i.e., an increase in prediction performance) were clipped to 1.

**Extended Data Figure 15:**
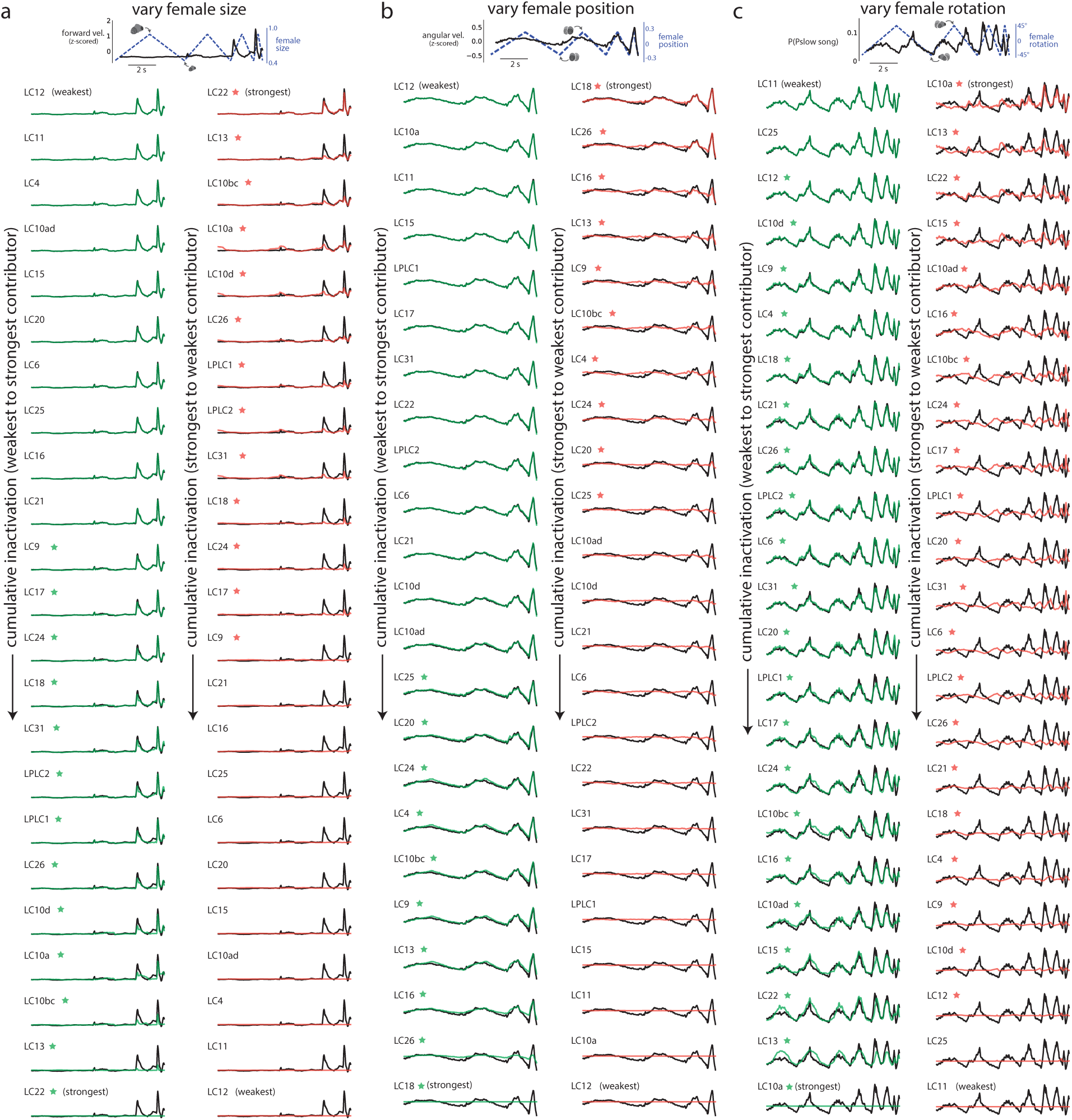
Cumulative inactivation to identify the model LC units necessary and sufficient to produce behavior for the dynamic stimulus sequences. We sought to identify which model LC neurons were necessary and sufficient for behavior in response to simple, dynamic stimulus sequences (e.g., varying female size while holding female position and rotation fixed, see Methods). Here, we show the results for cumulatively inactivating each model LC unit one-by-one; the ordering was determined in a greedy manner by choosing the next model LC unit that, once inactivated, led to the least change in behavior (same process as in Fig. 4**b** and **c**). We used these results to determine which model LC units were necessary and sufficient for these behaviors as reported in Figure 4d and e. **a.** Cumulative inactivation of model LC units for forward velocity (top, black trace) in response to a stimulus sequence in which the fictive female only varies her size (position and rotation remain constant; same dynamic stimulus sequence as in Fig. 4**d**). We observed that the model LC units from LC22 to LC9 (second column, red stars) needed to be inactivated to fully eliminate forward velocity (i.e., necessity); silencing the other model LC units (i.e., those without red stars in the right column) led to no change in behavior (left column, model LC units with green stars are sufficient). **b.** Results for angular velocity (top, black trace) in response to a stimulus sequence in which the fictive female only varies her position (same stimulus sequence as in Fig. 4**e**, left column). We observed that the model LC units from LC18 to LC25 were necessary and sufficient (right column, red stars). **c.** Results for probability of Pslow song (top, black trace) in response to a stimulus sequence in which the fictive female only varies her rotation (same stimulus sequence as in Fig. 4**e**, right column). We observed that the model LC units from LC10a to LC12 were necessary and sufficient (right column, red stars). These results demonstrate that a large number of model LC units contribute to specific behaviors (e.g., 21 out of 23 model LC units for the probability of Pslow song in response to a varying female rotation). They also demonstrate that while some model LC units are strong contributors (e.g., LC22 and LC13 for forward velocity in **a**), many weak contributors also play a role (e.g., in **a**, one must inactivate 13 model LC units—up to LC9—until the behavior is entirely extinguished (i.e., the red trace is flat).

**Extended Data Figure 16:**
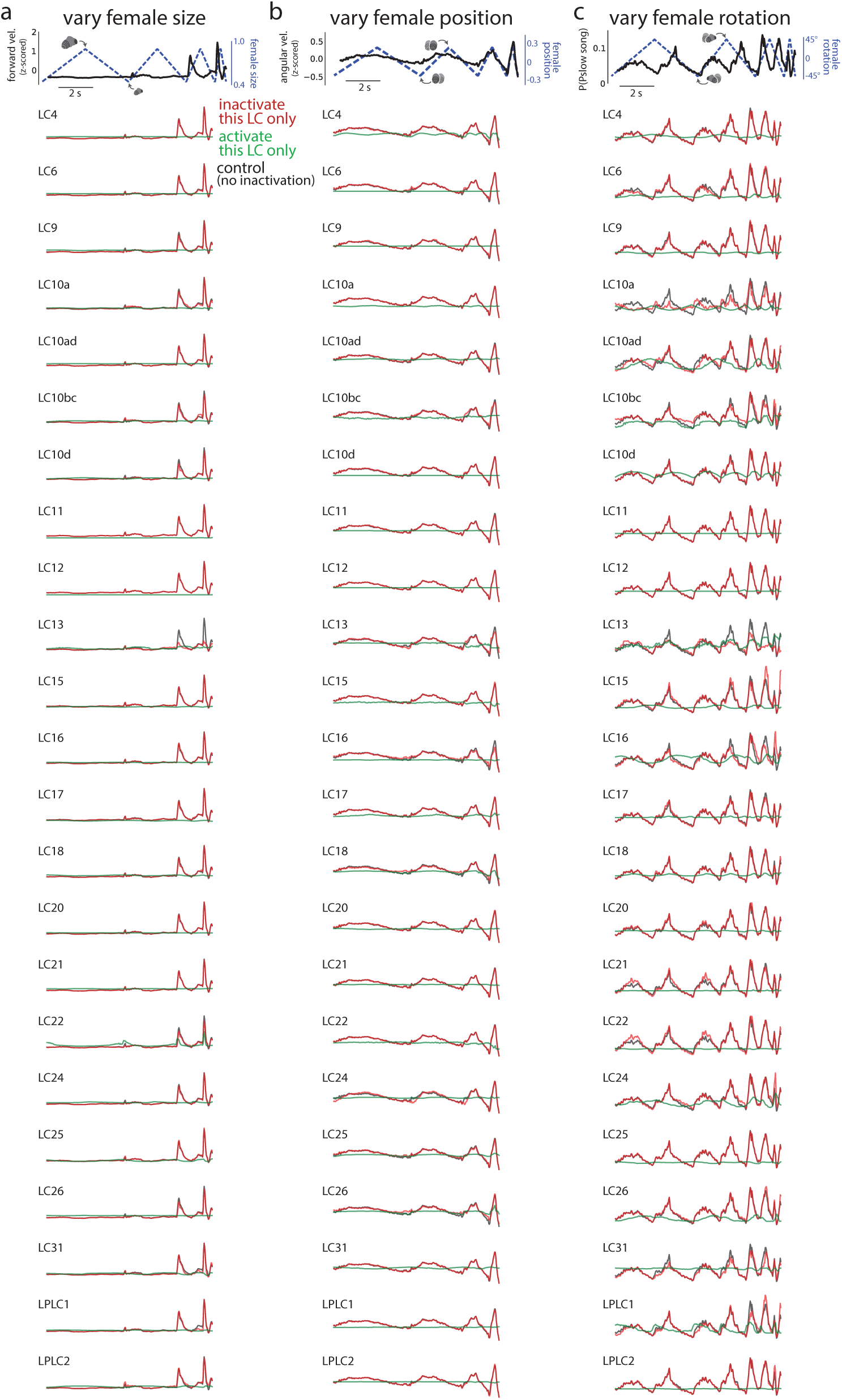
Inactivating and activating individual model LC units. To observe how an individual model LC unit may affect behavior (top row, black traces), we either inactivated only that single model LC unit while keeping all other model LC units active (i.e., knocking out a single model LC unit; red traces) or inactivated all model LC units except that model LC unit (i.e., one hot activation; green traces). We performed this for the three different dynamic stimulus sequences (same sequences as in Fig. 4**d** and **e**). We considered the behavioral outputs of forward velocity (**a**), angular velocity (**b**), and P(Pslow song) (**c**), all of which varied with the corresponding visual parameters (top row, black traces vary with dashed lines). Inactivating any single model LC unit resulted in little to no change in behavior (red traces overlap with black traces), consistent with our single-unit inactivations for naturalistic stimuli (Ext. Data Fig. 14). Likewise, one-hot activation of any single model LC unit produced little variation in behavioral output (most green traces are flat). We observed small but noticeable changes in behavior for model LC13 and LC22 in **a**; LC4, LC13, LC15, LC18, LC22, and LC26 in **b**; and model LC10a, LC10ad, LC10bc, LC10d, LC13, LC15, LC16, LC21, LC22, LC24, LC26, LC31, and LPLC1 in **c**. Overall, these results indicate that no single model LC unit solely contributes to one behavioral output; combinations of model LC units need to be inactivated/activated in order to see appreciable changes in behavioral output.

## Supplemental Tables

**Supplemental Table 1.**
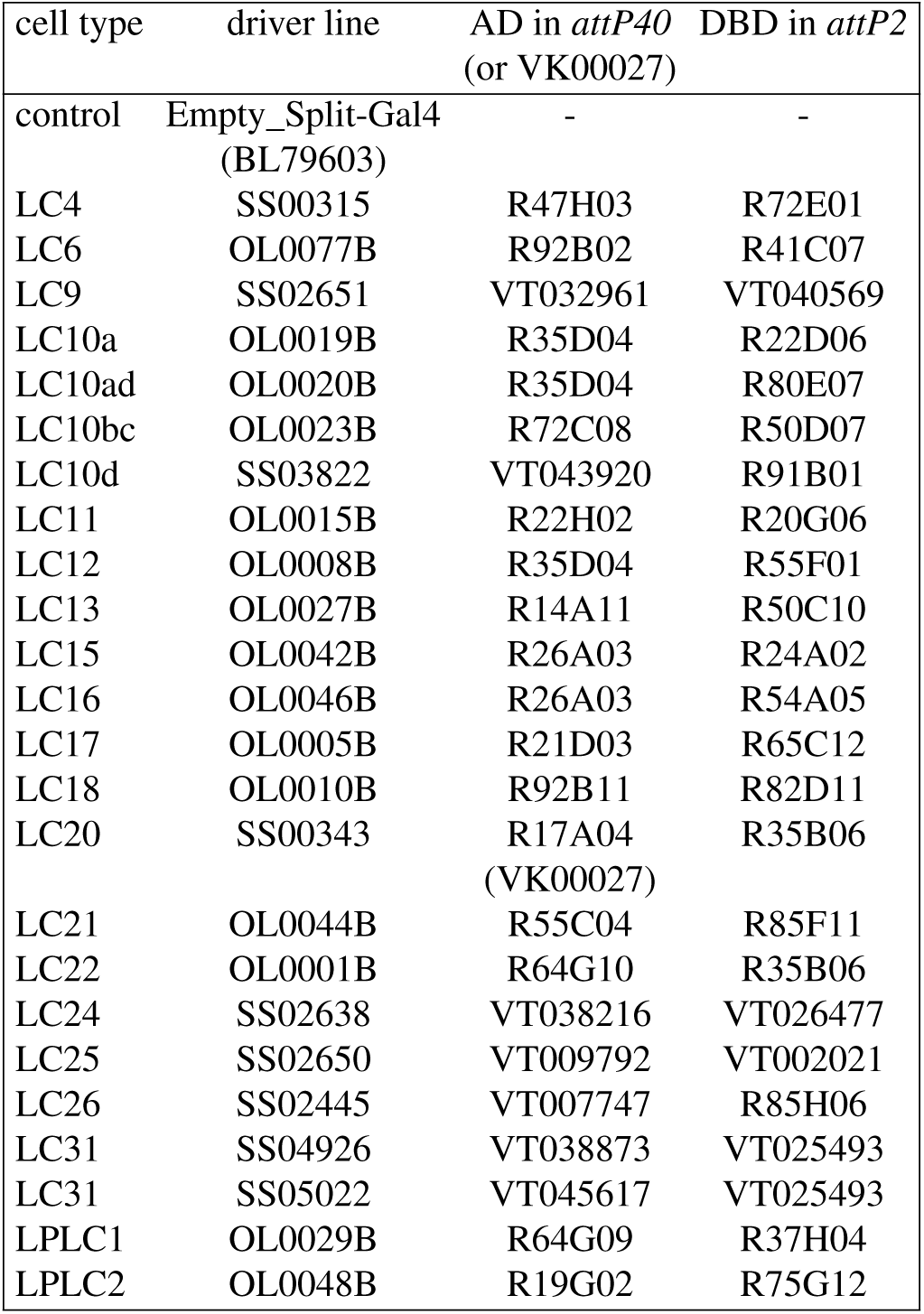
Genetic line information for each of the LC types. ALL LC lines (Wu et al. eLife) and the spGAL4 control line were generously provided by M. Reiser, A. Nern, and G. Rubin. The LC31 lines were generously provided by A. Nern and G. Rubin ahead of publication (Nern et al., *in prep.*). The empty stable split GAL4 line (spGAL4) was used as a control line (BL79603): w[1118]; Py[+t7.7] w[+mC]=p65.AD.UwattP40; Py[+t7.7] w[+mC]=GAL4.DBD.UwattP2 [64]. Males were paired with females of the following genotype: PIBL (pheromone-insensitive and blind) *w*+*/w*+*; GMR-hid/+; orco/ orco* (recombined into the CS background, see below)

In Extended Data Figure 1, the CS control data was from a previous study [9], listed as Wildtype 2, Canton S laboratory strain. Blind males (BL) were of the following genotype and also recombined into the Canton S background: w+/w+; GMR-hid/GMR-hid, also from the same previous study [9].

More information can be found at the following websites:

- https://splitgal4.janelia.org/precomputed/LC%20Paper.html
- https://flybase.org/reports/FBst0028996.html
- https://flybase.org/reports/FBtp0001264.html

**Supplemental Table 2.** We crossed each of the 5 LC split lines to GCaMP6f (+/+; UAS GCaMP6f/CyO; +, provided by the Clandinin Lab) and imaged from the male progeny.

| cell type | driver line | AD in *attP40* | DBD in *attP2* |
| --- | --- | --- | --- |
| LC6 | OL0070B | R91B01 | R22A07 |
| LC11 | OL0015B | R22H02 | R20G06 |
| LC12 | OL0008B | R35D04 | R55F01 |
| LC15 | OL0042B | R26A03 | R24A02 |
| LC17 | OL0005B | R21D03 | R65C12 |

